# Production of the anticancer drug intermediate strictosidinic acid in engineered yeast

**DOI:** 10.1101/2025.10.29.685393

**Authors:** Benedikt Seligmann, Shenyu Liu, Mai Huynh, Thu-Thuy T. Dang, Jakob Franke

## Abstract

Strictosidinic acid is a key intermediate in the biosynthetic pathway of camptothecin, a plant alkaloid that serves as a precursor for semisynthetic anticancer drugs. At the moment, camptothecin is mainly sourced from trees, causing limited supply and high costs. Improving access to strictosidinic acid would help to elucidate yet unknown biosynthetic steps and in the long term enable sustainable production of camptothecin in heterologous hosts. While structurally similar to the common monoterpene indole alkaloid precursor strictosidine, strictosidinic acid has not been the target of metabolic engineering efforts before. Here, we present a strategy to produce strictosidinic acid from glucose and tryptophan in engineered yeast. First, we create a basic strain that generates 75 mg/L strictosidine. We further optimise this strain by introducing a membrane steroid binding protein and a second copy of the farnesyl pyrophosphate synthase mutant gene *ERG20*^*WW*^, boosting strictosidine levels by 5.5-fold to 398 mg/L. At these higher titres, a previously overlooked shunt product, (2*E*,6*E*)-2,6-dimethylocta-2,6-dienedioic acid (DOA), was identified that diverts flux from the pathway. Lastly, we reprogrammed our strictosidine strain to strictosidinic acid production by four genomic modifications. Final fed-batch cultivation in shake flasks resulted in 843 mg/L strictosidine or 548 mg/L strictosidinic acid, respectively, after 168 hours. Taken together, our work now grants access to strictosidinic acid by metabolic engineering, while revealing strategies to further enhance the production of strictosidine and related monoterpene indole alkaloids. These findings will help to produce plant alkaloids in microbial cell factories in the future at scale.

## 1. Introduction

Several clinically used anticancer drugs are derived from the plant alkaloid camptothecin (**1**) (Khaiwa et al., 2021; Martino et al., 2017) (Fig. 1). For example, topotecan (**2**) is used to treat ovarian cancer (Khaiwa et al., 2021) and irinotecan is employed against colorectal cancer (Fujita et al., 2015). Even to date, new derivatives of camptothecin (**1**) are in drug development (Khaiwa et al., 2021; X. Wang et al., 2023). Many clinically relevant camptothecin (**1**) derivatives are produced by semisynthetic conversion of camptothecin (**1**) isolated from trees such as *Camptotheca acuminata* (Goel and Jain, 2024). Therefore, the natural supply of this alkaloid currently represents a critical factor for our ability to manufacture camptothecin-based anticancer drugs. As such, there is a high demand to develop innovative approaches for the biotechnological production of camptothecin (**1**) in microbial hosts.

**Fig. 1.**
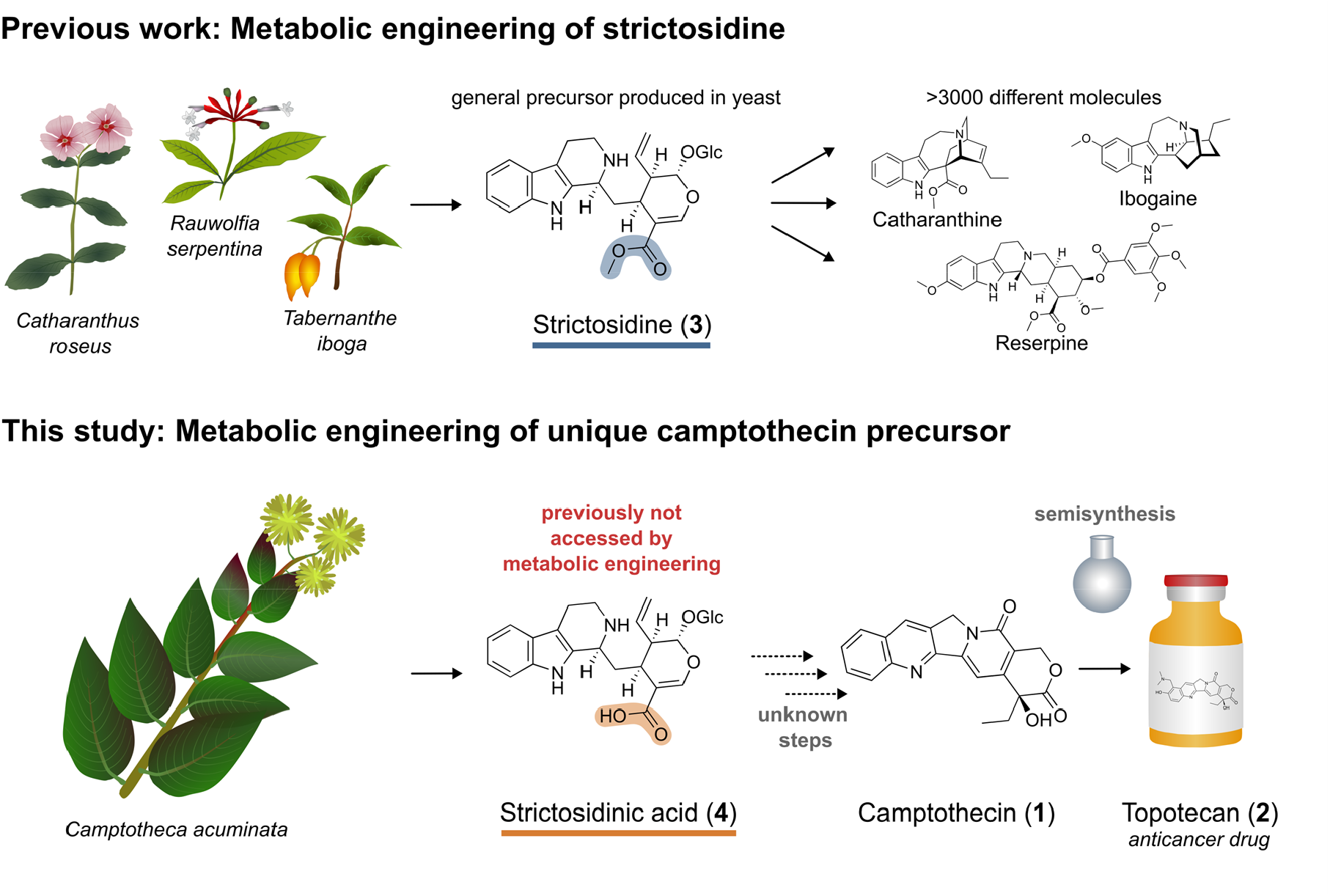
Strictosidinic acid (4) is the precursor of clinically used anticancer drugs derived from the plant alkaloid camptothecin (1). In contrast to the more common monoterpene indole alkaloid precursor strictosidine (**3**), strictosidinic acid (**4**) has not yet been accessed by metabolic engineering.

Biosynthetically, camptothecin (**1**) is a monoterpene indole alkaloid with a unique twist (Fig. 1). Whereas all monoterpene indole alkaloids are derived from the central biosynthetic precursor strictosidine (**3**) (O”Connor and Maresh, 2006), camptothecin (**1**) represents an exception from this paradigm. Current evidence suggests that strictosidinic acid (**4**) rather than strictosidine (**3**) is the precursor of camptothecin (**1**) in the plant *C. acuminata* (Sadre et al., 2016).

The common monoterpene indole alkaloid precursor strictosidine (**3**) has been a very popular target compound for the metabolic engineering community. In 2015, production of strictosidine (**3**) in engineered yeast was described for the first time (Brown et al., 2015). Since then, several further studies have improved the capacity of yeast to produce strictosidine (**3**) and strictosidine-derived alkaloids (Bradley et al., 2023; Dror et al., 2024; Holtz et al., 2024; Liu et al., 2022; Misa et al., 2022; Zhang et al., 2022). In addition, heterologous production of strictosidine (**3**) has also been accomplished in *Pichia pastoris* (Gao et al., 2023) and the model plant *Nicotiana benthamiana* (Dudley et al., 2022). The highest reported titres for *de novo* strictosidine (**3**) production in yeast were 91 mg/L in fed-batch microbioreactors (Holtz et al., 2024) and ∼30-40 mg/L in culture tubes (Dror et al., 2024).

In contrast to strictosidine (**3**), the camptothecin precursor strictosidinic acid (**4**) has not been a target of metabolic engineering efforts so far. The biosynthetic pathway of camptothecin (**1**) downstream of strictosidinic acid (**4**) is not yet elucidated (Kang et al., 2021; Pu et al., 2019), with a few scattered exceptions (Chen et al., 2024; Nguyen et al., 2021; Pu et al., 2024, 2023; Zhang et al., 2024). One of the reasons is the limited commercial availability of camptothecin (**1**) pathway intermediates such as strictosidinic acid (**4**) which hampers efforts to elucidate the biochemical pathway. Developing a yeast platform capable of producing strictosidinic acid (**4**) would not only represent an excellent starting point for the biotechnological production of camptothecin (**1**) in the future, but would also facilitate pathway elucidation efforts by improving access to rare intermediates of camptothecin (**1**) biosynthesis (Kwan et al., 2023). Here, we engineer yeast (*Saccharomyces cerevisiae*) to produce strictosidinic acid (**4**) from glucose and tryptophan (**5**). First, we set up a basic strictosidine (**3**) platform strain containing 25 genetic modifications compared to the background strain. Next, we optimized strictosidine (**3**) production by screening twelve membrane steroid binding protein homologues to enhance cytochrome P450 monooxygenase activity, and by testing farnesyl pyrophosphate synthase mutant combinations to favour monoterpene rather than sesquiterpene production. The high product titres allowed us to identify a previously overlooked geraniol overoxidation shunt product (**6**) that partially traps carbon flux towards strictosidine (**3**) and strictosidinic acid (**4**). Lastly, we show that removal of a methyltransferase and a cytochrome P450 monooxygenase gene and addition of two other cytochrome P450 monooxygenase genes allows efficient reprogramming from strictosidine (**3**) to strictosidinic acid (**4**) production. Final yeast strains produced 843 mg/L strictosidine (**3**) or 548 mg/L strictosidinic acid (**4**), respectively, after 168 hours in a fed-batch system. Our study demonstrates general strategies to further improve the production of monoterpene indole alkaloids in yeast; more importantly, it sets the starting point for the sustainable production of camptothecin (**1**) in a microbial host.

## 2. Material and Methods

### 2.1. Construction of plasmids, integration fragments, and yeast strains

Genes, promoters, and terminators used in this study were either cloned from yeast gDNA, plant cDNA or ordered as synthetic genes. For amplification of plant genes from *Ailanthus altissima, Arabidopsis thaliana, Catharanthus roseus, Citrus sinensis*, and *Nepeta cataria*, RNA from leaves was isolated using the GeneJET Plant RNA Purification Mini Kit (Thermo Scientific) and converted into cDNA using the SuperScript™ IV VILO™ Master Mix (Invitrogen). *CYP72A565, CYP72A610*, and *MSBP* genes from *C. roseus* and *C. acuminata* as well as *A. thaliana MSBP2*-*4* were obtained as synthetic genes from Twist Bioscience.

All integration plasmids for use with the EasyClone-MarkerFree (Jessop-Fabre et al., 2016) or Expanded EasyClone-MarkerFree (Babaei et al., 2021) vector set were assembled by USER cloning (Bitinaite et al., 2007). Point mutations were generated by site-directed mutagenesis of genes with In-

Fusion cloning (Takara Bio). Custom guide RNA (gRNA) plasmids were generated using In-Fusion mutagenesis (Takara Bio) or KLD mutagenesis (New England Biolabs) with gRNA plasmid pCfB3045 (Jessop-Fabre et al., 2016) as template.

Afterwards, *E. coli* DH5α competent cells (New England Biolabs) were transformed with assembled constructs, plated on lysogeny broth (LB) agar plates containing 100 µg/mL ampicillin, and grown overnight at 37 °C. Colonies were picked, in case of cloning reactions screened by colony PCR, followed by growing overnight cultures in 5 mL LB medium with 100 µg/mL ampicillin at 37 °C and shaking at 210 rpm. Plasmid DNA was isolated from overnight cultures using Wizard® Plus SV Minipreps (Promega) and verified by Sanger sequencing (Microsynth or Eurofins).

All yeast strains used in this study are based on ST7574 (Euroscarf). For integration of genes into yeast chromosomes, EasyClone-MarkerFree plasmids were linearised using NotI-FD (Thermo Scientific) or NotI-HF (New England Biolabs) to generate linear integration fragments. Integration fragments for custom deletions were generated by PCR or using primer pairs with overlapping complementary 3’ ends. Yeast strains were transformed with linear integration fragments together with the corresponding gRNA plasmid either from the EasyClone-MarkerFree vector set (Jessop-Fabre et al., 2016), the Expanded EasyClone-MarkerFree vector set (Babaei et al., 2021), or with a custom gRNA plasmid. Yeast transformation was done using lithium acetate/single-stranded carrier DNA/PEG protocol (Gietz and Schiestl, 2007). Successful integration or deletion of genes was confirmed by yeast colony PCR or Sanger sequencing.

All yeast strains from this study are listed in Supplementary Table 1. Genes are provided in Supplementary Table 2, as well as primers (Supplementary Table 3 – Supplementary Table 8) and gRNA sequences (Supplementary Table 9). The relationships of yeast strains from this study are shown in Supplementary Fig. 1.

### 2.2. Cultivation of yeast strains

For general propagation and transformation, yeast strains were grown at 30 °C in standard YPD medium (yeast extract 1% (w/v), peptone 2% (w/v), glucose 1% (w/v)).

For screening of yeast cultures in small scale, 4 mL primary yeast cultures were inoculated in YPD medium in 50 mL centrifuge tubes containing 200 µg/mL geneticin (G-418). After growth for 16-18 hours at 30 °C and 210 rpm shaking speed, the primary cultures were used to inoculate 4 mL secondary cultures in YPD medium with 2 mM tryptophan in 50 mL centrifuge tubes without antibiotics to OD600 0.1. Secondary culture tubes contained a sterile pipette tip to improve mixing and prevent pelleting, and were covered with a loose aluminium foil lid for improved gas exchange. The secondary cultures were grown at 30 °C and 210 rpm shaking speed. After 48 hours, 2% ethanol was fed; in case of substrate feeding experiments, 2% ethanol with 50 mM of intermediates (final concentration 1 mM) was used instead. The cultivation was stopped after a total of 120 hours of incubation.

For fed-batch cultivations in shake flasks, 4 mL primary yeast cultures were inoculated in YPD medium in 50 mL centrifuge tubes containing 200 µg/mL geneticin. After growth for 16-18 hours at 30 °C and 210 rpm shaking speed, the primary cultures were used to inoculate 20 mL seed cultures in concentrated YPD medium (yeast extract 2% (w/v), peptone 4% (w/v), glucose 1% (w/v)) with 2 mM tryptophan in 250 mL shake flasks without antibiotics to OD600 of 2. After growth for 4 hours at 30 °C and 210 rpm shaking speed, the seed cultures were used to inoculate 50 mL concentrated YPD medium in 250 mL baffled flasks and membrane lids for improved aeration. At 30 hours after inoculation, 2% ethanol and 2% concentrated YP media (yeast extract 2% (w/v), peptone 4% (w/v)) with 25 mM tryptophan (final concentration 0.5 mM) was fed to the cultures. Starting at 48 hours after inoculation, 1% ethanol and 1% concentrated YP media (yeast extract 2% (w/v), peptone 4% (w/v) with 25 mM tryptophan (final concentration 0.25 mM) were fed to the cultures every 24 hours. The total cultivation time was 168 hours.

### 2.3. Metabolites and chemicals

Metabolites used in this study either as reference compounds, as standards for quantification, or for feeding experiments are listed in Supplementary Table 10.

Chemicals not mentioned specifically were purchased from Fisher Scientific, Carl Roth, or Sigma Aldrich. Silica gel 60M (40-60 µM) and TLC plates (F254) were purchased from Macherey-Nagel. Catnip oil was purchased from Piping Rock (USA).

### 2.4. General chemical methods

Analytical and semipreparative LC-MS analyses were performed on an Agilent Infinity II 1260 system consisting of a G7167A autosampler, G7116A column thermostat, G7111B quaternary pump, G7110B make-up pump, G7115A diode array detector, G1364F fraction collector, and G6125B single quadrupole mass spectrometer equipped with an ESI source (positive mode, 4000 V, 12 L/min drying gas, 350 °C gas temperature). Formic acid (0.1% v/v) was added to the mobile phase as indicated. The columns and gradients used are described in the corresponding method sections.

Automated flash chromatography was performed on a Biotage Isolera One with the stationary phases, solvents and gradients described in the corresponding method sections.

NMR spectra were recorded using Bruker Ultrashield 400, Ultrashield 500 or Ascend 600 MHz spectrometers operating at 400, 500 and 600 MHz for ^1^H NMR and at 100, 126 and 151 MHz for ^13^C NMR. Chloroform-d, methanol-d4, or DMSO-d6 were used as solvents. Chemical shifts were referenced relative to the residual solvent signals (chloroform-d: δH = 7.26 ppm, δC = 77.16 ppm; methanol-d4: δH = 3.31 ppm, δC = 49.00 ppm; DMSO-d6: δH = 2.50 ppm, δC = 39.52 ppm) and expressed in δ values (ppm), with coupling constants reported in Hz. Analysis was conducted with TopSpin (Version 4.4.1) or MestReNova (Version 14.2).

HRMS measurements were carried out on a Waters Acquity UPLC coupled to a Waters QTof Premier mass spectrometer.

Synthetic methods are described in Supplementary methods.

### 2.5. Metabolite analysis

For the extraction of metabolites, 250 µL of the yeast cultures were mixed with 1 mL methanol. The suspension was homogenized using a bead beating homogenizer (FastPrep-24Tm 5G) and glass beads (⌀ = 0.5 mm, shaking at 6.0 m/s for 40 s with two cycles). After centrifugation at 17,000 × *g*, the supernatant was filtered and analysed by LC-MS. Samples were separated on a C18 column (Poroshell 120 EC-C18, 100 × 4.6 mm, 2.7 μm). The column temperature was set to 50 °C. As mobile phase, solvent A (water with 0.1% (v/v) formic acid) and solvent B (acetonitrile with 0.1% (v/v) formic acid) were used. The following gradient at a flow rate of 1.0 mL/min was used: 0-1 min, 5% B; 1-11 min, 5-55% B; 11-16 min, 55-100% B; 16-18 min, 100% B; 18-18.5 min, 100-5% B. Mass spectra were obtained in scan mode in a range of *m*/*z* 100-600 at a fragmentor voltage of 100 V. Quantification of metabolites was performed with calibration curves generated from reference compounds. Sources of metabolites used are listed in Supplementary Table 10.

### 2.6. Isolation and identification of (2E,6E)-2,6-dimethylocta-2,6-dienedioic acid (DOA) (6)

A 50 mL culture of yeast strain BSY101 was centrifuged and the sterile filtrated supernatant was concentrated under reduced pressure, suspended in MeOH (15 mL) and filtered. The remaining solid residue was again suspended in MeOH and filtered; this was repeated three times (3 × 10 mL). The filtrates were collected, combined and purified by flash chromatography (SNAP Ultra 25 g; A: DCM, B: MeOH; 2 CV 5% B, 10 CV 5-80% B, 3 CV 80% B, 1 CV 80-100% B, 2 CV 100% B) to give crude **6**. After further purification by semipreparative LC-MS (Kinetex 5 µm C18 100 Å 250 x 10 mm, A: H2O + 0.1 % v/v formic acid, B: MeCN+ 0.1 % v/v formic acid; flowrate: 5 mL/min; temperature: 40 °C; gradient: 0-10 min 20-30% B, 10-10.5 min 30-95% B, 10.5-13 min 95% B, 13-13.5 min 95-20% B, 13.5-15 min 20% B), **6** was obtained as a pale-yellowish solid (2.3 mg).

HR-ESI-MS: found [M+Na]^+^ 221.0783 (calcd. for C10H14O4Na^+^ 221.0784).

NMR spectra and data are shown in Supplementary Fig. 2-6 and Supplementary Table 11.

### 2.7. Bioinformatic search for MSBP homologues

To identify *MSBP* homologues, sequence datasets from *C. acuminata* (D. Zhao et al., 2017), *C. roseus* (Li et al., 2023), and *A. altissima* (Chuang et al., 2022), were used to perform custom BLASTN searches with Geneious Prime® 2024.0.5.

## 3. Results

### 3.1. Generation of a basic strictosidine platform strain

Several studies have already described the production of strictosidine (**3**) and strictosidine-derived alkaloids in yeast and related systems (Bradley et al., 2023; Brown et al., 2015; Dror et al., 2024; Gao et al., 2023; Holtz et al., 2024; Liu et al., 2022; Misa et al., 2022; Zhang et al., 2022). Considering the large structural similarity between strictosidine (**3**) and strictosidinic acid (**4**), we envisioned that reprogramming a strictosidine (**3**)-producing strain to produce strictosidinic acid (**4**) should be feasible. Therefore, our strategy was to first set up and optimise a system for strictosidine (**3**) production rather than directly aiming to produce our target compound strictosidinic acid (**4**). Accordingly, we constructed a first basic strictosidine (**3**) strain (strain BSY84) using the EasyClone-MarkerFree system (Babaei et al., 2021; Jessop-Fabre et al., 2016). Combining previous strategies from literature (Bradley et al., 2023; Brown et al., 2015; Dror et al., 2024; Gao et al., 2023; Holtz et al., 2024; Liu et al., 2022; Misa et al., 2022; Zhang et al., 2022), our yeast strain contained a total of 25 genetic modifications, consisting of overexpression of 15 plant genes, one mutant yeast gene, 3 native yeast genes, one promoter switch and 5 endogenous gene deletions (Fig. 2). Starting from the mevalonate (MVA) pathway building blocks isopentenyl pyrophosphate (IPP) (**7**) and dimethylallyl pyrophosphate (DMAPP) (**8**), geranyl pyrophosphate (GPP) (**9**) was produced *via* yeast Erg20. Flux to monoterpenoids rather than to farnesyl pyrophosphate (FPP) (**10**) was reduced by using mutated yeast Erg20 variants and a promoter switch (Kong et al., 2023) as further described in section 3.2. GPP (**9**) is then converted to geraniol (**11**) by truncated geraniol synthase, which lacks a plastid transit peptide to enforce cytosolic localisation. Several oxidation steps in the so called iridoid pathway *via* 8-hydroxygeraniol (**12**), 8-oxogeranial (**13**), and nepetalactol (**14**) lead to 7-deoxyloganetic acid (**15**). After glycosylation to 7-deoxyloganic acid (**16**), this intermediate is oxidised by 7-deoxyloganic acid hydroxylase (7DLH) to loganic acid (**17**), methylated by loganic acid methyltransferase (LAMT) to loganin (**18**), and further oxidised by secologanin synthase (SLS) to secologanin (**19**). In parallel, tryptophan (**5**) is decarboxylated to tryptamine (**20**). As the last step, secologanin (**19**) and tryptamine (**20**) are merged by strictosidine synthase (STR) to generate strictosidine (**3**). NADPH availability was further increased using ZWF1 (Brown et al., 2015; H. Wang et al., 2023).

**Fig. 2.**
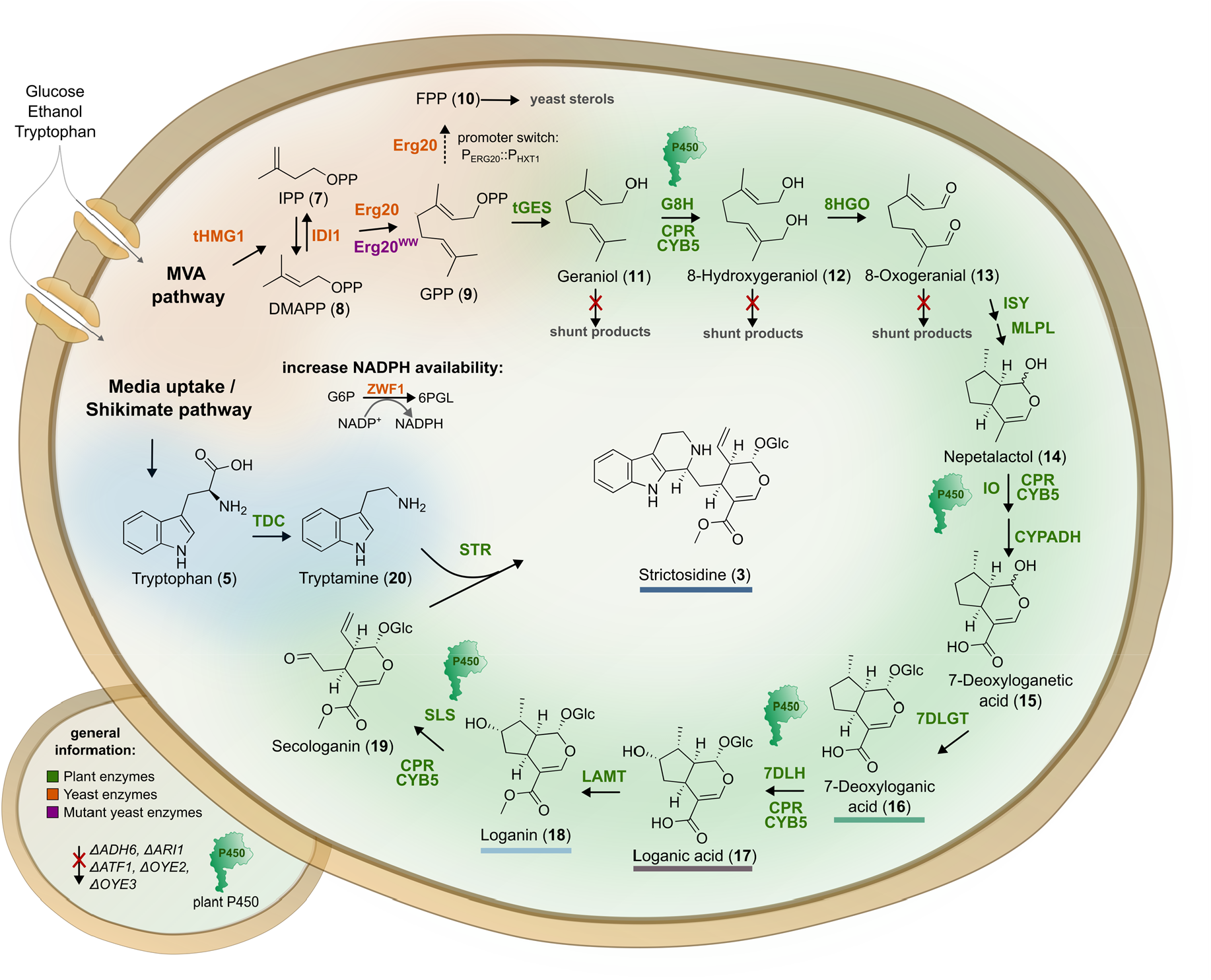
Engineering strategy in yeast to generate a platform strain for the production of strictosidine (3). The resulting strain BSY84 produced strictosidine (**3**) at a titre of 75 mg/L. Plant enzymes are shown in green, yeast enzymes in orange, and mutant yeast enzymes in purple. The four P450s in the pathway are highlighted by small green protein icons. MVA, mevalonate; tHMG1, truncated hydroxymethylglutaryl-CoA reductase 1; IPP, isopentenyl pyrophosphate; DMAPP, dimethylallyl pyrophosphate; IDI1, isopentenyl diphosphate isomerase 1; Erg20^WW^, double mutant of Erg20 (F96W, N127W); FPP, farnesyl pyrophosphate; GPP, geranyl pyrophosphate; tGES, truncated geraniol synthase; G8H, geraniol 8-hydroxylase; CPR, cytochrome P450 reductase; CYB5, cytochrome b5; 8HGO, 8-hydroxygeraniol oxidoreductase; ISY, iridoid synthase; MLPL, major latex protein-like protein; IO, iridoid oxidase; CYPADH, alcohol dehydrogenase 2; 7DLGT, 7-deoxyloganetic acid glucosyltransferase; 7DLH, 7-deoxyloganic acid hydroxylase; LAMT, loganic acid methyltransferase; SLS, secologanin synthase; TDC, tryptophan decarboxylase; STR, strictosidine synthase; ADH6, alcohol dehydrogenase 6; ARI1, NADPH-dependent aldehyde reductase, ATF1, alcohol acetyltransferase 1; OYE2, old yellow enzyme 2; OYE3, old yellow enzyme 3.

Strictosidine (**3**) was identified by LC-MS analysis in comparison to the retention time and mass spectrum of a standard (Supplementary Fig. 7). After 120 hours cultivation, our platform strain BSY84 produced strictosidine (**3**) at a level of 75 mg/L, which was close to the highest reported strictosidine (**3**) titres of 91 mg/L in fed-batch microbioreactors (Holtz et al., 2024), and outperformed the strictosidine (**3**) yield of ∼30-40 mg/L reached in culture tubes (Dror et al., 2024).

### 3.2. Improving strictosidine production

After successfully establishing strictosidine (**3**) production in yeast, we next aimed at improving this system further. Cytochrome P450 monooxygenases (P450s) from plants often exhibit limited activity in non-plant hosts and thereby cause metabolic bottlenecks (Jiang et al., 2021). In plants, a membrane steroid binding protein (MSBP) was identified in lignin biosynthesis that fosters P450 activity, possibly by stabilising P450s or by improving their interaction with electron transfer partner proteins (Gou et al., 2018) (Fig. 3A). This beneficial activity of MSBP has been harnessed to increase triterpene production in yeast (Liu et al., 2024; Yang et al., 2023). As the iridoid pathway leading to strictosidine (**3**) includes four P450s (Fig. 2), we hypothesised that MSBP might also improve strictosidine (**3**) production. So far, it remains unclear whether MSBP homologues from different plants are specifically tailored to certain biochemical pathways or plant systems, or function more universally to improve P450 activity. To investigate this point, we selected a panel of twelve MSBP homologues for functional evaluation (Fig. 3B, Supplementary Table 2). This included two homologues from the camptothecin (**1**) producing plant *C. acuminata* and two from the model monoterpene indole alkaloid producer *C. roseus*. For comparison with non-monoterpene indole alkaloid producers and more distantly related organisms, we additionally selected three homologues from *Citrus sinensis* and *Ailanthus altissima*, plants known for production of triterpenoids and to a lesser extent acridone and β-carboline alkaloids (Fernandes da Silva et al., 2022), four from *A. thaliana*, mostly known as a glucosinolate producer (Sønderby et al., 2010), and *S. cerevisiae* Dap1, the closest homologue of *At*MSBP1 in the host organism with 31.6% amino acid sequence identity (González et al., 2022, 2022). With the exception of AtMSBP2 and *At*MSBP4, all MSBP homologues including yeast *Sc*Dap1p led to a strong, 2.0 – 4.0-fold increase in strictosidine (**3**) levels (Fig. 3B, Supplementary Fig. 8). The highest titres were obtained for *C. acuminata* MSBP1, resulting in 291 mg/L strictosidine (**3**) (strain BSY212).

**Fig. 3.**
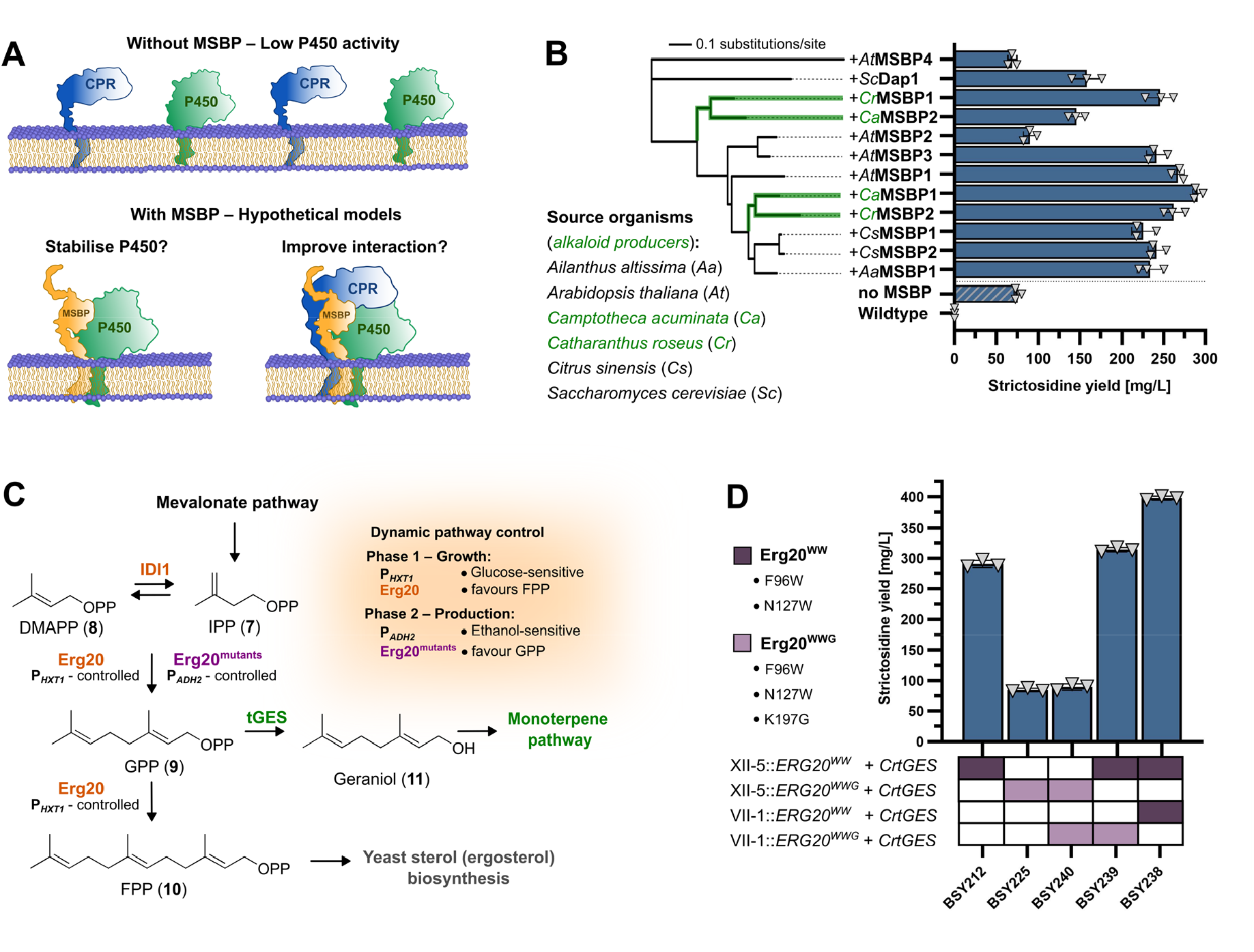
Membrane steroid binding proteins (MSBP) and a second copy of the double mutant gene *ERG20*^*WW*^ improve strictosidine (3) production in yeast. Up to 398 mg/L strictosidine (**3**) were produced. **A)** Overview of hypothetical MSBP function according to (Gou et al., 2018). **B)** Phylogenetic tree of the twelve MSBP homologues tested here and their effect on strictosidine (**3**) titre. The unrooted neighbour-joining tree was constructed based on amino acid sequences. Bar plot shows means ± standard deviation and data points from three independently grown yeast colonies. Species codes and MSBP tree branches of the monoterpene indole alkaloid producing plants *C. acuminata* (*Ca*) and *C. roseus* (*Cr*) are highlighted in green. **C)** Schematic pathway showing the branchpoint function of Erg20. In the growth phase, wild-type Erg20 (orange) under control of the glucose-sensitive promoter P*HXT1* favours FPP production towards yeast sterols (Kong et al., 2023), whereas in the production phase, Erg20 mutants (purple) under control of the ethanol-sensitive promoter P*ADH2* favour GPP production towards monoterpenes. **D)** Effect of including one or two copies of the *ERG20* mutants *ERG20*^*WW*^ (F96W, N127W) and *ERG20*^*WWG*^ (F96W, N127W, K197G) on strictosidine (**3**) levels. Bar plot shows means ± standard deviation and data points from three independently grown yeast colonies.

To further increase strictosidine (**3**) production, we next aimed to increase flux toward the monoterpene key intermediate secologanin (**19**). This intermediate and its precursors, like all monoterpenes, are derived from GPP (**9**). In yeast, GPP (**9**) occurs as a transient intermediate produced by the farnesyl pyrophosphate synthase Erg20 (Fig. 3C). Under normal conditions, Erg20 further prenylates GPP (**9**) en route to FPP (**10**) and ultimately yeast sterols (ergosterol). Thereby, Erg20 severely limits metabolic flux towards monoterpenes. Since deletion of *ERG20* is lethal, mutant variants favouring formation of GPP (**9**) over FPP (**10**) have been developed (Ignea et al., 2014; J. Zhao et al., 2017). The double mutant Erg20^WW^ (F96W, N127W) has been used previously to enhance monoterpene indole alkaloid production in yeast (Bradley et al., 2023; Zhang et al., 2022) and was therefore incorporated into our base strain BSY84. A recently reported triple mutant, Erg20^WWG^ with an additional K197G mutation, further shifts product formation toward GPP (Bernard et al., 2024; Zhang et al., 2019), but has not been tested in multistep monoterpene pathways yet. To assess its performance, we constructed strains differing in the type and copy number of *ERG20* mutants (Fig. 3D, Supplementary Fig. 9). Surprisingly, replacing *ERG20*^*WW*^ with *ERG20*^*WWG*^ (strain BSY225) led to a drastic drop by 71% in strictosidine (**3**) production to 85 mg/L. This effect was also not alleviated by introducing a second copy of *ERG20*^*WWG*^ (strain BSY240). A strain carrying a combination of *ERG20*^*WW*^ and *ERG20*^*WWG*^ (strain BSY239) performed comparably to only a single copy of *ERG20*^*WW*^. However, addition of a second copy of *ERG20*^*WW*^ increased strictosidine (**3**) titre by 37% to 398 mg/L in strain BSY238.

### 3.3. Discovery of a geraniol overoxidation shunt product

When we analysed chromatograms of our engineered yeast strains, we noticed that – besides strictosidine – another major peak was present that showed a comparably strong increase in peak area during our engineering efforts but could not be assigned to any of the known pathway intermediates and shunt products (Fig. 4A). The UV spectrum of this peak suggested that it did not contain indole and might therefore be a monoterpene shunt product (Fig. 4A). To clarify its structure, we isolated the compound **6** from culture supernatants for structure elucidation. NMR spectroscopy revealed that this compound **6** is (2*E*,6*E*)-2,6-dimethylocta-2,6-dienedioic acid (DOA), and is likely an overoxidation shunt product of 8-oxogeranial (**13**) and related intermediates. To the best of our knowledge, this shunt product has not been reported before in the context of engineering iridoid biosynthesis.

**Fig. 4.**
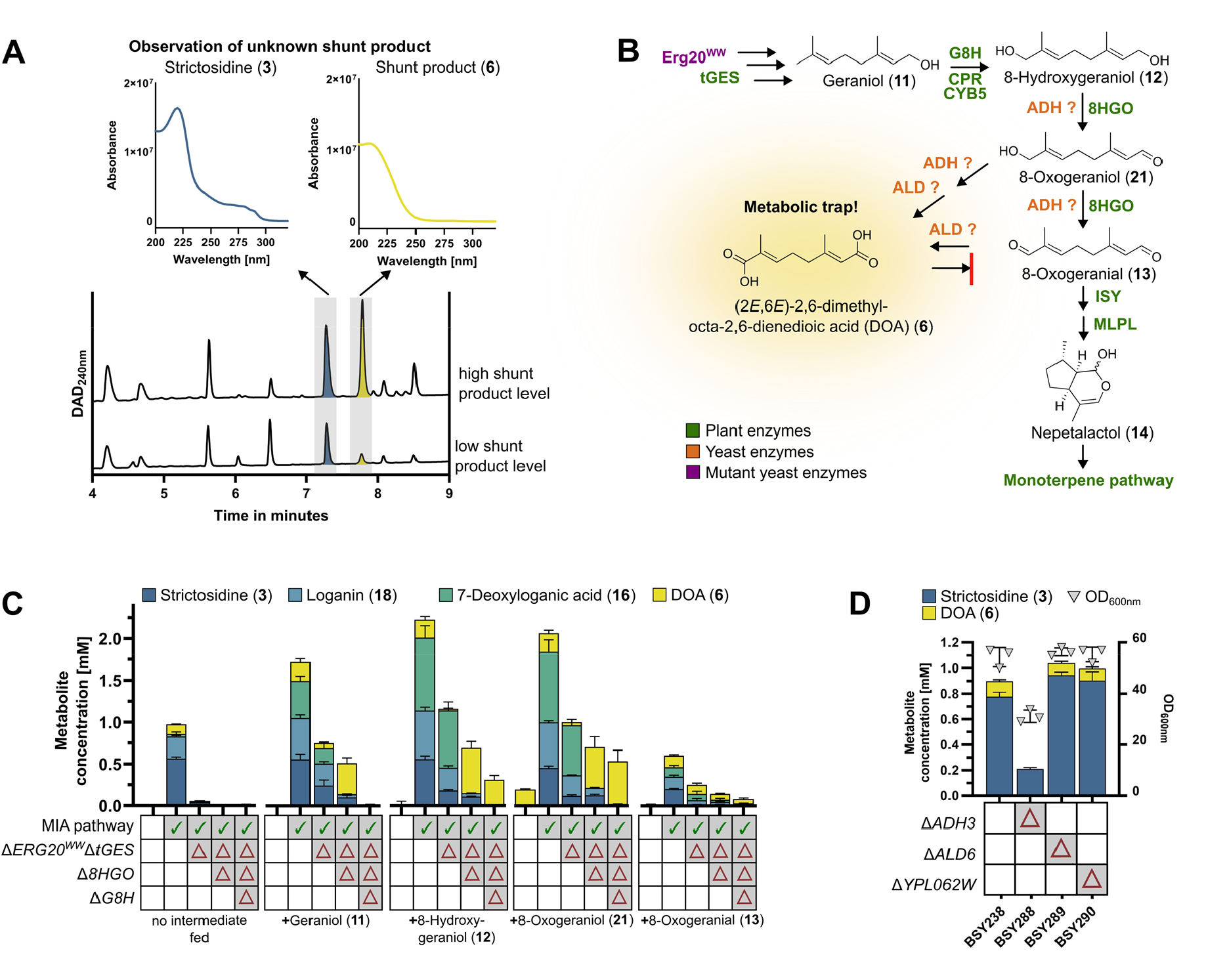
Identification of the monoterpene shunt product (2*E*,6*E*)-2,6-dimethylocta-2,6-dienedioic acid (DOA) (6). **A)** Observation of an unknown shunt product peak in yeast strains producing high strictosidine **(3)** titres. Comparison of UV spectra with strictosidine **(6)** indicated lack of an indole ring system. **B)** Proposed pathways leading to DOA **(6)**. The structure of DOA (**6**) was confirmed by NMR spectroscopy. **C)** Feeding experiment to determine the metabolic origin of DOA **(6)**. 1 mM of the substrates were fed. Stacked bar charts show means ± standard deviation from three independently grown yeast colonies. **D)** Effect of deleting *ADH3, ALD6*, and *YPL062W* on strictosidine (**3**) and DOA (**6**) concentrations. Stacked bar charts show means ± standard deviation of metabolite concentrations (left axis) from three independently grown yeast colonies; OD600 values with standard deviation are additionally shown as grey triangles (right axis). ADH, alcohol dehydrogenase; ALD, aldehyde dehydrogenase.

The structure of DOA (**6**) suggests that it is derived from one or several of the pathway intermediates geraniol (**11**), 8-hydroxygeraniol (**12**), 8-oxogeraniol (**21**), or 8-oxogeranial (**13**). To better understand how DOA (**6**) is formed, we fed various pathway intermediates to different yeast strains and quantified the formation of DOA (**6**), strictosidine (**3**), and the intermediates 7-deoxyloganic acid (**16**) and loganin (**18**) (Fig. 4C, Supplementary Fig. 10). Feeding of 8-oxogeranial (**13**) had a noticeable negative impact on strain growth in terms of OD600, which is not surprising, as this dialdehyde can potentially act as a protein crosslinker; all other tested conditions showed very similar growth behaviour (Supplementary Fig. 10). Notably, formation of DOA (**6**) was already observable in the wildtype strain ST7574 upon feeding of 8-oxogeraniol (**21**) but none of the other intermediates. This suggests that background yeast enzymes can convert 8-oxogeraniol (**21**) into DOA (**6**). When geraniol (**11**) was fed, DOA (**6**) formation was abolished upon deletion of *G8H*, indicating that no yeast background enzyme can functionally replace G8H. In contrast, DOA (**6**) was formed in high titres from 8-hydroxygeraniol (**12**) and 8-oxogeraniol (**21**) even when *8HGO* was deleted. This suggested that DOA (**6**) can be formed from any of the pathway intermediates 8-hydroxygeraniol (**12**), 8-oxogeraniol (**21**), and 8-oxogeranial (**13**). Presumably, this involves alcohol dehydrogenases (ADH) or aldehyde dehydrogenases (ALD), which are common enzymes in yeast and known to act on monoterpenes (Brown et al., 2015; de Smidt et al., 2008; Navarro-Aviño et al., 1999; Ohashi et al., 2021) (Fig. 4B). The most likely candidates are Adh3, which is involved in formation of the structurally related geranic acid (Ohashi et al., 2021), and Ald6, as it is a major cytosolic ALD in yeast (Saint-Prix et al., 2004) and deletion of the *ALD6* core promoter *YPL062W* increased terpenoid production in yeast (Chen et al., 2019). We therefore tested if deletion of *ADH3*, direct deletion of *ALD6*, or *ALD6* downregulation via *YPL062W* deletion might circumvent DOA (**6**) formation and improve strictosidine (**3**) production. Deletion of *ADH3* had an overall strong negative effect on both strictosidine (**3**) production and strain growth (Fig. 4D). In contrast, *ALD6* and *YPL062W* deletion both led to a slight increase in strictosidine (**3**) but no substantial change in DOA (**6**) levels under our common growth conditions (Fig. 4D).

### 3.4. Reprogramming to strictosidinic acid production

After successfully boosting the formation of strictosidine (**3**) in our yeast strains, we started the reprogramming to produce strictosidinic acid (**4**) instead of strictosidine (**3**). In principle, three critical steps were considered for this aim (Fig. 5A): 1) The gene encoding LAMT should be deleted, as it is no longer necessary. 2) The STR from *C. roseus* might not be suitable or at least not optimal for condensing secologanic acid (**4**) with tryptamine (**20**); instead, a different STR homologue might perform better. 3) The P450 SLS might need to be swapped, as it might not efficiently accept loganic acid (**17**) instead of loganin (**18**) as a substrate. Instead, the reported bifunctional P450s CYP72A565/610 from *C. acuminata* could be more suitable for secologanic acid (**4**) formation and at the same time also replace 7DLH as reported (Yang et al., 2019).

**Fig. 5.**
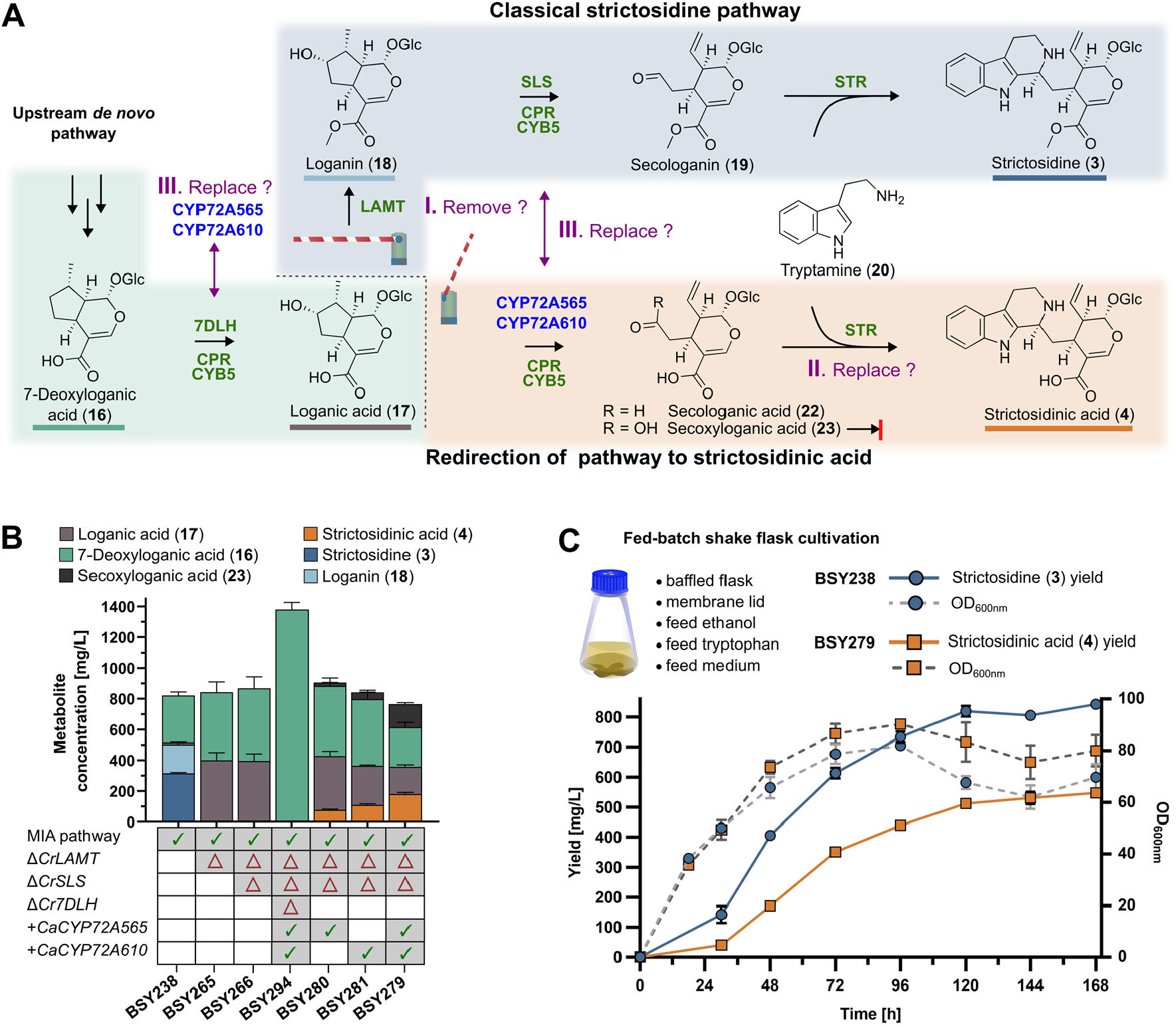
Reprogramming a strictosidine-producing yeast strain to produce strictosidinic acid (4). **A)** Comparison of pathways to strictosidine (**3**) and strictosidinic acid (**4**), respectively. Potential pivots that were targeted for the reprogramming are highlighted in purple. **B)** Step-wise engineering from strictosidine (**3**) production to strictosidinic acid (**4**) production. The stacked bar chart shows means ± standard deviation from three independently grown yeast colonies. **C)** Production of strictosidine (**3**) (strain BSY238) and strictosidinic acid (**4**) (strain BSY279) in a fed-batch shake flask system as time course of product yield (left y axis) and OD600 (right y axis). Strains were grown in 250 mL baffled shake flasks with membrane lids in a fed-batch process with feeding ethanol and concentrated YP medium with additional tryptophan (**5**). Data points in the line graph represent means ± standard deviation from three independently grown yeast colonies.

First, we deleted the *LAMT* gene from our strictosidine (**3**) producer strain BSY238. As expected, this fully abolished strictosidine (**3**) production in the resulting strain BSY265 (Fig. 5B). Instead, 7-deoxyloganic acid (**16**) and the LAMT substrate loganic acid (**17**) accumulated. Next, we wanted to ensure that the *C. roseus* STR would be suitable for producing strictosidinic acid (**4**) instead of strictosidine (**3**). To test if this *Cr*STR tolerates secologanic acid (**22**) instead of secologanin (**19**), we performed a feeding experiment with a yeast strain harbouring only *CrSTR* (BSY43) (Supplementary Fig. 11). As a positive control, we confirmed that the native substrate secologanin (**19**) was converted very efficiently into strictosidine (**3**) up to 400 mg/L of fed substrate. Feeding of crude secologanic acid (**22**), containing ca. 200 mg/L of the desired substrate, resulted in formation of 234 mg/L strictosidinic acid (**4**), corresponding to a molar conversion yield of ca. 85%. This demonstrates that – even though the *C. roseus* STR normally only produces strictosidine (**3**) and not strictosidinic acid (**4**) in its native environment – this enzyme is very well capable of sustaining strictosidinic acid (**4**) production even for high precursor levels at least up to 200 mg/L secologanic acid. Swapping *CrSTR* against a different *STR* gene was therefore not considered necessary.

Finally, we focused on the P450 swaps to enable strictosidinic acid (**4**) production. For this purpose, we deleted the P450 gene *CrSLS*, resulting in strain BSY266; as SLS acts downstream of loganin (**18**), this did not have any noticeable metabolic effect. Afterwards, we replaced the *Cr7DLH* gene with the *CYP72A565* and *CYP72A610* genes from *C. acuminata* (Yang et al., 2019). Surprisingly, even though 7DLH activity was previously reported for CYP72A565 and CYP72A610 *in vitro* (Miller et al., 2021; Yang et al., 2019), the resulting strain BSY294 only produced 7-deoxyloganic acid (**16**), albeit at a very high titre of 1380 mg/L. This suggested that 7DLH cannot be replaced by CYP72A565 and CYP72A610 *in vivo*. Instead of deleting *7DLH*, we therefore simply added *CYP72A565* to the *7DLH*-containing strain BSY266. Gratifyingly, the resulting strain BSY280 produced strictosidinic acid (**4**). The metabolite profile and titres of strain BSY281 containing *CYP72A610* instead of *CYP72A565* looked very similar, corroborating previous reports that both P450s are largely functional equivalent (Miller et al., 2021; Yang et al., 2019). The highest strictosidinic acid (**4**) titre of 181 mg/L was obtained in strain BSY279 containing both *CYP72A565* and *CYP72A610*, even though the total metabolite level decreased slightly and the overoxidation product secoxyloganic acid (**23**) was observed in substantial amounts.

Lastly, we performed a time-course experiment in shake flasks to monitor the growth and metabolite formation of our main strictosidine (**3**) producing strain BSY238 and our main strictosidinic acid (**4**) producer BSY279 (Fig. 5D). Both strains reached their maximum cell density at around 96 h. Metabolite production reached maximum efficiency at around 30 hours, thanks to the auto-inducible promoters used in our system responding to ethanol accumulation. After 168 h (7 days), the highest titres of 843 mg/L strictosidine (**3**) and 548 mg/L strictosidinic acid (**4**) were observed.

## 4. Discussion

### 4.1. Improved production of strictosidine

Even though strictosidine (**3**) has been a highly popular target for metabolic engineering, substantial improvements are still required before biotechnological production of this metabolite and particularly of downstream alkaloids will become economically viable (Zhang et al., 2022). Our study now indicates that MSBP, a second copy of *ERG20*^*WW*^, and avoidance of the shunt product DOA (**6**) are central factors for engineering monoterpene indole alkaloid biosynthetic pathways that should be considered in future studies. Most importantly, we demonstrate that *MSBP* co-expression increases strictosidine (**3**) production by approximately fourfold, confirming that this protein, previously shown to boost triterpenoid production in yeast (Liu et al., 2024), also strongly increases monoterpene indole alkaloid production. Recently, beneficial activity of MSBP was also observed for further plant alkaloid pathways in yeast (Xu et al., 2025). It has been unknown whether MSBPs align with different biosynthetic pathways. By screening a panel of twelve MSBP homologues from different sources, we determined that most showed very similar performance independent of its biological origin, even though four MSBPs (*At*MSBP4, *Sc*Dap1, *At*MSBP2, *Ca*MSBP2) possessed substantially weaker activity. We therefore conclude that inclusion of an MSBP in yeast is very important for pathways involving plant P450s, but the choice of MSBP is not critical, as long as an established *MSBP* gene is selected.

The pivotal role of Erg20 and its mutant variants for controlling flux into monoterpene biosynthesis in yeast has been well recognised in multiple studies (Fischer et al., 2011; Ignea et al., 2014; Jiang et al., 2017; Westfall et al., 2012; Zhao et al., 2016). Recently, the triple mutant Erg20^WWG^ was described to further increase GPP levels over Erg20^WW^ and applied for the production of the simple monoterpenes geraniol (**11**) and (*S*)-limonene (Bernard et al., 2024). These results could also be confirmed for linalool formation in the yeast *Rhodotorula toruloides* (Zheng et al., 2025). However, for production of the simple monoterpene α-terpineol in yeast, Erg20^WWG^ performed worse than Erg20^WW^ (Zhang et al., 2019). Here, we tested Erg20^WWG^ for the first time in a more complex pathway that involves further transformations of monoterpenes. To our surprise, in our system, we observed a drastic decrease in strictosidine levels by 71% when Erg20^WW^ was replaced by Erg20^WWG^ (Fig. 3D); cell growth was not affected by this change (Supplementary Fig. 9). Erg20^WWG^ incorporates the previously reported mutation K197G (Bernard et al., 2024; Fischer et al., 2011). K197 is thought to interact with pyrophosphate groups of the substrates (Bernard et al., 2024; Fischer et al., 2011). One possible explanation why in our system Erg20^WWG^ did not perform as well as reported for simple unmodified monoterpenes is that the mutation K197G might negatively affect the kinetic properties of the enzyme. These effects would become more pronounced in longer pathways, which require a steady supply of geraniol substrate to fuel the downstream pathway. Alternatively, the additional K197G mutation might destabilise the enzyme. Nonetheless, even though Erg20^WW^ outperformed Erg20^WWG^ in our system, Erg20^WW^ still seems to be rate-limiting, as inclusion of a second copy of *ERG20*^*WW*^ led to a 37% increase in strictosidine (**3**) titres. Future studies focused on monoterpenoid and diterpenoid metabolic engineering should therefore consider including at least two copies of *ERG20*^*WW*^.

The high productivity of our strictosidine (**3**) producing strain BSY238 enabled us to identify a yet unreported shunt product of the iridoid pathway, DOA (**6**) (Fig. 4B). It is well known that several intermediates of iridoid biosynthesis are readily converted to various shunt products in yeast (Brown et al., 2015). For this reason, our strain included deletions of the genes *ADH6, ARI1, ATF1, OYE2*, and *OYE3* (Fig. 2), which were previously linked to shunt product formation (Brown et al., 2015; Zhang et al., 2022). Nonetheless, this did not prevent formation of DOA (**6**). Most likely, there are further unrecognised oxidases in yeast, such as ADHs or ALDs, that help to detoxify reactive intermediates in this pathway and lead to DOA (**6**) formation. This toxicity is also underlined by the reduced growth (Supplementary Fig. 10) and metabolite production (Fig. 4C) during our feeding experiments with the potential protein crosslinker 8-oxogeranial (**13**). We suspected that *ADH3* or *ALD6* could be involved in DOA (**6**) formation based on previously reported matching biochemical activity (Chen et al., 2019; Ohashi et al., 2021). *ADH3* deletion had a deleterious effect on overall strictosidine (**3**) production and strain growth, possibly due to changes to ethanol metabolism and a subsequent redox imbalance (de Smidt et al., 2008). While neither deletion of *ALD6* directly nor indirect downregulation *via YPL062W* deletion affected DOA (**6**) formation under our standard growth conditions substantially, we observed a certain reduction of DOA (**6**) levels that was strongly dependent on growth conditions. It is also possible that multiple ADH/ALDs or other enzymes are jointly involved in the formation of DOA (**6**), and no single gene deletion is sufficient to fully circumvent DOA (**6**) formation. In the future, it will be important not only to identify the endogenous yeast enzyme(s) that generate DOA (**6**) but also to further improve the efficiency of the cyclisation step catalysed by ISY (Geu-Flores et al., 2012) to prevent accumulation of toxic intermediates that have to be detoxified to sustain yeast growth.

Our engineered strain BSY238 reached 843 mg/L strictosidine (**3**) after 168 h in a shake flask system, an 8.8-fold increase compared to the highest reported titre for strictosidine (**6**) production in yeast of 91 mg/L in fed-batch microbioreactors (Holtz et al., 2024). In comparison to the ∼30-40 mg/L obtained in culture tubes (Dror et al., 2024), our increase in titre is ∼20-27-fold. We anticipate that further improvements such as optimisation of culture conditions will soon enable gram-scale titres of strictosidine (**3**) as a major milestone en route to microbial production of plant alkaloids for industrial purposes. It will need to be tested how these high levels of strictosidine (**3**) translate to downstream alkaloids.

### 4.2. Production of strictosidinic acid

In this work, we achieved the first production of strictosidinic acid (**4**) in a heterologous organism, based on glucose and tryptophan. Gratifyingly, our strategy to first develop a high-performance strictosidine producing strain and then subsequently reprogramme it towards strictosidinic acid (**4**) production worked very well. In a shake flask fed-batch system, this resulted in a final titre of 548 mg/L strictosidinic acid (**4**) for strain BSY279 after 168 hours (Fig. 5D). Furthermore, our reprogramming strategy also allowed us to gain further insights into the metabolic pathways of monoterpene indole alkaloids. A surprising finding was that neither CYP72A565 nor CYP72A610 from *C. acuminata* could functionally replace 7DLH from *C. roseus* (Fig. 5B), even though the same activity was previously reported *in vitro* (Miller et al., 2021; Yang et al., 2019). This result underlines that caution is required regarding the transferability of *in vitro* data to *in vivo* situations. Nonetheless, the strain BSY294 is an excellent *de novo* producer of 7-deoxyloganic acid (**16**) at a level of 1380 mg/L and could therefore be useful as a platform strain for future metabolic engineering of iridoids.

A second surprise was that STR from *C. roseus*, a previously well-studied enzyme from the most well-studied producer of monoterpene indole alkaloids (Loris et al., 2007; Maresh et al., 2008), could sustain high level production of strictosidinic acid (**4**) without any enzyme engineering, even though strictosidinic acid (**4**) or related metabolites are not known from *C. roseus*. In combination with the previously reported substrate flexibility of STR (Chen et al., 2006; Loris et al., 2007; McCoy et al., 2006), this suggests that there is yet unexploited potential of CrSTR to access further non-natural strictosidine (**3**) and strictosidinic acid (**4**) analogues by synthetic biology even at high titres.

Lastly, our results indicate that there is still substantial room for further strain improvements. Our final strains still contained several hundred mg/L of the incompletely oxidised intermediates 7-deoxyloganic acid (**16**) and loganic acid (**17**), indicating that 7DLH, SLS, CYP72A565, and CYP72A610 activity is still not fully efficient. At the high titres achieved, it will be essential to investigate which factor limits P450 activity the most in our system. Possible bottlenecks could be NADPH supply (Brown et al., 2015; Kong et al., 2023), membrane space on the endoplasmic reticulum (H. Wang et al., 2023; Yang et al., 2023), interactions with redox partners (Yang et al., 2023), or sufficient protein levels of P450 enzymes themselves. On the other hand, not only P450 activity could be limiting, but also the capacity of CrSTR to accept secologanic acid (**22**) at high titres. While we did not see accumulation of secologanic acid (**22**), we observed that secoxyloganic acid (**23**) was formed, which cannot be fused with tryptamine by STR anymore. Similar overoxidation reactivity was observed for other SLS homologues before (Dugé de Bernonville et al., 2015; Miller et al., 2021; Rodríguez-López et al., 2021). To decrease the formation of secoxyloganic acid (**23**), future engineering efforts could concentrate either on improving STR activity to completely convert secologanic acid (**22**) before it is further oxidised, or on engineering CYP72A565/CYP72A610 to avoid this overoxidation side activity.

Our work now enables further investigation of camptothecin (**1**) biosynthesis. First, no strictosidinic acid synthase is known from *C. acuminata* so far. Our yeast strains provide a test system to examine potential strictosidinic acid synthase candidates *in vivo*; alternatively, the high titres of strictosidinic acid produced make the purification of strictosidinic acid (**4**) as a substrate for *in vitro* assays straightforward as well. Furthermore, a major knowledge gap in camptothecin (**1**) biosynthesis is how strictosamide, the next intermediate proposed to occur in the pathway, is formed from strictosidinic acid (**4**) or strictosidine (**3**). With our yeast strains, gene candidates can be directly plugged in to identify the enzyme(s) responsible for strictosamide formation. Ultimately, this will enable complete elucidation and reconstitution of camptothecin (**1**) biosynthesis in yeast to provide a more sustainable production system that is independent of harvesting the original producer trees.

## 5. Conclusion

In this work, we presented a strategy to produce strictosidinic acid (**4**), a key intermediate of the anticancer drug precursor camptothecin (**1**), in engineered yeast from glucose and tryptophan (**5**) at titres up to 548 mg/L in a shake flask system. Our study highlights key factors that enable switching from strictosidine (**3**) to strictosidinic acid (**4**) production. Furthermore, we showed that MSBP and a second copy of *ERG20*^*WW*^ can boost strictosidine (**3**) levels up to 843 mg/L, which allowed us to identify the previously overlooked shunt product DOA (**6**). Within ten years after the first reported production of strictosidine (**3**) in yeast at a level of ca. 0.5 mg/L (Brown et al., 2015), the titre of this key metabolite has now been improved by more than three orders of magnitude, building on contributions from many groups in the field. Our work will facilitate further elucidation of camptothecin (**1**) biosynthesis and will help to produce camptothecin (**1**) and other economically important monoterpene indole alkaloids in engineered microorganisms in the future.

## Acknowledgements

We thank Sarah Krause and Petra Melloh for excellent technical support and Yvonne Leye, Tino Schulenberg and Miriam Fent for excellent horticultural support. The EasyClone-MarkerFree Vector Set (Addgene kit no. 1000000098) and the Expanded EasyClone-MarkerFree toolkit (Addgene kit #1000000190) were gifts from Irina Borodina.

## Data availability

Data will be made available on request.

## Author contributions

**Benedikt Seligmann:** Formal analysis, Investigation, Methodology, Visualization, Writing – original draft. **Shenyu Liu:** Investigation, Methodology. **Mai Huynh:** Resources. **Thu-Thuy T. Dang:** Writing - review & editing, Funding acquisition. **Jakob Franke:** Conceptualization, Funding acquisition, Project administration, Supervision, Writing – original draft.

## Supporting information

Additional Supporting Information can be found online.

