## Supplementary information for "Production of the anticancer drug intermediate strictosidinic acid in engineered yeast"

### Supplementary methods

#### Production of strictosidine (3) and strictosidinic acid (4) by yeast biotransformations

*S. cerevisiae* strains BSY43 expressing *C. roseus* strictosidine synthase gene (*CrSTR*) or BSY44 expressing *CrSTR* and *C. roseus* loganic acid methyltransferase gene (*CrLAMT*) were used for the whole cell production of strictosidine (**3**) and strictosidinic acid (**4**). Cultures of BSY43 or BSY44 were supplemented with substrates (2 mM tryptamine (**20**) and approximately 1 mM of either secologanin (**19**) or secologanic acid (**22**)) 24 h after inoculation and incubated for an additional 96 h to produce strictosidine (**3**) or strictosidinic acid (**4**).

For the purification of strictosidine (**3**), the culture supernatant was purified on XAD-4 resin ($H_2O$-EtOH gradient, 1:0 to 0:1, v/v). Afterwards, the crude product was suspended in MeOH (1 × 20 mL, 2 × 10), filtered, and concentrated under reduced pressure. Sequential flash chromatography was performed for purification. The first step used the KP-Sil 25 g column (solvent A, DCM; solvent B, MeOH; 2 CV 5% B, 10 CV 5–80% B, 14 CV 80% B). The second purification step employed the Sfär C18 D 12 g column (solvent A, $H_2O$; solvent B, MeCN; solvent C, MeOH; solvent D, DCM; 2 CV 95% A with 5% B, 10 CV 95% A with 5% B to 45% A with 55% B, 2 CV 45% A with 55% B, 2 CV 45% A with 55% B to 100% B, 1 CV 100% B, 14 CV 50% C with 50% D). The final purification was performed by semipreparative LC-MS (gradient: 0–10 min, 10–30% B, 10–10.5 min 30–95% B, 10.5–13 min 95% B, 13–13.5 min 95–10% B, 13.5–15 min 10% B).

For the purification of strictosidinic acid (**4**), the culture supernatant was purified on XAD-4 resin using a gradient of $H_2O$-EtOH (1:0 to 0:1, v/v). The crude material was suspended in MeOH (20 mL) and filtered. The filtrate was concentrated under reduced pressure and purified by sequential flash chromatography. The first purification was performed on a KP-Sil 50 g column (solvent A, DCM; solvent B, MeOH; 2 CV 5% B, 10 CV 5–80% B, 2 CV 80% B, 1 CV 80 – 100% B, 2 CV 100% B). The second purification was carried out on an Sfär C18 D 12 g column (solvent A, $H_2O$; B, MeCN; 2 CV 5% B, 10 CV 5–55% B, 2 CV 55–100% B, 2 CV 100% B). The final purification was performed by semipreparative LC-MS (gradient: 0–10 min, 10–30% B, 10–10.5 min 30–95% B, 10.5–13 min 95% B, 13–13.5 min 95–10% B, 13.5–15 min 10% B).

**Semisynthesis of nepetalactol (14) from catnip oil**

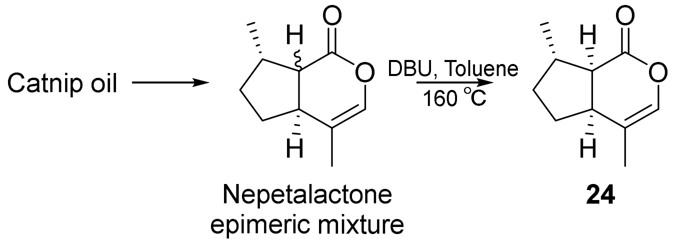


Nepetalactol (**14**) was synthesized *via* (*cis*-*trans*)-nepetalactone (**24**) following a procedure by Liblikas *et al*. (Liblikas et al., 2005). Catnip oil (1.54 g) was first purified by silica column chromatography (petroleum ether : ethyl acetate from 100:1 to 30:1) yielding a mixture (1.32 g) of (*cis*-*trans*)-nepetalactone (**24**) and (*trans*-*cis*)-nepetalactone. The resulting nepetalactone mixture (991 mg, 5.96 mmol, 1.0 eq.) was dissolved in toluene (45 mL) with 1,8-diazabicyclo[5.4.0]undec-7-ene (DBU) (1011 mg, 6.56 mmol, 1.1 eq.) in a round bottom flask. The reaction mixture was refluxed at 160 °C for 2 hours. Solvent was removed under reduced pressure and the residue was purified with silica (petroleum ether : ethyl acetate from 100:0 to 15:1), obtaining (*cis*-*trans*)-nepetalactone (**24**) (566 mg, 57%) as pale-yellow oil. The NMR spectra and HR-MS of (*cis*-*trans*)-nepetalactone (**24**) matched the previous report (Liblikas et al., 2005).

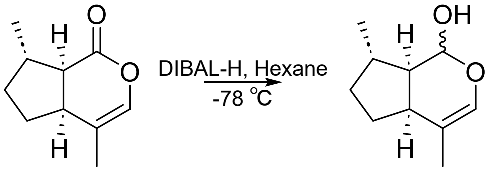


**24** **14**

To a solution of (*cis*-*trans*)-nepetalactone (**24**) (555 mg, 3.34 mmol, 1.0 eq.) in dry hexane (20 mL), diisobutylaluminium hydride (DIBAL-H) (1 N in hexane, 3.7 mL, 3.7 mmol, 1.11 eq.) was added dropwise at −78 °C. The reaction mixture was then stirred for 2 hours at −78 °C and Baeckström reagent ($Na_2SO_4 \cdot 10\ H_2O$ : Celite = 1:1 v/v, 5 g) was added as one portion. The suspension was allowed to warm up to room temperature and followed filtered. The filtrate was collected and concentrated under vacuum. The resulting colorless oil was then purified by silica column chromatography (petroleum ether : ethyl acetate = from 100:1 to 10:1) to give nepetalactol (**14**) as a colorless oil (405 mg, 72%). The NMR spectra and HR-MS of **14** matched the previous report (Liblikas et al., 2005).

**Synthesis of 8-hydroxygeraniol (12), 8-oxogeranial (13) and 8-oxogeraniol (21)**

8-Hydroxygeraniol (**12**), 8-oxogeranial (**13**), and 8-oxogeraniol (**21**) were synthesised from geranyl acetate (**25**) according to procedures by Bat-Erdene *et al.* (Bat-Erdene et al., 2021).

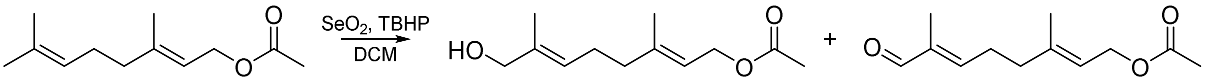


**25** **26** **27**

Geranyl acetate (**25**) (1.008 g, 5.14 mmol, 1.0 eq.) was dissolved in dry dichloromethane (DCM) (26 mL) in a flame dried Schlenk flask and mixed with *tert*-butyl hydroperoxide (TBHP) (5 N in decane, 3 mL, 15 mmol, 2.92 eq.) and $SeO_2$ (224 mg, 2.02 mmol, 0.39 eq.). After 2.5 hours stirring at room temperature, the solvent was removed under reduced pressure. The resulting oil was then dissolved in 50 mL ethyl acetate and washed with deionized water (2 × 20 mL), sat. $NaHCO_3$ (20 mL), deionized water (10 mL) and brine (10 mL). The combined aqueous layers were then back-extracted with ethyl acetate (2 × 50 mL). The organic layers were combined and dried over $Na_2SO_4$ and concentrated *in vacuo*. The resulting oil was purified by silica column chromatography (petroleum ether : ethyl acetate from 6:1 to 2:1) giving 8-hydroxygeranyl acetate (**26**) (484 mg, 44%) and 8-oxogeranyl acetate (**27**) (232 mg, 21%). The NMR spectra and HR-MS of **26** and **27** matched previous reports (Dawson et al., 1996; Ippoliti et al., 2018).

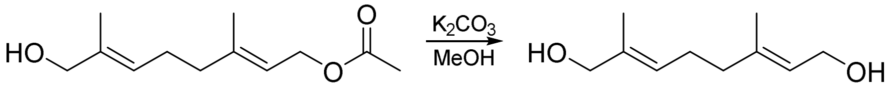


**26** **12**

**8-Hydroxygeraniol (12):** To a solution of 8-hydroxygeranyl acetate (**26**) (480 mg, 2.26 mmol, 1.0 eq.) in MeOH (95%, v/v, 10 mL) $K_2CO_3$ (157 mg, 1.14 mmol, 0.5 eq.) was added in one portion. The mixture was stirred at room temperature for 90 mins and afterwards concentrated under reduced pressure. The residue was dissolved in ether (40 mL) and washed with deionized water (2 × 20 mL). The aqueous layers were back-extracted with ether (2 × 20 mL). The organic layers were combined and solvent was removed *in vacuo*. Final purification with silica column chromatography (petroleum ether : ethyl acetate from 1:1 to 1:2) gave 8-hydroxygeraniol (**12**) (347 mg, 90%) as a colorless oil. The NMR spectra and HR-MS of **12** matched the previous report (Ippoliti et al., 2018).

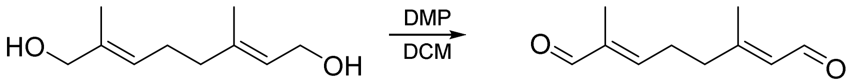


**10** **13**

**8-Oxogeranial (13):** In a round bottom flask flushed with $N_2$, 8-hydroxygeraniol (**12**) (222 mg, 1.30 mmol, 1.0 eq.) was dissolved in dry DCM (13 mL). Dess–Martin periodinane (DMP) (1.33 g, 3.14 mmol, 2.4 eq.) was added at room temperature in one portion. After 80 mins the reaction was quenched by adding a solution (12 mL) of sat. $NaHCO_3$, sat. $Na_2S_2O_3$ and deionized water (1:1:1, v/v). The DCM layer was collected and the aqueous layer was extracted with DCM (3 × 13 mL). All DCM layers were combined and dried over $Na_2SO_4$. The solvent was removed under reduced pressure and the resulting residue was purified by silica column chromatography (petroleum ether : ethyl acetate = 2:1) giving 8-oxogeranial (**13**) as a colorless oil. The NMR spectra and HR-MS of **13** matched previous report (Dawson et al., 1996).

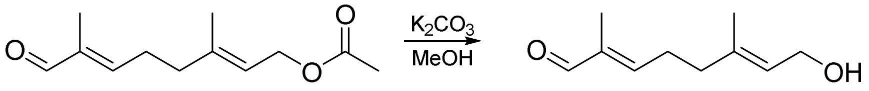

27 21

**8-Oxogeraniol (21):** To a solution of 8-oxogeranyl acetate (**27**) (231 mg, 1.1 mmol, 1.0 eq.) in MeOH (95%, v/v, 10 mL), $K_2CO_3$ (95 mg, 0.69 mmol, 0.63 eq.) was added in one portion. The mixture was stirred at room temperature for 75 mins and afterwards concentrated under reduced pressure. The residue was dissolved in ether (20 mL) and washed with deionized water (2 × 10 mL). The aqueous layers were back-extracted with ether (2 × 10 mL). The organic layers were combined and solvent was removed *in vacuo*. Final purification by silica column chromatography (petroleum ether : ethyl acetate = 2:1) gave 8-oxogeraniol (**21**) (137 mg, 73%) as a colorless oil. The NMR spectra and HR-MS of **21** matched the previous report (Dawson et al., 1996).

**Preparation of 7-deoxyloganic acid (7-DLA) (16) by yeast biotransformation**

For the preparation of 7-deoxyloganic acid (**16**) 2 mM nepetalactol (**14**) was fed to 250 mL of a 24 hours old yeast culture of BSY28 in baffled flasks. After 24 hours incubation, the sterile filtrated yeast culture supernatant (250 mL) was concentrated under reduced pressure and the residue was purified by two rounds of flash chromatography (1 – KP-Sil 50 g; A: DCM, B: MeOH; 2 CV 5% B, 10 CV 5% - 80% B, 3 CV 80% B; 2 – SNAP 10 g; A: DCM, B: MeOH; 1 CV 5% - 15% B, 55 CV 15% B, 2 CV 15% - 50% B, 7 CV 50% B) followed by silica column chromatography (DCM : MeOH with 0.1% formic acid from 19:1 to 17:3). Final purification was carried out by semipreparative LC-MS (Kinetex 5 µm C18 100 Å 250 x 10 mm, A: $H_2O$ + 0.1 % v/v formic acid, B: MeCN+ 0.1 % v/v formic acid; flowrate: 5 mL/min; temperature: 40 °C; gradient: 0-10 min 15-21.7% B, 10-11 min 21.7-100% B, 11-13 min 100% B, 13-13.5 min 100-15% B, 13.5-15 min 15% B). 7-Deoxyloganic acid (**16**) (5.3 mg) was obtained as a pale-yellowish solid. The NMR spectra and HR-MS of **16** matched previously reported data (Teng et al., 2005).

### Supplementary figures

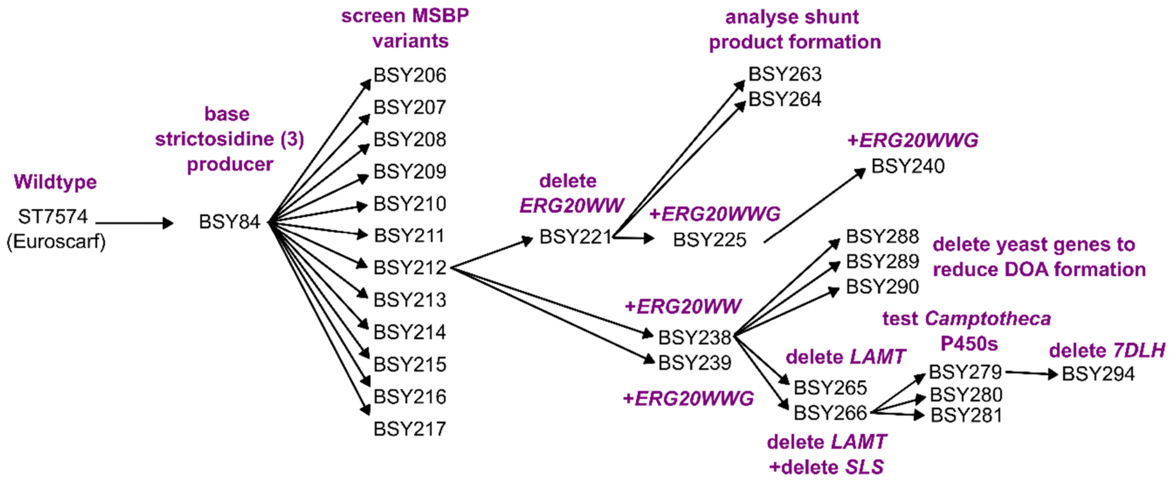


**Supplementary Fig. 1. Relationships of yeast strains generated in this study.**

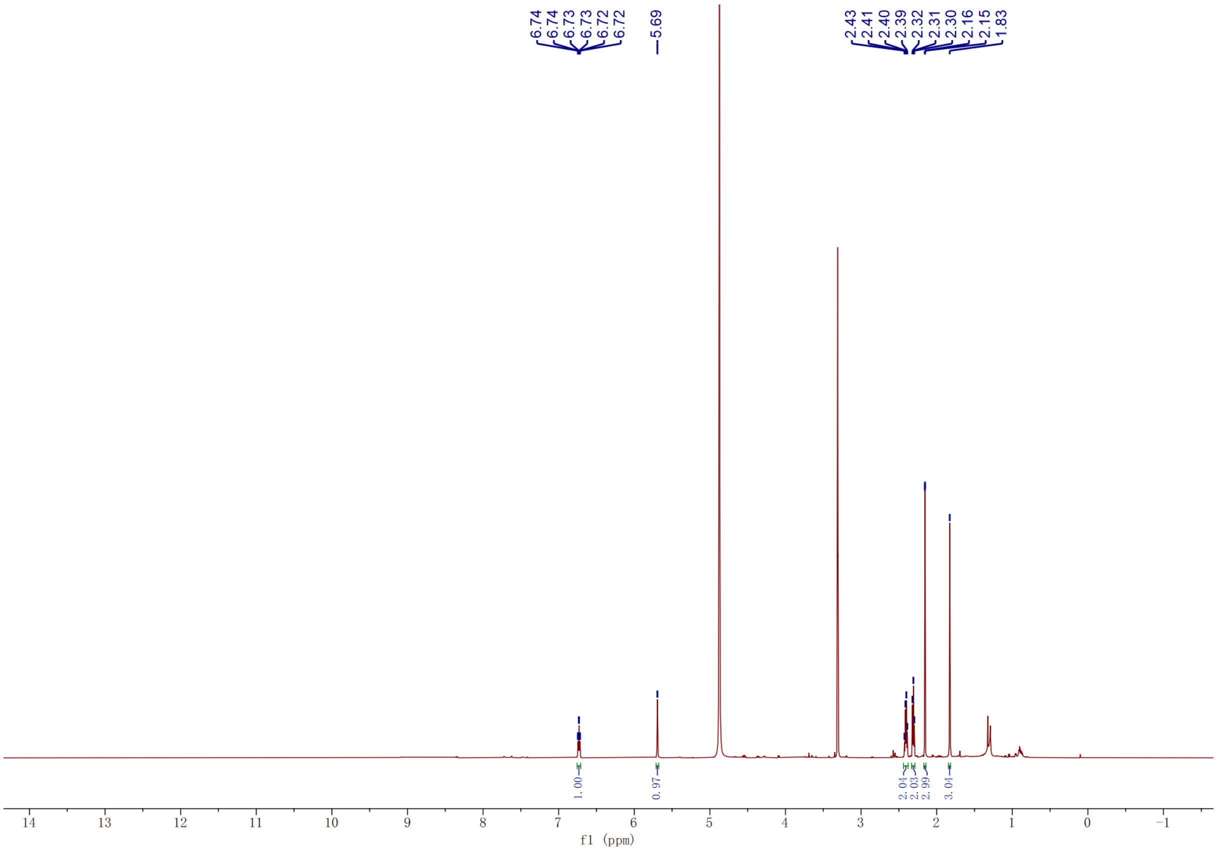


**Supplementary Fig. 2. $^1$H spectrum of (2*E*,6*E*)-2,6-dimethylocta-2,6-dienedioic acid (DOA) (6) (MeOH-$d_4$, 600 MHz, 298 K).**

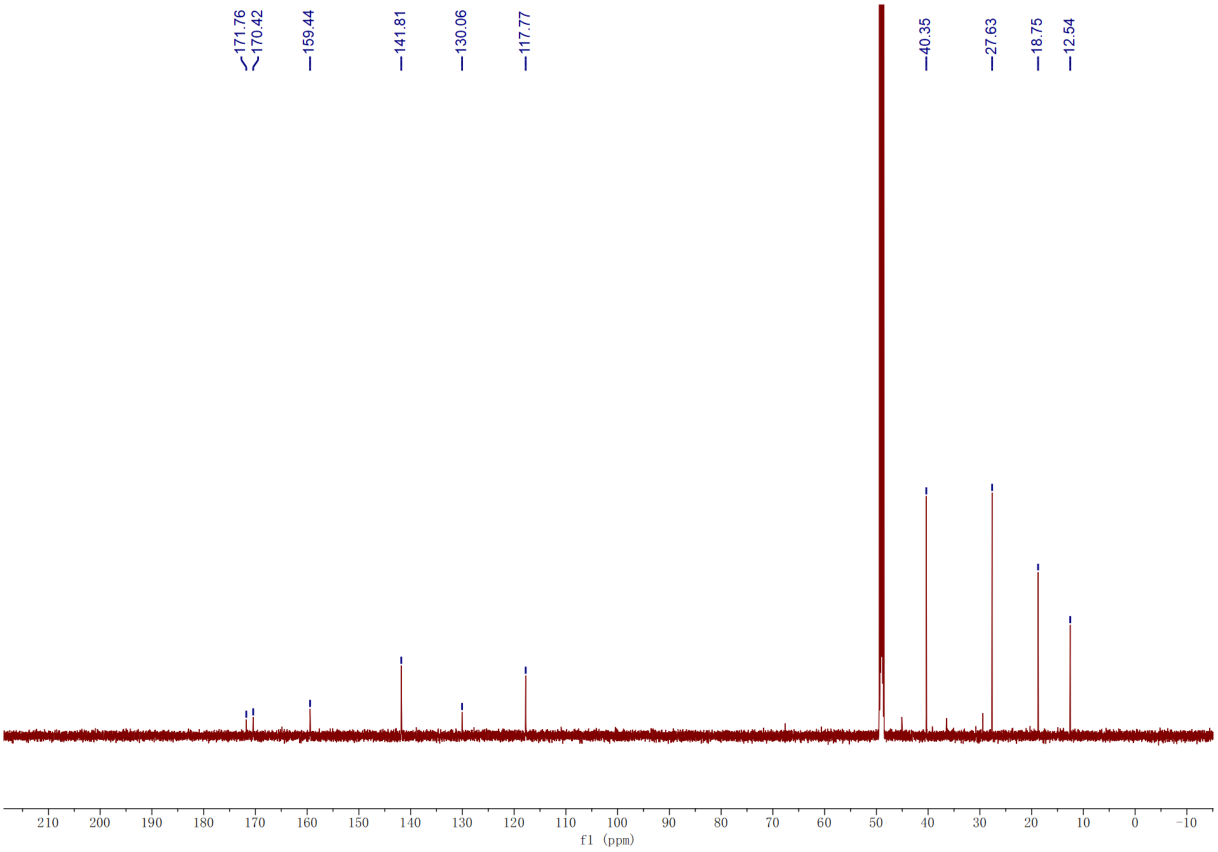


**Supplementary Fig. 3.** $^{13}$**C spectrum of (2*E*,6*E*)-2,6-dimethylocta-2,6-dienedioic acid (DOA) (6) (MeOH-d$_4$, 151 MHz, 298 K).**

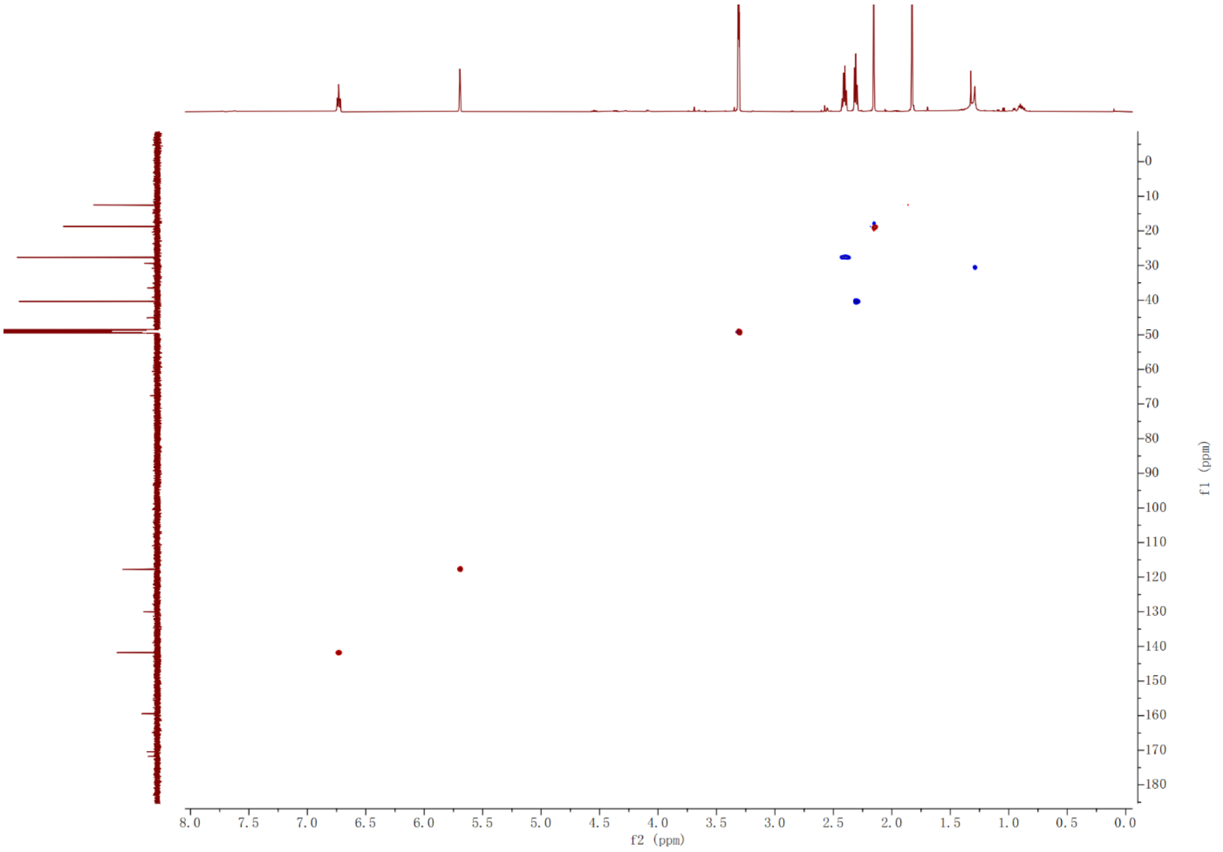


**Supplementary Fig. 4. HSQC spectrum of (2*E*,6*E*)-2,6-dimethylocta-2,6-dienedioic acid (DOA) (6) (MeOH-d$_4$, 600 MHz, 298 K).**

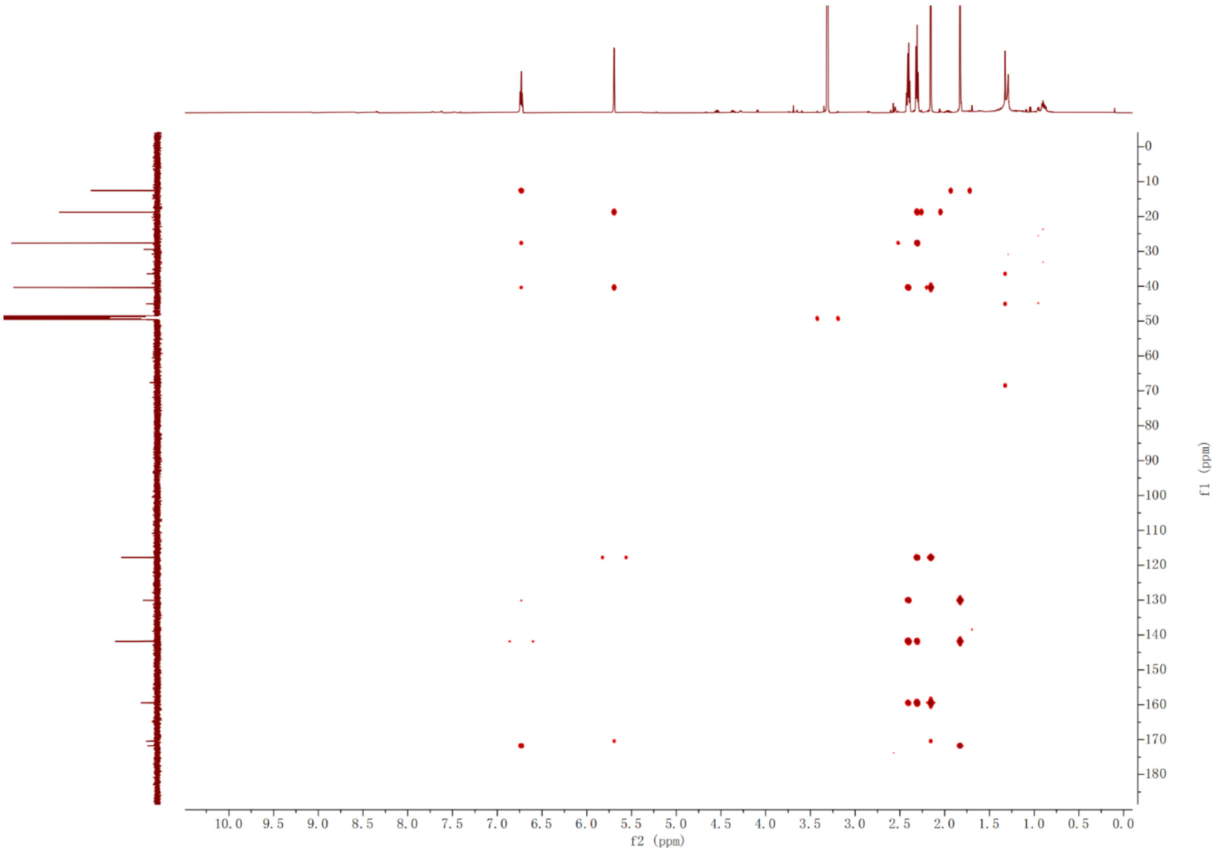


**Supplementary Fig. 5. HMBC spectrum of (2*E*,6*E*)-2,6-dimethylocta-2,6-dienedioic acid (DOA) (6) (MeOH-$d_4$, 600 MHz, 298 K).**

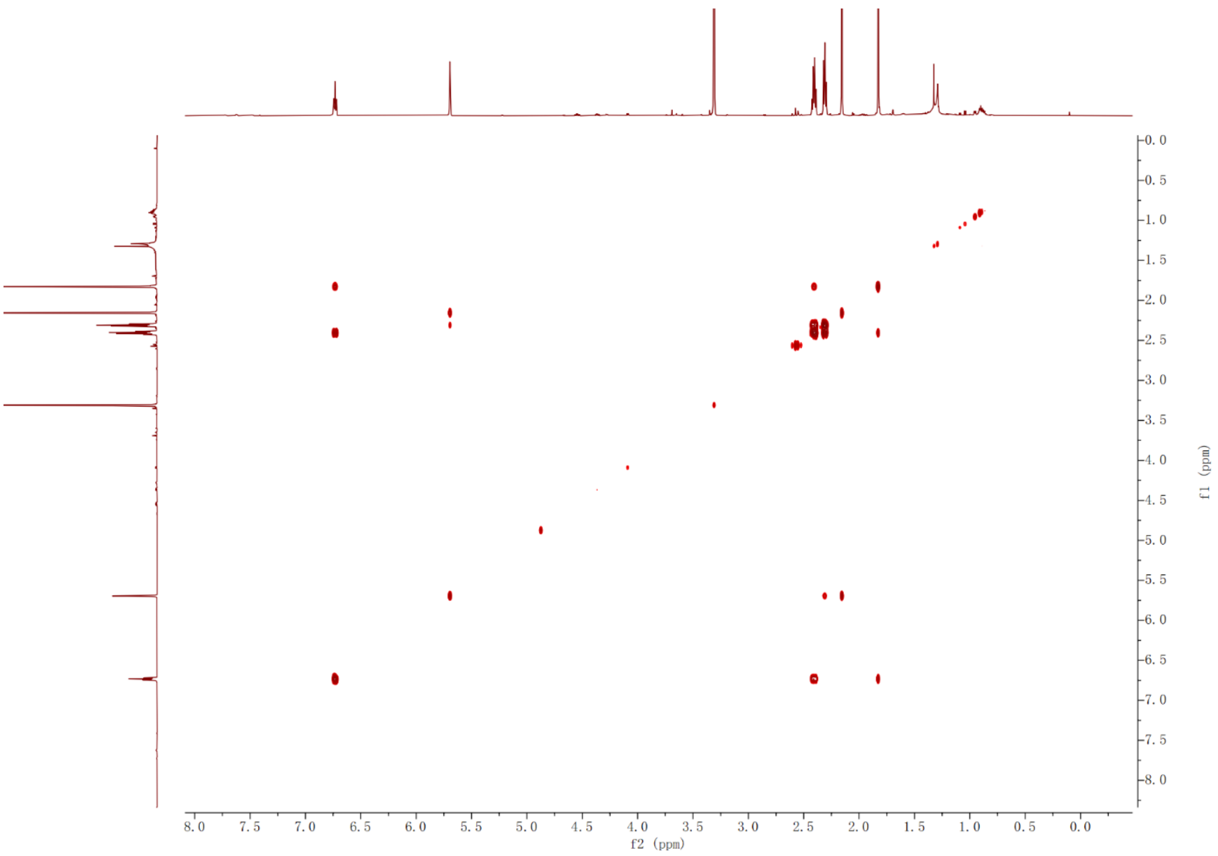


**Supplementary Fig. 6. COSY spectrum of (2*E*,6*E*)-2,6-dimethylocta-2,6-dienedioic acid (DOA) (6) (MeOH-$d_4$, 600 MHz, 298 K).**

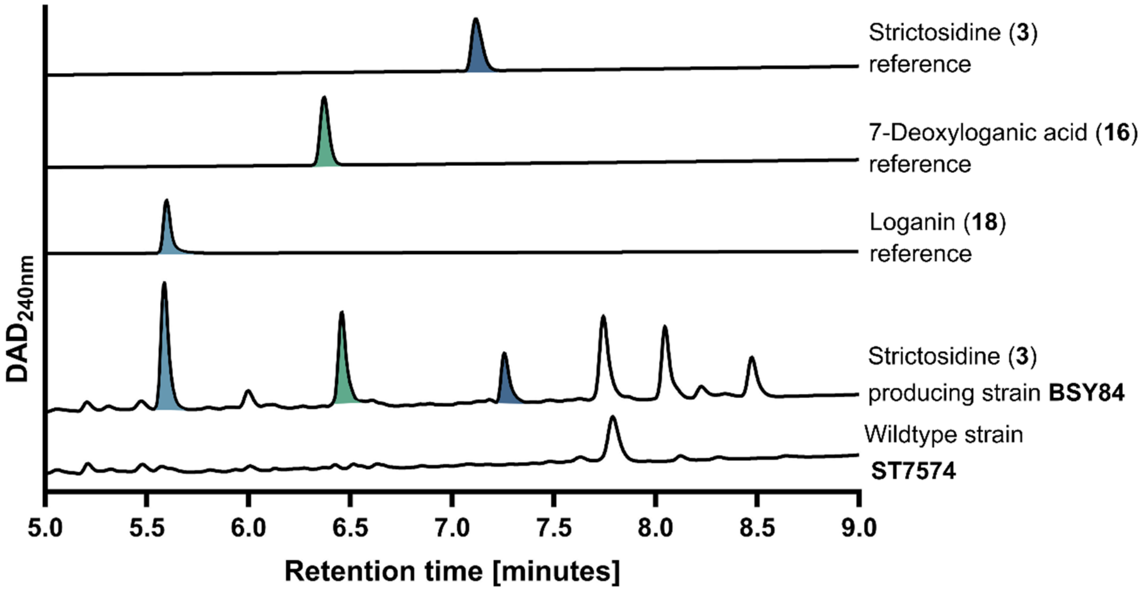


**Supplementary Fig. 7. Metabolite profile of the strictosidine (3) platform strain BSY84.**

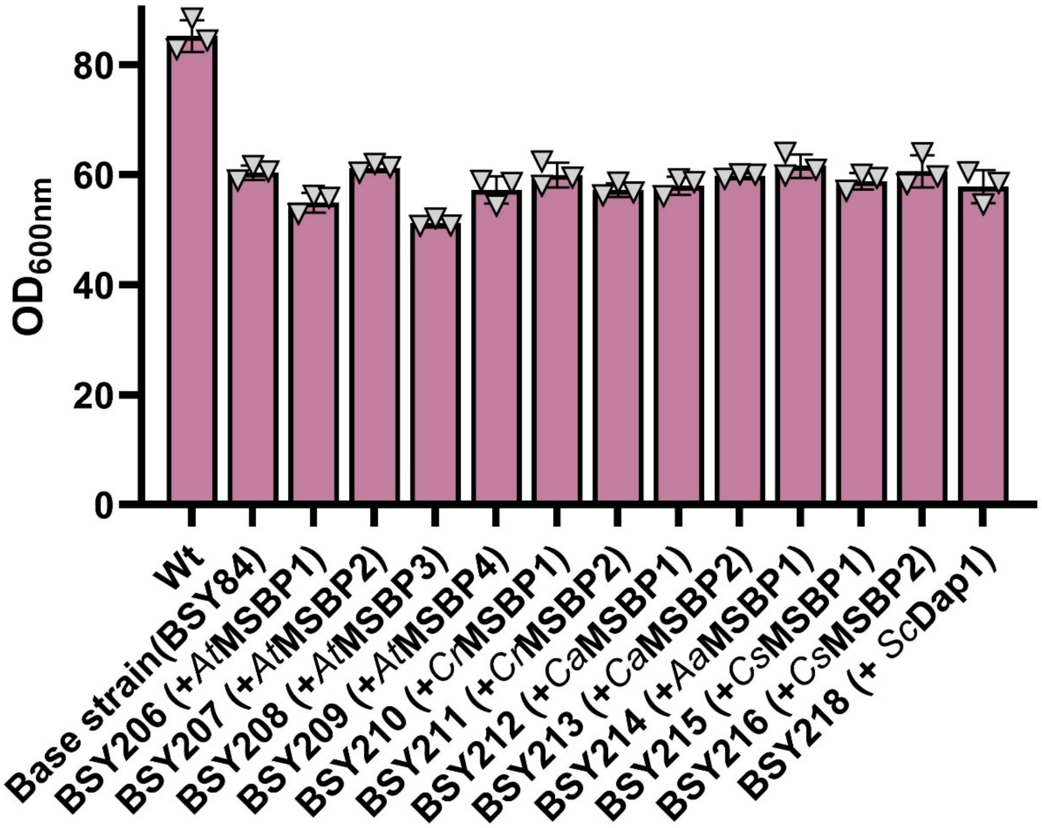


**Supplementary Fig. 8. Growth of yeast strains expressing *MSBP* homologues.**

$OD_{600nm}$ values corresponding to metabolite production shown in Fig. 3B. Bar plot shows means ± standard deviation and data points from three independently grown yeast colonies.

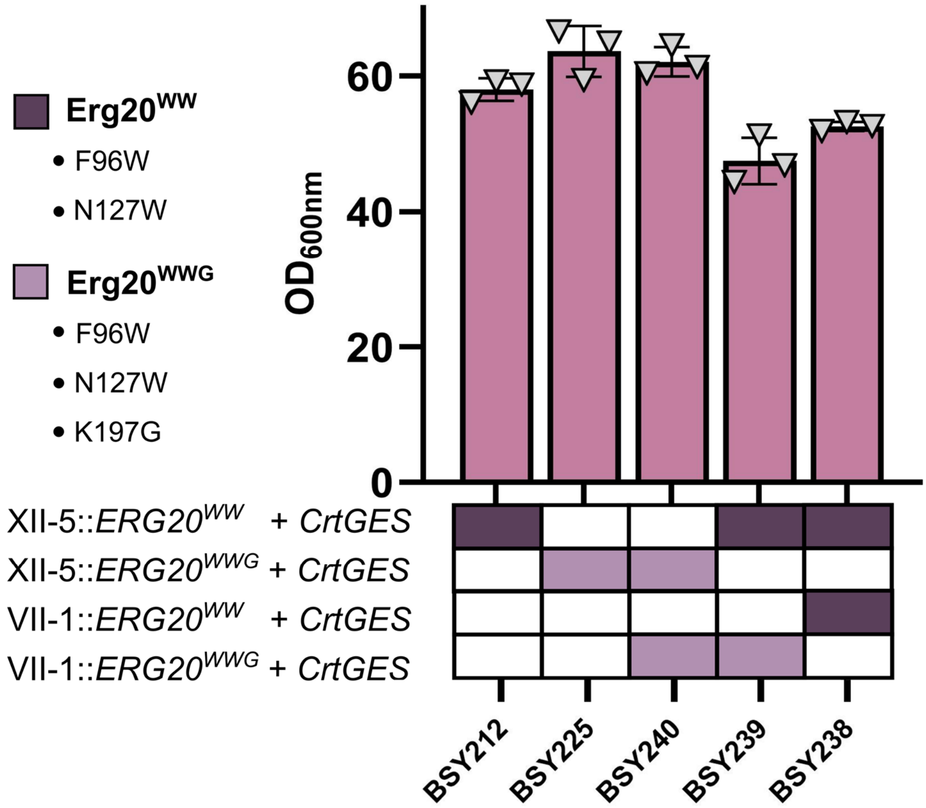


**Supplementary Fig. 9. Growth of yeast strains harbouring *ERG20* mutant variants.**

$OD_{600nm}$ values corresponding to metabolite production shown in Fig. 3D. Bar plot shows means ± standard deviation and data points from three independently grown yeast colonies.

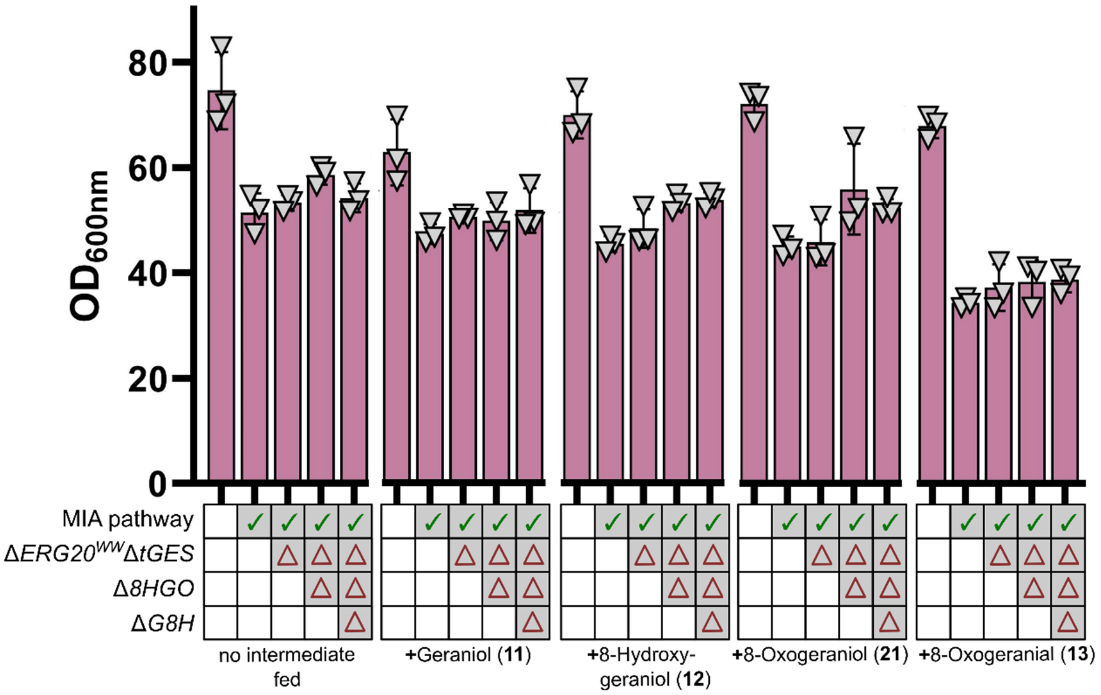


**Supplementary Fig. 10. Growth of yeast strains upon feeding of pathway intermediates.**

$OD_{600nm}$ values corresponding to metabolite production shown in Fig. 4C. Bar plot shows means ± standard deviation and data points from three independently grown yeast colonies.

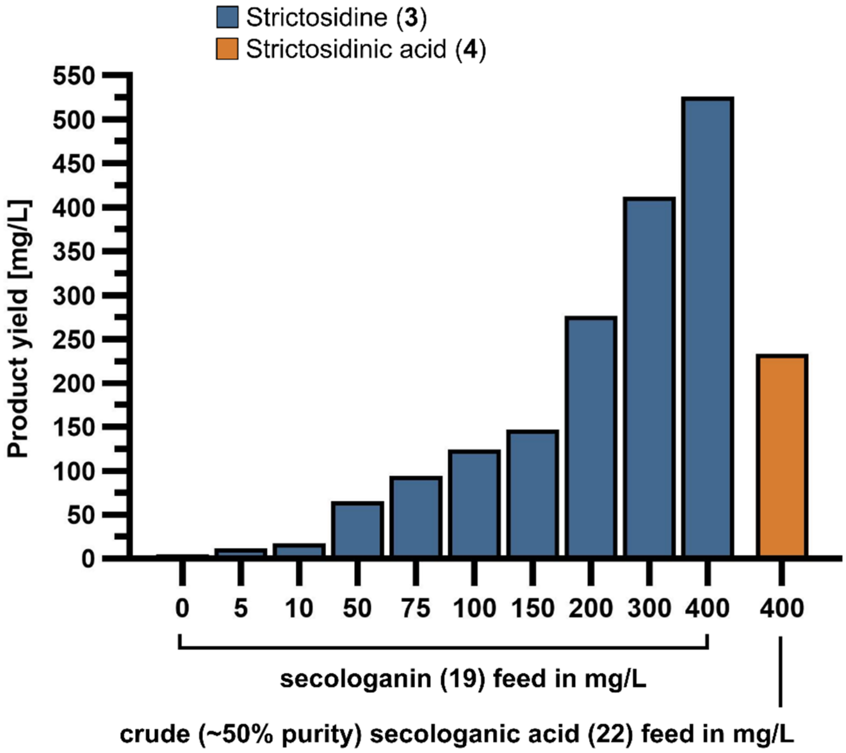


**Supplementary Fig. 11. Testing the conversion efficiency of *Cr*STR by feeding secologanin (19) or secologanic acid (23) to yeast strain BSY43.**

### Supplementary tables

#### Supplementary Table 1. List of yeast strains used in this study.

Genotypic parts in bold indicate changes compared to the parent strain.

$^{opt}$ indicates codon-optimised genes.

| Name | Genotype | Reference |
|---|---|---|
| ST7574 | MATa; HIS3; TRP1; LEU2; TRP1; URA3; MAL2-8c SUC2 + pCfB2312 (2µ cas9 KanMX) | Euroscarf |
| BSY1 | ST7574, **X-2::$P_{ADH2}$-CrCPR-$T_{ADH1}$, $P_{PCK1}$-CrCYB5-$T_{CYC1}$** | This study |
| BSY28 | ST7574, X-2::$P_{ADH2}$-CrCPR-$T_{ADH1}$, $P_{PCK1}$-CrCYB5-$T_{CYC1}$ ; **XI-2::$P_{PCK1}$-CrIO-$T_{ADH1}$, $P_{ADH2}$-Cr7DLGT-$T_{CYC1}$ ; XII-2::$P_{PCK1}$-CrCYPADH-$T_{ADH1}$** | This study |
| BSY43 | ST7574, **XI-5::$P_{ADH2}$-CrSTR-$T_{ADH1}$** | This study |
| BSY44 | ST7574, **XI-5::$P_{ADH2}$-CrSTR-$T_{ADH1}$; X-4::$P_{ADH2}$-CrLAMT-$T_{ADH1}$** | This study |
| BSY84 | ST7574, X-2::$P_{ADH2}$-CrCPR-$T_{ADH1}$, $P_{PCK1}$-CrCYB5-$T_{CYC1}$ ; **XI-2::$P_{PCK1}$-CrIO-$T_{ADH1}$, $P_{ADH2}$-Cr7DLGT-$T_{CYC1}$ ; XII-2::$P_{ADH2}$-CrCYPADH-$T_{ADH1}$, $P_{ICL1}$-Cr7DLH-$T_{CYC1}$ ; XI-3::$P_{ICL1}$-CrLAMT-$T_{ADH1}$, $P_{ADH2}$-CrSLS-$T_{CYC1}$ ; XI-5::$P_{ADH2}$-CrSTR-$T_{ADH1}$ ; X-3::$P_{PCK1}$-CrGOR-$T_{ADH1}$, $P_{ADH2}$-CrG8H-$T_{CYC1}$ ; ΔOYE2, ΔOYE3 ; ΔATF1, ΔARI1 ; ΔADH6 ; X-4::$P_{ADH2}$-NcISY2-$T_{ADH1}$, $P_{ICL1}$-NcMLPLa-$T_{CYC1}$ ; XII-5::$P_{MLS1}$-tCrGES-$T_{ADH1}$, $P_{ADH2}$-ScERG20$^{WW}$-$T_{CYC1}$ ;XI-1:: $P_{PCK1}$-SctHMGR-$T_{ADH1}$, $P_{MLS1}$-ScIDI1-$T_{CYC1}$; $P_{ERG20\Delta}$::$P_{HXT1}$ ; XII4::$P_{ADH2}$-CrTDC-$T_{ADH1}$, $P_{MLS1}$-ScZWF1-$T_{CYC1}$** | This study |
| BSY101 | ST7574, X-2::$P_{ADH2}$-CrCPR-$T_{ADH1}$, $P_{PCK1}$-CrCYB5-$T_{CYC1}$ ; XI-2::$P_{PCK1}$-CrIO-$T_{ADH1}$, $P_{ADH2}$-Cr7DLGT-$T_{CYC1}$ ; XII-2::$P_{ADH2}$-CrCYPADH-$T_{ADH1}$, $P_{ICL1}$-Cr7DLH-$T_{CYC1}$ ; XI-3::$P_{ICL1}$-CrLAMT-$T_{ADH1}$, $P_{ADH2}$-CrSLS-$T_{CYC1}$ ; XI-5::$P_{ADH2}$-CrSTR-$T_{ADH1}$ ; X-3::$P_{PCK1}$-CrGOR-$T_{ADH1}$, $P_{ADH2}$-CrG8H-$T_{CYC1}$ ; ΔOYE2, ΔOYE3 ; ΔATF1, ΔARI1 ; ΔADH6 ; X-4::$P_{ADH2}$-NcISY2-$T_{ADH1}$, $P_{ICL1}$-NcMLPLa-$T_{CYC1}$ ; XII-5::$P_{MLS1}$-tCrGES-$T_{ADH1}$, $P_{ADH2}$-ScERG20$^{WW}$-$T_{CYC1}$ ;XI-1:: $P_{PCK1}$-SctHMGR-$T_{ADH1}$, $P_{MLS1}$-ScIDI1-$T_{CYC1}$; $P_{ERG20\Delta}$::$P_{HXT1}$ ; XII4::$P_{ADH2}$-CrTDC-$T_{ADH1}$, $P_{MLS1}$-ScZWF1-$T_{CYC1}$ ;**IV-1::$P_{ADH2}$-CaMDR-$T_{ADH1}$** | This study |
| BSY206 | ST7574, X-2::$P_{ADH2}$-CrCPR-$T_{ADH1}$, $P_{PCK1}$-CrCYB5-$T_{CYC1}$ ; XI-2::$P_{PCK1}$-CrIO-$T_{ADH1}$, $P_{ADH2}$-Cr7DLGT-$T_{CYC1}$ ; XII-2::$P_{ADH2}$-CrCYPADH-$T_{ADH1}$, $P_{ICL1}$-Cr7DLH-$T_{CYC1}$ ; XI-3::$P_{ICL1}$-CrLAMT-$T_{ADH1}$, $P_{ADH2}$-CrSLS-$T_{CYC1}$ ; XI-5::$P_{ADH2}$-CrSTR-$T_{ADH1}$ ; X-3::$P_{PCK1}$-CrGOR-$T_{ADH1}$, $P_{ADH2}$-CrG8H-$T_{CYC1}$ ; ΔOYE2, ΔOYE3 ; ΔATF1, ΔARI1 ; ΔADH6 ; X-4::$P_{ADH2}$-NcISY2-$T_{ADH1}$, $P_{ICL1}$-NcMLPLa-$T_{CYC1}$ ; XII-5::$P_{MLS1}$-tCrGES-$T_{ADH1}$, $P_{ADH2}$-ScERG20$^{WW}$-$T_{CYC1}$ ; XI-1:: $P_{PCK1}$-SctHMGR-$T_{ADH1}$, $P_{MLS1}$-ScIDI1-$T_{CYC1}$; $P_{ERG20\Delta}$::$P_{HXT1}$ ; XII-4::$P_{ADH2}$-CrTDC-$T_{ADH1}$, $P_{MLS1}$-ScZWF1-$T_{CYC1}$ ; **II-1:: $P_{ADH2}$-AtMSBP1**-$T_{ADH1}$ | This study |
| BSY207 | ST7574, X-2::$P_{ADH2}$-CrCPR-$T_{ADH1}$, $P_{PCK1}$-CrCYB5-$T_{CYC1}$ ; XI-2::$P_{PCK1}$-CrIO-$T_{ADH1}$, $P_{ADH2}$-Cr7DLGT-$T_{CYC1}$ ; XII-2::$P_{ADH2}$-CrCYPADH-$T_{ADH1}$, $P_{ICL1}$-Cr7DLH-$T_{CYC1}$ ; XI-3::$P_{ICL1}$-CrLAMT-$T_{ADH1}$, $P_{ADH2}$-CrSLS-$T_{CYC1}$ ; XI-5::$P_{ADH2}$-CrSTR-$T_{ADH1}$ ; X-3::$P_{PCK1}$-CrGOR-$T_{ADH1}$, $P_{ADH2}$-CrG8H-$T_{CYC1}$ ; ΔOYE2, ΔOYE3 ; ΔATF1, ΔARI1 ; ΔADH6 ; X-4::$P_{ADH2}$-NcISY2-$T_{ADH1}$, $P_{ICL1}$-NcMLPLa-$T_{CYC1}$ ; XII-5Δ$P_{MLS1}$-tCrGES-$T_{ADH1}$, $P_{ADH2}$-ScERG20$^{WW}$-$T_{CYC1}$ ; XI-1:: $P_{PCK1}$-SctHMGR-$T_{ADH1}$, $P_{MLS1}$-ScIDI1-$T_{CYC1}$; $P_{ERG20\Delta}$::$P_{HXT1}$ ; XII-4::$P_{ADH2}$-CrTDC-$T_{ADH1}$, $P_{MLS1}$-ScZWF1-$T_{CYC1}$ ; **II-1:: $P_{ADH2}$-AtMSBP2**-$T_{ADH1}$ | This study |
| BSY208 | ST7574, X-2::$P_{ADH2}$-CrCPR-$T_{ADH1}$, $P_{PCK1}$-CrCYB5-$T_{CYC1}$ ; XI-2::$P_{PCK1}$-CrIO-$T_{ADH1}$, $P_{ADH2}$-Cr7DLGT-$T_{CYC1}$; XII-2::$P_{ADH2}$-CrCYPADH-$T_{ADH1}$, $P_{ICL1}$-Cr7DLH-$T_{CYC1}$; XI-3::$P_{ICL1}$-CrLAMT-$T_{ADH1}$, $P_{ADH2}$-CrSLS-$T_{CYC1}$; XI-5::$P_{ADH2}$-CrSTR-$T_{ADH1}$ ; X-3::$P_{PCK1}$-CrGOR-$T_{ADH1}$, $P_{ADH2}$-CrG8H-$T_{CYC1}$ ; ΔOYE2, ΔOYE3 ; ΔATF1, ΔARI1 ; ΔADH6 ; X-4::$P_{ADH2}$-NcISY2-$T_{ADH1}$, $P_{ICL1}$-NcMLPLa-$T_{CYC1}$ ; XII-5::$P_{MLS1}$-tCrGES-$T_{ADH1}$, $P_{ADH2}$-ScERG20$^{WW}$-$T_{CYC1}$ ; XI-1:: $P_{PCK1}$-SctHMGR-$T_{ADH1}$, $P_{MLS1}$-ScIDI1-$T_{CYC1}$; $P_{ERG20\Delta}$::$P_{HXT1}$ ; XII-4::$P_{ADH2}$-CrTDC-$T_{ADH1}$, $P_{MLS1}$-ScZWF1-$T_{CYC1}$ ; **II-1:: $P_{ADH2}$-AtMSBP3**-$T_{ADH1}$ | This study |
| BSY209 | ST7574, X-2::$P_{ADH2}$-CrCPR-$T_{ADH1}$, $P_{PCK1}$-CrCYB5-$T_{CYC1}$ ; XI-2::$P_{PCK1}$-CrIO-$T_{ADH1}$, $P_{ADH2}$-Cr7DLGT-$T_{CYC1}$ ; XII-2::$P_{ADH2}$-CrCYPADH-$T_{ADH1}$, $P_{ICL1}$-Cr7DLH-$T_{CYC1}$ ; XI-3::$P_{ICL1}$-CrLAMT-$T_{ADH1}$, $P_{ADH2}$-CrSLS-$T_{CYC1}$ ; XI-5::$P_{ADH2}$-CrSTR-$T_{ADH1}$ ; X-3::$P_{PCK1}$-CrGOR-$T_{ADH1}$, $P_{ADH2}$-CrG8H-$T_{CYC1}$ ; ΔOYE2, ΔOYE3 ; ΔATF1, ΔARI1 ; ΔADH6 ; X-4::$P_{ADH2}$-NcISY2-$T_{ADH1}$, $P_{ICL1}$-NcMLPLa-$T_{CYC1}$ ; XII-5::$P_{MLS1}$-tCrGES-$T_{ADH1}$, $P_{ADH2}$-ScERG20$^{WW}$-$T_{CYC1}$ ; XI-1:: $P_{PCK1}$-SctHMGR-$T_{ADH1}$, $P_{MLS1}$-ScIDI1-$T_{CYC1}$; $P_{ERG20\Delta}$::$P_{HXT1}$ ; XII-4::$P_{ADH2}$-CrTDC-$T_{ADH1}$, $P_{MLS1}$-ScZWF1-$T_{CYC1}$ ; **II-1:: $P_{ADH2}$-AtMSBP4**-$T_{ADH1}$ | This study |

| | | |
|---|---|---|
| BSY210 | ST7574, X-2::$P_{ADH2}$-CrCPR-$T_{ADH1}$, $P_{PCK1}$-CrCYB5-$T_{CYC1}$ ; XI-2::$P_{PCK1}$-CrIO-$T_{ADH1}$, $P_{ADH2}$-Cr7DLGT-$T_{CYC1}$ ; XII-2::$P_{ADH2}$-CrCYPADH-$T_{ADH1}$, $P_{ICL1}$Cr7DLH-$T_{CYC1}$ ; XI-3::$P_{ICL1}$-CrLAMT-$T_{ADH1}$, $P_{ADH2}$-CrSLS-$T_{CYC1}$ ; XI-5::$P_{ADH2}$-CrSTR-$T_{ADH1}$ ; X-3::$P_{PCK1}$-CrGOR-$T_{ADH1}$, $P_{ADH2}$-CrG8H-$T_{CYC1}$ ; ΔOYE2, ΔOYE3 ; ΔATF1, ΔARI1 ; ΔADH6 ; X-4::$P_{ADH2}$-NcISY2-$T_{ADH1}$, $P_{ICL1}$-NcMLPLa-$T_{CYC1}$ ; XII-5::$P_{MLS1}$-tCrGES-$T_{ADH1}$, $P_{ADH2}$-ScERG20$^{WW}$-$T_{CYC1}$ ; XI-1:: $P_{PCK1}$-SctHMGR-$T_{ADH1}$, $P_{MLS1}$-ScIDI1-$T_{CYC1}$; $P_{ERG20\Delta}$::$P_{HXT1}$ ; XII-4::$P_{ADH2}$-CrTDC-$T_{ADH1}$, $P_{MLS1}$-ScZWF1-$T_{CYC1}$ ; **II-1:: $P_{ADH2}$-CrMSBP1-$T_{ADH1}$** | This study |
| BSY211 | ST7574, X-2::$P_{ADH2}$-CrCPR-$T_{ADH1}$, $P_{PCK1}$-CrCYB5-$T_{CYC1}$ ; XI-2::$P_{PCK1}$-CrIO-$T_{ADH1}$, $P_{ADH2}$-Cr7DLGT-$T_{CYC1}$ ; XII-2::$P_{ADH2}$-CrCYPADH-$T_{ADH1}$, $P_{ICL1}$Cr7DLH-$T_{CYC1}$ ; XI-3::$P_{ICL1}$-CrLAMT-$T_{ADH1}$, $P_{ADH2}$-CrSLS-$T_{CYC1}$ ; XI-5::$P_{ADH2}$-CrSTR-$T_{ADH1}$ ; X-3::$P_{PCK1}$-CrGOR-$T_{ADH1}$, $P_{ADH2}$-CrG8H-$T_{CYC1}$ ; ΔOYE2, ΔOYE3 ; ΔATF1, ΔARI1 ; ΔADH6 ; X-4::$P_{ADH2}$-NcISY2-$T_{ADH1}$, $P_{ICL1}$-NcMLPLa-$T_{CYC1}$ ; XII-5::$P_{MLS1}$-tCrGES-$T_{ADH1}$, $P_{ADH2}$-ScERG20$^{WW}$-$T_{CYC1}$ ; XI-1:: $P_{PCK1}$-SctHMGR-$T_{ADH1}$, $P_{MLS1}$-ScIDI1-$T_{CYC1}$; $P_{ERG20\Delta}$::$P_{HXT1}$ ; XII-4::$P_{ADH2}$-CrTDC-$T_{ADH1}$, $P_{MLS1}$-ScZWF1-$T_{CYC1}$ ; **II-1:: $P_{ADH2}$-CrMSBP2-$T_{ADH1}$** | This study |
| BSY212 | ST7574, X-2::$P_{ADH2}$-CrCPR-$T_{ADH1}$, $P_{PCK1}$-CrCYB5-$T_{CYC1}$ ; XI-2::$P_{PCK1}$-CrIO-$T_{ADH1}$, $P_{ADH2}$-Cr7DLGT-$T_{CYC1}$ ; XII-2::$P_{ADH2}$-CrCYPADH-$T_{ADH1}$, $P_{ICL1}$Cr7DLH-$T_{CYC1}$ ; XI-3::$P_{ICL1}$-CrLAMT-$T_{ADH1}$, $P_{ADH2}$-CrSLS-$T_{CYC1}$ ; XI-5::$P_{ADH2}$-CrSTR-$T_{ADH1}$ ; X-3::$P_{PCK1}$-CrGOR-$T_{ADH1}$, $P_{ADH2}$-CrG8H-$T_{CYC1}$ ; ΔOYE2, ΔOYE3 ; ΔATF1, ΔARI1 ; ΔADH6 ; X-4::$P_{ADH2}$-NcISY2-$T_{ADH1}$, $P_{ICL1}$-NcMLPLa-$T_{CYC1}$ ; XII-5::$P_{MLS1}$-tCrGES-$T_{ADH1}$, $P_{ADH2}$-ScERG20$^{WW}$-$T_{CYC1}$ ; XI-1:: $P_{PCK1}$-SctHMGR-$T_{ADH1}$, $P_{MLS1}$-ScIDI1-$T_{CYC1}$; $P_{ERG20\Delta}$::$P_{HXT1}$ ; XII-4::$P_{ADH2}$-CrTDC-$T_{ADH1}$, $P_{MLS1}$-ScZWF1-$T_{CYC1}$ ; **II-1:: $P_{ADH2}$-CaMSBP1-$T_{ADH1}$** | This study |
| BSY213 | ST7574, X-2::$P_{ADH2}$-CrCPR-$T_{ADH1}$, $P_{PCK1}$-CrCYB5-$T_{CYC1}$ ; XI-2::$P_{PCK1}$-CrIO-$T_{ADH1}$, $P_{ADH2}$-Cr7DLGT-$T_{CYC1}$ ; XII-2::$P_{ADH2}$-CrCYPADH-$T_{ADH1}$, $P_{ICL1}$Cr7DLH-$T_{CYC1}$ ; XI-3::$P_{ICL1}$-CrLAMT-$T_{ADH1}$, $P_{ADH2}$-CrSLS-$T_{CYC1}$ ; XI-5::$P_{ADH2}$-CrSTR-$T_{ADH1}$ ; X-3::$P_{PCK1}$-CrGOR-$T_{ADH1}$, $P_{ADH2}$-CrG8H-$T_{CYC1}$ ; ΔOYE2, ΔOYE3 ; ΔATF1, ΔARI1 ; ΔADH6 ; X-4::$P_{ADH2}$-NcISY2-$T_{ADH1}$, $P_{ICL1}$-NcMLPLa-$T_{CYC1}$ ; XII-5::$P_{MLS1}$-tCrGES-$T_{ADH1}$, $P_{ADH2}$-ScERG20$^{WW}$-$T_{CYC1}$ ; XI-1:: $P_{PCK1}$-SctHMGR-$T_{ADH1}$, $P_{MLS1}$-ScIDI1-$T_{CYC1}$; $P_{ERG20\Delta}$::$P_{HXT1}$ ; XII-4::$P_{ADH2}$-CrTDC-$T_{ADH1}$, $P_{MLS1}$-ScZWF1-$T_{CYC1}$ ; **II-1:: $P_{ADH2}$-CaMSBP2-$T_{ADH1}$** | This study |
| BSY214 | ST7574, X-2::$P_{ADH2}$-CrCPR-$T_{ADH1}$, $P_{PCK1}$-CrCYB5-$T_{CYC1}$ ; XI-2::$P_{PCK1}$-CrIO-$T_{ADH1}$, $P_{ADH2}$-Cr7DLGT-$T_{CYC1}$ ; XII-2::$P_{ADH2}$-CrCYPADH-$T_{ADH1}$, $P_{ICL1}$Cr7DLH-$T_{CYC1}$ ; XI-3::$P_{ICL1}$-CrLAMT-$T_{ADH1}$, $P_{ADH2}$-CrSLS-$T_{CYC1}$ ; XI-5::$P_{ADH2}$-CrSTR-$T_{ADH1}$ ; X-3::$P_{PCK1}$-CrGOR-$T_{ADH1}$, $P_{ADH2}$-CrG8H-$T_{CYC1}$ ; ΔOYE2, ΔOYE3 ; ΔATF1, ΔARI1 ; ΔADH6 ; X-4::$P_{ADH2}$-NcISY2-$T_{ADH1}$, $P_{ICL1}$-NcMLPLa-$T_{CYC1}$ ; XII-5::$P_{MLS1}$-tCrGES-$T_{ADH1}$, $P_{ADH2}$-ScERG20$^{WW}$-$T_{CYC1}$ ; XI-1:: $P_{PCK1}$-SctHMGR-$T_{ADH1}$, $P_{MLS1}$-ScIDI1-$T_{CYC1}$; $P_{ERG20\Delta}$::$P_{HXT1}$ ; XII-4::$P_{ADH2}$-CrTDC-$T_{ADH1}$, $P_{MLS1}$-ScZWF1-$T_{CYC1}$ ; **II-1:: $P_{ADH2}$-AaMSBP1-$T_{ADH1}$** | This study |
| BSY215 | ST7574, X-2::$P_{ADH2}$-CrCPR-$T_{ADH1}$, $P_{PCK1}$-CrCYB5-$T_{CYC1}$ ; XI-2::$P_{PCK1}$-CrIO-$T_{ADH1}$, $P_{ADH2}$-Cr7DLGT-$T_{CYC1}$ ; XII-2::$P_{ADH2}$-CrCYPADH-$T_{ADH1}$, $P_{ICL1}$Cr7DLH-$T_{CYC1}$ ; XI-3::$P_{ICL1}$-CrLAMT-$T_{ADH1}$, $P_{ADH2}$-CrSLS-$T_{CYC1}$ ; XI-5::$P_{ADH2}$-CrSTR-$T_{ADH1}$ ; X-3::$P_{PCK1}$-CrGOR-$T_{ADH1}$, $P_{ADH2}$-CrG8H-$T_{CYC1}$ ; ΔOYE2, ΔOYE3 ; ΔATF1, ΔARI1 ; ΔADH6 ; X-4::$P_{ADH2}$-NcISY2-$T_{ADH1}$, $P_{ICL1}$-NcMLPLa-$T_{CYC1}$ ; XII-5::$P_{MLS1}$-tCrGES-$T_{ADH1}$, $P_{ADH2}$-ScERG20$^{WW}$-$T_{CYC1}$ ; XI-1:: $P_{PCK1}$-SctHMGR-$T_{ADH1}$, $P_{MLS1}$-ScIDI1-$T_{CYC1}$; $P_{ERG20\Delta}$::$P_{HXT1}$ ; XII-4::$P_{ADH2}$-CrTDC-$T_{ADH1}$, $P_{MLS1}$-ScZWF1-$T_{CYC1}$ ; **II-1:: $P_{ADH2}$-CsMSBP1-$T_{ADH1}$** | This study |
| BSY216 | ST7574, X-2::$P_{ADH2}$-CrCPR-$T_{ADH1}$, $P_{PCK1}$-CrCYB5-$T_{CYC1}$ ; XI-2::$P_{PCK1}$-CrIO-$T_{ADH1}$, $P_{ADH2}$-Cr7DLGT-$T_{CYC1}$ ; XII-2::$P_{ADH2}$-CrCYPADH-$T_{ADH1}$, $P_{ICL1}$Cr7DLH-$T_{CYC1}$ ; XI-3::$P_{ICL1}$-CrLAMT-$T_{ADH1}$, $P_{ADH2}$-CrSLS-$T_{CYC1}$ ; XI-5::$P_{ADH2}$-CrSTR-$T_{ADH1}$ ; X-3::$P_{PCK1}$-CrGOR-$T_{ADH1}$, $P_{ADH2}$-CrG8H-$T_{CYC1}$ ; ΔOYE2, ΔOYE3 ; ΔATF1, ΔARI1 ; ΔADH6 ; X-4::$P_{ADH2}$-NcISY2-$T_{ADH1}$, $P_{ICL1}$-NcMLPLa-$T_{CYC1}$ ; XII-5::$P_{MLS1}$-tCrGES-$T_{ADH1}$, $P_{ADH2}$-ScERG20$^{WW}$-$T_{CYC1}$ ; XI-1:: $P_{PCK1}$-SctHMGR-$T_{ADH1}$, $P_{MLS1}$-ScIDI1-$T_{CYC1}$; $P_{ERG20\Delta}$::$P_{HXT1}$ ; XII-4::$P_{ADH2}$-CrTDC-$T_{ADH1}$, $P_{MLS1}$-ScZWF1-$T_{CYC1}$ ; **II-1:: $P_{ADH2}$-CsMSBP2-$T_{ADH1}$** | This study |
| BSY217 | ST7574, X-2::$P_{ADH2}$-CrCPR-$T_{ADH1}$, $P_{PCK1}$-CrCYB5-$T_{CYC1}$ ; XI-2::$P_{PCK1}$-CrIO-$T_{ADH1}$, $P_{ADH2}$-Cr7DLGT-$T_{CYC1}$ ; XII-2::$P_{ADH2}$-CrCYPADH-$T_{ADH1}$, $P_{ICL1}$Cr7DLH-$T_{CYC1}$ ; XI-3::$P_{ICL1}$-CrLAMT-$T_{ADH1}$, $P_{ADH2}$-CrSLS-$T_{CYC1}$ ; XI-5::$P_{ADH2}$-CrSTR-$T_{ADH1}$ ; X-3::$P_{PCK1}$-CrGOR-$T_{ADH1}$, $P_{ADH2}$-CrG8H-$T_{CYC1}$ ; ΔOYE2, ΔOYE3 ; ΔATF1, ΔARI1 ; ΔADH6 ; X-4::$P_{ADH2}$-NcISY2-$T_{ADH1}$, $P_{ICL1}$-NcMLPLa-$T_{CYC1}$ ; XII-5::$P_{MLS1}$-tCrGES-$T_{ADH1}$, $P_{ADH2}$-ScERG20$^{WW}$-$T_{CYC1}$ ; XI-1:: $P_{PCK1}$-SctHMGR-$T_{ADH1}$, $P_{MLS1}$-ScIDI1-$T_{CYC1}$; $P_{ERG20\Delta}$::$P_{HXT1}$ ; XII-4::$P_{ADH2}$-CrTDC-$T_{ADH1}$, $P_{MLS1}$-ScZWF1-$T_{CYC1}$ ; **II-1:: $P_{ADH2}$-ScDAP1-$T_{ADH1}$** | This study |
| BSY221 | ST7574, X-2::$P_{ADH2}$-CrCPR-$T_{ADH1}$, $P_{PCK1}$-CrCYB5-$T_{CYC1}$ ; XI-2::$P_{PCK1}$-CrIO-$T_{ADH1}$, $P_{ADH2}$-Cr7DLGT-$T_{CYC1}$ ; XII-2::$P_{ADH2}$-CrCYPADH-$T_{ADH1}$, $P_{ICL1}$Cr7DLH-$T_{CYC1}$ ; XI-3::$P_{ICL1}$-CrLAMT-$T_{ADH1}$, $P_{ADH2}$-CrSLS-$T_{CYC1}$ ; XI-5::$P_{ADH2}$-CrSTR-$T_{ADH1}$ ; X-3::$P_{PCK1}$-CrGOR-$T_{ADH1}$, $P_{ADH2}$-CrG8H-$T_{CYC1}$ ; ΔOYE2, ΔOYE3 ; ΔATF1, ΔARI1 ; ΔADH6 ; X-4::$P_{ADH2}$-NcISY2-$T_{ADH1}$, $P_{ICL1}$-NcMLPLa-$T_{CYC1}$ ;XI-1:: $P_{PCK1}$-SctHMGR-$T_{ADH1}$, $P_{MLS1}$-ScIDI1-$T_{CYC1}$; $P_{ERG20\Delta}$::$P_{HXT1}$ ; XII-4::$P_{ADH2}$-CrTDC-$T_{ADH1}$, $P_{MLS1}$ScZWF1-$T_{CYC1}$ ; II-1:: $P_{ADH2}$-CaMSBP1-$T_{ADH1}$<br><br>Note: Strain was generated by deletion of XII-5::$P_{MLS1}$-tCrGES-$T_{ADH1}$, $P_{ADH2}$-ScERG20$^{WW}$-$T_{CYC1}$ from strain BSY212 | This study |
| BSY225 | ST7574, X-2::$P_{ADH2}$-CrCPR-$T_{ADH1}$, $P_{PCK1}$-CrCYB5-$T_{CYC1}$ ; XI-2::$P_{PCK1}$-CrIO-$T_{ADH1}$, $P_{ADH2}$-Cr7DLGT-$T_{CYC1}$ ; XII-2::$P_{ADH2}$-CrCYPADH-$T_{ADH1}$, $P_{ICL1}$Cr7DLH-$T_{CYC1}$ ; XI-3::$P_{ICL1}$- | This study |

| | | |
|---|---|---|
| | CrLAMT-$T_{ADH1}$, $P_{ADH2}$-CrSLS-$T_{CYC1}$ ; XI-5::$P_{ADH2}$-CrSTR-$T_{ADH1}$ ; X-3::$P_{PCK1}$-CrGOR-$T_{ADH1}$, $P_{ADH2}$-CrG8H-$T_{CYC1}$ ; ΔOYE2, ΔOYE3 ; ΔATF1, ΔARI1 ; ΔADH6 ; X-4::$P_{ADH2}$-NcISY2-$T_{ADH1}$, $P_{ICL1}$-NcMLPLa-$T_{CYC1}$ ; **XII-5::$P_{MLS1}$-tCrGES-$T_{ADH1}$, $P_{ADH2}$-ScERG20$^{WWG}$-$T_{CYC1}$** ; XI-1:: $P_{PCK1}$-SctHMGR-$T_{ADH1}$, $P_{MLS1}$-ScIDI1-$T_{CYC1}$; $P_{ERG20\Delta}$::$P_{HXT1}$ ; XII-4::$P_{ADH2}$-CrTDC-$T_{ADH1}$, $P_{MLS1}$-ScZWF1-$T_{CYC1}$ ; II-1:: $P_{ADH2}$-CaMSBP1-$T_{ADH1}$ | |
| BSY238 | ST7574, X-2::$P_{ADH2}$-CrCPR-$T_{ADH1}$, $P_{PCK1}$-CrCYB5-$T_{CYC1}$ ; XI-2::$P_{PCK1}$-CrIO-$T_{ADH1}$, $P_{ADH2}$-Cr7DLGT-$T_{CYC1}$ ; XII-2::$P_{ADH2}$-CrCYPADH-$T_{ADH1}$, $P_{ICL1}$Cr7DLH-$T_{CYC1}$ ; XI-3::$P_{ICL1}$-CrLAMT-$T_{ADH1}$, $P_{ADH2}$-CrSLS-$T_{CYC1}$ ; XI-5::$P_{ADH2}$-CrSTR-$T_{ADH1}$ ; X-3::$P_{PCK1}$-CrGOR-$T_{ADH1}$, $P_{ADH2}$-CrG8H-$T_{CYC1}$ ; ΔOYE2, ΔOYE3 ; ΔATF1, ΔARI1 ; ΔADH6 ; X-4::$P_{ADH2}$-NcISY2-$T_{ADH1}$, $P_{ICL1}$-NcMLPLa-$T_{CYC1}$ ; XII-5::$P_{MLS1}$-tCrGES-$T_{ADH1}$, $P_{ADH2}$-ScERG20$^{WW}$-$T_{CYC1}$ ; XI-1:: $P_{PCK1}$-SctHMGR-$T_{ADH1}$, $P_{MLS1}$-ScIDI1-$T_{CYC1}$; $P_{ERG20\Delta}$::$P_{HXT1}$ ; XII-4::$P_{ADH2}$-CrTDC-$T_{ADH1}$, $P_{MLS1}$-ScZWF1-$T_{CYC1}$ ; II-1:: $P_{ADH2}$-CaMSBP1-$T_{ADH1}$ ; **VII-1::$P_{MLS1}$-tCrGES-$T_{ADH1}$, $P_{ADH2}$-ScERG20$^{WW}$-$T_{CYC1}$** | This study |
| BSY239 | ST7574, X-2::$P_{ADH2}$-CrCPR-$T_{ADH1}$, $P_{PCK1}$-CrCYB5-$T_{CYC1}$ ; XI-2::$P_{PCK1}$-CrIO-$T_{ADH1}$, $P_{ADH2}$-Cr7DLGT-$T_{CYC1}$ ; XII-2::$P_{ADH2}$-CrCYPADH-$T_{ADH1}$, $P_{ICL1}$Cr7DLH-$T_{CYC1}$ ; XI-3::$P_{ICL1}$-CrLAMT-$T_{ADH1}$, $P_{ADH2}$-CrSLS-$T_{CYC1}$ ; XI-5::$P_{ADH2}$-CrSTR-$T_{ADH1}$ ; X-3::$P_{PCK1}$-CrGOR-$T_{ADH1}$, $P_{ADH2}$-CrG8H-$T_{CYC1}$ ; ΔOYE2, ΔOYE3 ; ΔATF1, ΔARI1 ; ΔADH6 ; X-4::$P_{ADH2}$-NcISY2-$T_{ADH1}$, $P_{ICL1}$-NcMLPLa-$T_{CYC1}$ ; XII-5::$P_{MLS1}$-tCrGES-$T_{ADH1}$, $P_{ADH2}$-ScERG20$^{WW}$-$T_{CYC1}$ ; XI-1:: $P_{PCK1}$-SctHMGR-$T_{ADH1}$, $P_{MLS1}$-ScIDI1-$T_{CYC1}$; $P_{ERG20\Delta}$::$P_{HXT1}$ ; XII-4::$P_{ADH2}$-CrTDC-$T_{ADH1}$, $P_{MLS1}$-ScZWF1-$T_{CYC1}$ ; II-1:: $P_{ADH2}$-CaMSBP1-$T_{ADH1}$ ; **VII-1::$P_{MLS1}$-tCrGES-$T_{ADH1}$, $P_{ADH2}$-ScERG20$^{WWG}$-$T_{CYC1}$** | This study |
| BSY240 | ST7574, X-2::$P_{ADH2}$-CrCPR-$T_{ADH1}$, $P_{PCK1}$-CrCYB5-$T_{CYC1}$ ; XI-2::$P_{PCK1}$-CrIO-$T_{ADH1}$, $P_{ADH2}$-Cr7DLGT-$T_{CYC1}$ ; XII-2::$P_{ADH2}$-CrCYPADH-$T_{ADH1}$, $P_{ICL1}$Cr7DLH-$T_{CYC1}$ ; XI-3::$P_{ICL1}$-CrLAMT-$T_{ADH1}$, $P_{ADH2}$-CrSLS-$T_{CYC1}$ ; XI-5::$P_{ADH2}$-CrSTR-$T_{ADH1}$ ; X-3::$P_{PCK1}$-CrGOR-$T_{ADH1}$, $P_{ADH2}$-CrG8H-$T_{CYC1}$ ; ΔOYE2, ΔOYE3 ; ΔATF1, ΔARI1 ; ΔADH6 ; X-4::$P_{ADH2}$-NcISY2-$T_{ADH1}$, $P_{ICL1}$-NcMLPLa-$T_{CYC1}$ ; XII-5::$P_{MLS1}$-tCrGES-$T_{ADH1}$, $P_{ADH2}$-ScERG20$^{WWG}$-$T_{CYC1}$ ; XI-1:: $P_{PCK1}$-SctHMGR-$T_{ADH1}$, $P_{MLS1}$-ScIDI1-$T_{CYC1}$; $P_{ERG20\Delta}$::$P_{HXT1}$ ; XII-4::$P_{ADH2}$-CrTDC-$T_{ADH1}$, $P_{MLS1}$-ScZWF1-$T_{CYC1}$ ; II-1:: $P_{ADH2}$-CaMSBP1-$T_{ADH1}$ ; **VII-1::$P_{MLS1}$-tCrGES-$T_{ADH1}$, $P_{ADH2}$-ScERG20$^{WWG}$-$T_{CYC1}$** | This study |
| BSY263 | ST7574, X-2::$P_{ADH2}$-CrCPR-$T_{ADH1}$, $P_{PCK1}$-CrCYB5-$T_{CYC1}$ ; XI-2::$P_{PCK1}$-CrIO-$T_{ADH1}$, $P_{ADH2}$-Cr7DLGT-$T_{CYC1}$ ; XII-2::$P_{ADH2}$-CrCYPADH-$T_{ADH1}$, $P_{ICL1}$Cr7DLH-$T_{CYC1}$ ; XI-3::$P_{ICL1}$-CrLAMT-$T_{ADH1}$, $P_{ADH2}$-CrSLS-$T_{CYC1}$ ; XI-5::$P_{ADH2}$-CrSTR-$T_{ADH1}$ ; ΔOYE2, ΔOYE3 ; ΔATF1, ΔARI1 ; ΔADH6 ; X-4::$P_{ADH2}$-NcISY2-$T_{ADH1}$, $P_{ICL1}$-NcMLPLa-$T_{CYC1}$ ;XI-1:: $P_{PCK1}$-SctHMGR-$T_{ADH1}$, $P_{MLS1}$-ScIDI1-$T_{CYC1}$; $P_{ERG20\Delta}$::$P_{HXT1}$ ; XII-4::$P_{ADH2}$-CrTDC-$T_{ADH1}$, $P_{MLS1}$ScZWF1-$T_{CYC1}$ ; II-1:: $P_{ADH2}$-CaMSBP1-$T_{ADH1}$<br><br>Note: Strain was generated by deletion of X-3::$P_{PCK1}$-CrGOR-$T_{ADH1}$, $P_{ADH2}$-CrG8H-$T_{CYC1}$ from strain BSY221 | This study |
| BSY264 | ST7574, X-2::$P_{ADH2}$-CrCPR-$T_{ADH1}$, $P_{PCK1}$-CrCYB5-$T_{CYC1}$ ; XI-2::$P_{PCK1}$-CrIO-$T_{ADH1}$, $P_{ADH2}$-Cr7DLGT-$T_{CYC1}$ ; XII-2::$P_{ADH2}$-CrCYPADH-$T_{ADH1}$, $P_{ICL1}$Cr7DLH-$T_{CYC1}$ ; XI-3::$P_{ICL1}$-CrLAMT-$T_{ADH1}$, $P_{ADH2}$-CrSLS-$T_{CYC1}$ ; XI-5::$P_{ADH2}$-CrSTR-$T_{ADH1}$ ; X-3::$P_{PCK1}$-CrGOR-$T_{ADH1}$, $P_{ADH2}$-CrG8H-$T_{CYC1}$ ; ΔOYE2, ΔOYE3 ; ΔATF1, ΔARI1 ; ΔADH6 ; X-4::$P_{ADH2}$-NcISY2-$T_{ADH1}$, $P_{ICL1}$-NcMLPLa-$T_{CYC1}$ ;XI-1:: $P_{PCK1}$-SctHMGR-$T_{ADH1}$, $P_{MLS1}$-ScIDI1-$T_{CYC1}$; $P_{ERG20\Delta}$::$P_{HXT1}$ ; XII-4::$P_{ADH2}$-CrTDC-$T_{ADH1}$, $P_{MLS1}$ScZWF1-$T_{CYC1}$ ; II-1:: $P_{ADH2}$-CaMSBP1-$T_{ADH1}$<br><br>Note: Strain was generated by deletion of X-3::$P_{ADH2}$-CrG8H-$T_{CYC1}$ from strain BSY221 | This study |
| BSY265 | ST7574, X-2::$P_{ADH2}$-CrCPR-$T_{ADH1}$, $P_{PCK1}$-CrCYB5-$T_{CYC1}$ ; XI-2::$P_{PCK1}$-CrIO-$T_{ADH1}$, $P_{ADH2}$-Cr7DLGT-$T_{CYC1}$ ; XII-2::$P_{ADH2}$-CrCYPADH-$T_{ADH1}$, $P_{ICL1}$Cr7DLH-$T_{CYC1}$ ; XI-3::$P_{ADH2}$-CrSLS-$T_{CYC1}$; XI-5::$P_{ADH2}$-CrSTR-$T_{ADH1}$ ; X-3::$P_{PCK1}$-CrGOR-$T_{ADH1}$, $P_{ADH2}$-CrG8H-$T_{CYC1}$ ; ΔOYE2, ΔOYE3 ; ΔATF1, ΔARI1 ; ΔADH6 ; X-4::$P_{ADH2}$-NcISY2-$T_{ADH1}$, $P_{ICL1}$-NcMLPLa-$T_{CYC1}$ ; XII-5::$P_{MLS1}$-tCrGES-$T_{ADH1}$, $P_{ADH2}$-ScERG20$^{WW}$-$T_{CYC1}$ ; XI-1:: $P_{PCK1}$-SctHMGR-$T_{ADH1}$, $P_{MLS1}$-ScIDI1-$T_{CYC1}$; $P_{ERG20\Delta}$::$P_{HXT1}$ ; XII-4::$P_{ADH2}$-CrTDC-$T_{ADH1}$, $P_{MLS1}$-ScZWF1-$T_{CYC1}$ ; II-1:: $P_{ADH2}$-CaMSBP1-$T_{ADH1}$ ; VII-1::$P_{MLS1}$-tCrGES-$T_{ADH1}$, $P_{ADH2}$-ScERG20$^{WW}$-$T_{CYC1}$<br><br>Note: Strain was generated by deletion of XI-3::$P_{ICL1}$-CrLAMT-$T_{ADH1}$ from strain BSY238 | This study |
| BSY266 | ST7574, X-2::$P_{ADH2}$-CrCPR-$T_{ADH1}$, $P_{PCK1}$-CrCYB5-$T_{CYC1}$ ; XI-2::$P_{PCK1}$-CrIO-$T_{ADH1}$, $P_{ADH2}$-Cr7DLGT-$T_{CYC1}$ ; XII-2::$P_{ADH2}$-CrCYPADH-$T_{ADH1}$, $P_{ICL1}$Cr7DLH-$T_{CYC1}$ ; ; XI-5::$P_{ADH2}$-CrSTR-$T_{ADH1}$ ; X-3::$P_{PCK1}$-CrGOR-$T_{ADH1}$, $P_{ADH2}$-CrG8H-$T_{CYC1}$ ; ΔOYE2, ΔOYE3 ; ΔATF1, ΔARI1 ; ΔADH6 ; X-4::$P_{ADH2}$-NcISY2-$T_{ADH1}$, $P_{ICL1}$-NcMLPLa-$T_{CYC1}$ ; XII-5::$P_{MLS1}$-tCrGES-$T_{ADH1}$, $P_{ADH2}$-ScERG20$^{WW}$-$T_{CYC1}$ ; XI-1:: $P_{PCK1}$-SctHMGR-$T_{ADH1}$, $P_{MLS1}$-ScIDI1-$T_{CYC1}$; $P_{ERG20\Delta}$::$P_{HXT1}$ ; XII-4::$P_{ADH2}$-CrTDC-$T_{ADH1}$, $P_{MLS1}$-ScZWF1-$T_{CYC1}$ ; II-1:: $P_{ADH2}$-CaMSBP1-$T_{ADH1}$ ; VII-1::$P_{MLS1}$-tCrGES-$T_{ADH1}$, $P_{ADH2}$-ScERG20$^{WW}$-$T_{CYC1}$<br><br>Note: Strain was generated by deletion of XI-3::$P_{ICL1}$-CrLAMT-$T_{ADH1}$, $P_{ADH2}$-CrSLS-$T_{CYC1}$ from strain BSY238 | This study |
| BSY279 | ST7574, X-2::$P_{ADH2}$-CrCPR-$T_{ADH1}$, $P_{PCK1}$-CrCYB5-$T_{CYC1}$ ; XI-2::$P_{PCK1}$-CrIO-$T_{ADH1}$, $P_{ADH2}$-Cr7DLGT-$T_{CYC1}$ ; XII-2::$P_{ADH2}$-CrCYPADH-$T_{ADH1}$, $P_{ICL1}$Cr7DLH-$T_{CYC1}$ ; **XI-3::$P_{ICL1}$-** | This study |

| | | |
|---|---|---|
| | **CaCYP72A565$^{opt}$-$T_{ADH1}$, $P_{ADH2}$-CaCYP72A610$^{opt}$-$T_{CYC1}$** ; XI-5::$P_{ADH2}$-CrSTR-$T_{ADH1}$ ; X-3::$P_{PCK1}$-CrGOR-$T_{ADH1}$, $P_{ADH2}$-CrG8H-$T_{CYC1}$ ; ΔOYE2, ΔOYE3 ; ΔATF1, ΔARI1 ; ΔADH6 ; X-4::$P_{ADH2}$-NcISY2-$T_{ADH1}$, $P_{ICL1}$-NcMLPLa-$T_{CYC1}$ ; XII-5::$P_{MLS1}$-tCrGES-$T_{ADH1}$, $P_{ADH2}$-ScERG20$^{WW}$-$T_{CYC1}$ ; XI-1:: $P_{PCK1}$-SctHMGR-$T_{ADH1}$, $P_{MLS1}$-ScIDI1-$T_{CYC1}$; $P_{ERG20\Delta}$::$P_{HXT1}$ ; XII-4::$P_{ADH2}$-CrTDC-$T_{ADH1}$, $P_{MLS1}$-ScZWF1-$T_{CYC1}$ ; II-1:: $P_{ADH2}$-CaMSBP1-$T_{ADH1}$ ; VII-1::$P_{MLS1}$-tCrGES-$T_{ADH1}$, $P_{ADH2}$-ScERG20$^{WW}$-$T_{CYC1}$ | |
| BSY280 | ST7574, X-2::$P_{ADH2}$-CrCPR-$T_{ADH1}$, $P_{PCK1}$-CrCYB5-$T_{CYC1}$ ; XI-2::$P_{PCK1}$-CrIO-$T_{ADH1}$, $P_{ADH2}$-Cr7DLGT-$T_{CYC1}$ ; XII-2::$P_{ADH2}$-CrCYPADH-$T_{ADH1}$, $P_{ICL1}$Cr7DLH-$T_{CYC1}$ ; **XI-3::$P_{ADH2}$-CaCYP72A565$^{opt}$-$T_{ADH1}$**; XI-5::$P_{ADH2}$-CrSTR-$T_{ADH1}$ ; X-3::$P_{PCK1}$-CrGOR-$T_{ADH1}$, $P_{ADH2}$-CrG8H-$T_{CYC1}$ ; ΔOYE2, ΔOYE3 ; ΔATF1, ΔARI1 ; ΔADH6 ; X-4::$P_{ADH2}$-NcISY2-$T_{ADH1}$, $P_{ICL1}$-NcMLPLa-$T_{CYC1}$ ; XII-5::$P_{MLS1}$-tCrGES-$T_{ADH1}$, $P_{ADH2}$-ScERG20$^{WW}$-$T_{CYC1}$ ;XI-1:: $P_{PCK1}$-SctHMGR-$T_{ADH1}$, $P_{MLS1}$-ScIDI1-$T_{CYC1}$; $P_{ERG20\Delta}$::$P_{HXT1}$ ; XII-4::$P_{ADH2}$-CrTDC-$T_{ADH1}$, $P_{MLS1}$-ScZWF1-$T_{CYC1}$ ; II-1:: $P_{ADH2}$-CaMSBP1-$T_{ADH1}$ ; VII-1::$P_{MLS1}$-tCrGES-$T_{ADH1}$, $P_{ADH2}$-ScERG20$^{WW}$-$T_{CYC1}$ | This study |
| BSY281 | ST7574, X-2::$P_{ADH2}$-CrCPR-$T_{ADH1}$, $P_{PCK1}$-CrCYB5-$T_{CYC1}$ ; XI-2::$P_{PCK1}$-CrIO-$T_{ADH1}$, $P_{ADH2}$-Cr7DLGT-$T_{CYC1}$ ; XII-2::$P_{ADH2}$-CrCYPADH-$T_{ADH1}$, $P_{ICL1}$Cr7DLH-$T_{CYC1}$ ; **XI-3::$P_{ADH2}$-CaCYP72A610$^{opt}$-$T_{ADH1}$**; XI-5::$P_{ADH2}$-CrSTR-$T_{ADH1}$ ; X-3::$P_{PCK1}$-CrGOR-$T_{ADH1}$, $P_{ADH2}$-CrG8H-$T_{CYC1}$ ; ΔOYE2, ΔOYE3 ; ΔATF1, ΔARI1 ; ΔADH6 ; X-4::$P_{ADH2}$-NcISY2-$T_{ADH1}$, $P_{ICL1}$-NcMLPLa-$T_{CYC1}$ ; XII-5::$P_{MLS1}$-tCrGES-$T_{ADH1}$, $P_{ADH2}$-ScERG20$^{WW}$-$T_{CYC1}$ ;XI-1:: $P_{PCK1}$-SctHMGR-$T_{ADH1}$, $P_{MLS1}$-ScIDI1-$T_{CYC1}$; $P_{ERG20\Delta}$::$P_{HXT1}$ ; XII-4::$P_{ADH2}$-CrTDC-$T_{ADH1}$, $P_{MLS1}$-ScZWF1-$T_{CYC1}$ ; II-1:: $P_{ADH2}$-CaMSBP1-$T_{ADH1}$ ; VII-1::$P_{MLS1}$-tCrGES-$T_{ADH1}$, $P_{ADH2}$-ScERG20$^{WW}$-$T_{CYC1}$ | This study |
| BSY288 | ST7574, X-2::$P_{ADH2}$-CrCPR-$T_{ADH1}$, $P_{PCK1}$-CrCYB5-$T_{CYC1}$ ; XI-2::$P_{PCK1}$-CrIO-$T_{ADH1}$, $P_{ADH2}$-Cr7DLGT-$T_{CYC1}$ ; XII-2::$P_{ADH2}$-CrCYPADH-$T_{ADH1}$, $P_{ICL1}$Cr7DLH-$T_{CYC1}$ ; XI-3::$P_{ICL1}$-CrLAMT-$T_{ADH1}$, $P_{ADH2}$-CrSLS-$T_{CYC1}$ ; XI-5::$P_{ADH2}$-CrSTR-$T_{ADH1}$ ; X-3::$P_{PCK1}$-CrGOR-$T_{ADH1}$, $P_{ADH2}$-CrG8H-$T_{CYC1}$ ; ΔOYE2, ΔOYE3 ; ΔATF1, ΔARI1 ; ΔADH6 ; X-4::$P_{ADH2}$-NcISY2-$T_{ADH1}$, $P_{ICL1}$-NcMLPLa-$T_{CYC1}$ ; XII-5::$P_{MLS1}$-tCrGES-$T_{ADH1}$, $P_{ADH2}$-ScERG20$^{WW}$-$T_{CYC1}$ ; XI-1:: $P_{PCK1}$-SctHMGR-$T_{ADH1}$, $P_{MLS1}$-ScIDI1-$T_{CYC1}$; $P_{ERG20\Delta}$::$P_{HXT1}$ ; XII-4::$P_{ADH2}$-CrTDC-$T_{ADH1}$, $P_{MLS1}$-ScZWF1-$T_{CYC1}$ ; II-1:: $P_{ADH2}$-CaMSBP1-$T_{ADH1}$ ; VII-1::$P_{MLS1}$-tCrGES-$T_{ADH1}$, $P_{ADH2}$-ScERG20$^{WW}$-$T_{CYC1}$; **ΔADH3** | This study |
| BSY289 | ST7574, X-2::$P_{ADH2}$-CrCPR-$T_{ADH1}$, $P_{PCK1}$-CrCYB5-$T_{CYC1}$ ; XI-2::$P_{PCK1}$-CrIO-$T_{ADH1}$, $P_{ADH2}$-Cr7DLGT-$T_{CYC1}$ ; XII-2::$P_{ADH2}$-CrCYPADH-$T_{ADH1}$, $P_{ICL1}$Cr7DLH-$T_{CYC1}$ ; XI-3::$P_{ICL1}$-CrLAMT-$T_{ADH1}$, $P_{ADH2}$-CrSLS-$T_{CYC1}$ ; XI-5::$P_{ADH2}$-CrSTR-$T_{ADH1}$ ; X-3::$P_{PCK1}$-CrGOR-$T_{ADH1}$, $P_{ADH2}$-CrG8H-$T_{CYC1}$ ; ΔOYE2, ΔOYE3 ; ΔATF1, ΔARI1 ; ΔADH6 ; X-4::$P_{ADH2}$-NcISY2-$T_{ADH1}$, $P_{ICL1}$-NcMLPLa-$T_{CYC1}$ ; XII-5::$P_{MLS1}$-tCrGES-$T_{ADH1}$, $P_{ADH2}$-ScERG20$^{WW}$-$T_{CYC1}$ ; XI-1:: $P_{PCK1}$-SctHMGR-$T_{ADH1}$, $P_{MLS1}$-ScIDI1-$T_{CYC1}$; $P_{ERG20\Delta}$::$P_{HXT1}$ ; XII-4::$P_{ADH2}$-CrTDC-$T_{ADH1}$, $P_{MLS1}$-ScZWF1-$T_{CYC1}$ ; II-1:: $P_{ADH2}$-CaMSBP1-$T_{ADH1}$ ; VII-1::$P_{MLS1}$-tCrGES-$T_{ADH1}$, $P_{ADH2}$-ScERG20$^{WW}$-$T_{CYC1}$; **ΔADH6** | This study |
| BSY290 | ST7574, X-2::$P_{ADH2}$-CrCPR-$T_{ADH1}$, $P_{PCK1}$-CrCYB5-$T_{CYC1}$ ; XI-2::$P_{PCK1}$-CrIO-$T_{ADH1}$, $P_{ADH2}$-Cr7DLGT-$T_{CYC1}$ ; XII-2::$P_{ADH2}$-CrCYPADH-$T_{ADH1}$, $P_{ICL1}$Cr7DLH-$T_{CYC1}$ ; XI-3::$P_{ICL1}$-CrLAMT-$T_{ADH1}$, $P_{ADH2}$-CrSLS-$T_{CYC1}$ ; XI-5::$P_{ADH2}$-CrSTR-$T_{ADH1}$ ; X-3::$P_{PCK1}$-CrGOR-$T_{ADH1}$, $P_{ADH2}$-CrG8H-$T_{CYC1}$ ; ΔOYE2, ΔOYE3 ; ΔATF1, ΔARI1 ; ΔADH6 ; X-4::$P_{ADH2}$-NcISY2-$T_{ADH1}$, $P_{ICL1}$-NcMLPLa-$T_{CYC1}$ ; XII-5::$P_{MLS1}$-tCrGES-$T_{ADH1}$, $P_{ADH2}$-ScERG20$^{WW}$-$T_{CYC1}$ ; XI-1:: $P_{PCK1}$-SctHMGR-$T_{ADH1}$, $P_{MLS1}$-ScIDI1-$T_{CYC1}$; $P_{ERG20\Delta}$::$P_{HXT1}$ ; XII-4::$P_{ADH2}$-CrTDC-$T_{ADH1}$, $P_{MLS1}$-ScZWF1-$T_{CYC1}$ ; II-1:: $P_{ADH2}$-CaMSBP1-$T_{ADH1}$ ; VII-1::$P_{MLS1}$-tCrGES-$T_{ADH1}$, $P_{ADH2}$-ScERG20$^{WW}$-$T_{CYC1}$; **ΔYPL062W** | This study |
| BSY294 | ST7574, X-2::$P_{ADH2}$-CrCPR-$T_{ADH1}$, $P_{PCK1}$-CrCYB5-$T_{CYC1}$ ; XI-2::$P_{PCK1}$-CrIO-$T_{ADH1}$, $P_{ADH2}$-Cr7DLGT-$T_{CYC1}$ ; XII-2::$P_{ADH2}$-CrCYPADH-$T_{ADH1}$ ; XI-3::$P_{ICL1}$-CaCYP72A565$^{opt}$-$T_{ADH1}$, $P_{ADH2}$-CaCYP72A610$^{opt}$-$T_{CYC1}$ ; XI-5::$P_{ADH2}$-CrSTR-$T_{ADH1}$ ; X-3::$P_{PCK1}$-CrGOR-$T_{ADH1}$, $P_{ADH2}$-CrG8H-$T_{CYC1}$ ; OYE2, ΔOYE3 ; ΔATF1, ΔARI1 ; ΔADH6 ; X-4::$P_{ADH2}$-NcISY2-$T_{ADH1}$, $P_{ICL1}$-NcMLPLa-$T_{CYC1}$ ; XII-5::$P_{MLS1}$-tCrGES-$T_{ADH1}$, $P_{ADH2}$-ScERG20$^{WW}$-$T_{CYC1}$ ; XI-1:: $P_{PCK1}$-SctHMGR-$T_{ADH1}$, $P_{MLS1}$-ScIDI1-$T_{CYC1}$; $P_{ERG20\Delta}$::$P_{HXT1}$ ; XII-4::$P_{ADH2}$-CrTDC-$T_{ADH1}$, $P_{MLS1}$-ScZWF1-$T_{CYC1}$ ; II-1:: $P_{ADH2}$-CaMSBP1-$T_{ADH1}$ ; VII-1::$P_{MLS1}$-tCrGES-$T_{ADH1}$, $P_{ADH2}$-ScERG20$^{WW}$-$T_{CYC1}$<br><br>Note: Strain was generated by deletion of XII-2::$P_{ICL1}$Cr7DLH-$T_{CYC1}$ from strain BSY279 | This study |

**Supplementary Table 2. List of genes used in this study.**

Nucleotide sequences that were amplified and differed from the deposited sequences are provided; differing positions are highlighted in red.

Codon-optimised sequences and nucleotide sequences not available on GenBank are also provided.

| Gene product | Origin | Short name | GenBank or Reference |
|---|---|---|---|
| NADPH-cytochrome P450 reductase | *Catharanthus roseus* | *Cr*CPR | X69791.1 |
| NADPH-cytochrome P450 reductase (amplified sequence) | ATGGATTCTAGCTCGGAGAAGTTGTCGCCGTTCGAATTGATGAGCGCGATCTTGAAGG<br>GAGCTAAATTAGATGGGTCTAACTCTTCAGATTCTGGCGTAGCTGTGTCGCCGGCAGT<br>TATGGCTATGTTGTTGGAGAATAAGGAGTTAGTGATGATTTTGACTACTTCAGTGGCG<br>GTTTTGATCGGTTGTGTCGTAGTTTTGATATGGCGGCGATCTTCCGGATCGGGTAAAA<br>AAGTCGTGGAGCCTCCGAAGCTCATAGTGCCTAAATCTGTTGTAGAACCGGAGGAAAT<br>TGATGAAGGGAAGAAGAAATTTACCATATTTTTTGGAACACAAACTGGAACAGCTGAA<br>GGCTTCGCTAAGGCTCTAGCTGAGGAAGCCAAAGCTCGATATGAAAAGGCAGTTATCA<br>AAGTGATTGATATAGATGATTATGCGGCTGATGATGAAGAATACGAGGAGAAATTCAG<br>AAAAGAGACCTTGGCATTTTTCATCTTGGCCACGTATGGAGATGGTGAGCCAACCGAC<br>AATGCTGCAAGGTTCTACAAATGGTTTGTAGAGGGAAATGATAGAGGGGACTGGCTAA<br>AGAATCTGCAATATGGAGTTTTTGGCCTTGGTAACAGACAATATGAGCATTTCAACAA<br>GATTGCTAAAGTGGTGGATGAGAAAGTTGCTGAACAGGGTGGTAAGCGGATTGTTCCA<br>TTGGGTCTGGGAGACGATGACCAGTGCATTGAAGATGACTTTGCTGCATGGCGTGAGA<br>ATGTATGGCCTGAGTTGGATAACTTGCTCCGGGATGAGGATGATACAACTGTTTCTAC<br>AACCTACACTGCTGCTATTCCAGAATATCGTGTTGTGTTCCCTGACAAATCAGATTCA<br>CTTATTTCAGAAGCAAATGGCCATGCCAATGGTTATGCTAATGGCAACACCGTATATG<br>ATGCCCAGCATCCTTGCAGATCTAATGTTGCAGTGAGGAAGGAGCTTCATACTCCAGC<br>ATCTGATCGTTCTTGCACCCATTTGGAGTTTGACATTGCTGGCACTGGCCTTTCATAT<br>GGAACTGGAGATCATGTTGGAGTGTACTGTGATAATCTATCTGAAACCGTGGAGGAGG<br>CTGAGAGATTACTGAATTTACCCCCAGAAACTTATTTCTCGCTTCATGCTGATAAAGA<br>GGATGGAACCCCACTTGCTGGGAGCTCATTGCCTCCTCCTTTCCCACCTTGTACTCTA<br>AGAACCGCCCTCACTCGTTATGCAGATCTCTTAAATACTCCTAAGAAGTCTGCTTTGT<br>TAGCTCTAGCAGCTTATGCATCTGATCCAAATGAGGCCGATCGTCTAAAATATCTTGC<br>TTCTCCAGCCGGAAAGGATGAATATGCTCAGTCACTAGTTGCAAATCAGAGAAGCCTC<br>CTCGAGGTCATGGCTGAATTTCCATCAGCAAAGCCTCCTCTTGGAGTATTCTTTGCAG<br>CAATTGCTCCACGCCTCCAACCCAGATTCTATTCTATATCGTCTTCTCCAAGGATGGC<br>ACCATCTAGAATTCATGTCACTTGTGCACTTGTTTATGAAAAAACACCTGGAGGACGA<br>ATTCACAAGGGTGTGTGTTCGACATGGATGAAGAATGCCATTCCATTGGAGGAAAGCC<br>GTGACTGCAGCTGGGCTCCTATCTTTGTCAGGCAGTCTAACTTCAAACTCCCTGCCGA<br>TCCTAAAGTGCCTGTTATAATGATCGGCCCTGGTACTGGACTAGCTCCCTTCAGAGGA<br>TTCCTTCAGGAAAGATTAGCTCTGAAGGAAGAAGGAGCTGAACTTGGTACTGCAGTTT<br>TCTTTTTTGGATGCAGGAACCGCAAAATGGATTACATCTATGAAGATGAGCTAAACCA<br>TTTCCTTGAAATTGGTGCACTTTCCGAGCTACTTGTTGCTTTCTCACGTGAGGGACCC<br>ACTAAGCAGTATGTGCAACACAAGATGGCAGAAAAGGCTTCTGATATTTGGAGGATGA<br>TTTCTGATGGAGCATATGTTTACGTCTGCGGTGATGCCAAAGGCATGGCCAGGGATGT<br>CCACAGAACTCTCCACACCATTGCTCAAGAGCAGGGATCGATGGATAGCACACAGGCT<br>GAGGGTTTTGTGAAGAATCTGCAAATGACCGGAAGGTATCTCCGAGATGTCTGGTGA | | |
| Cytochrome b5 | *Catharanthus roseus* | CrCYB5 | KP411010.1 |
| Iridoid oxidase | *Catharanthus roseus* | *Cr*IO | KF591593.1 |
| 7-Deoxyloganic acid hydroxylase | *Catharanthus roseus* | *Cr*7DLH | KF302067.1 |
| 7-Deoxyloganetic acid glycosyltransferase | *Catharanthus roseus* | *Cr*7DLGT | KF415118.1 |
| Alcohol dehydrogenase 2 | *Catharanthus roseus* | *Cr*CYPADH | KP411012.1 |
| Loganic acid O-methyltransferase | *Catharanthus roseus* | *Cr*LAMT | KF415116.1 |
| Secologanin synthase | *Catharanthus roseus* | *Cr*SLS | KM524261.1 |
| Strictosidine synthase | *Catharanthus roseus* | *Cr*STR | X61932.1 |
| 8-Hydroxygeraniol oxidoreductase | *Catharanthus roseus* | *Cr*8HGO | KF302069.1 |
| Geraniol-8-hydroxylase | *Catharanthus roseus* | *Cr*G8H | AJ251269.1 |
| Iridoid synthase | *Nepeta cataria* | *Nc*ISY2 | KY882234.1 |
| Iridoid cyclase | *Nepeta cataria* | *Nc*MLPLa | MT108281.1 |

<table>
<tr><td colspan="2">Geraniol synthase</td><td>Catharanthus roseus</td><td>CrGES</td><td>JN882024.1</td></tr>
<tr><td colspan="2">Tryptophan decarboxylase</td><td>Catharanthus roseus</td><td>CrTDC</td><td>MG748691.1</td></tr>
<tr><td rowspan="16">Membrane steroid binding protein</td><td></td><td>Arabidopsis thaliana</td><td>AtMSBP1</td><td>BT000922.1</td></tr>
<tr><td>AtMSBP1 amplified from cDNA</td><td colspan="3">ATGGCGTTAGAACTATGGCAAACTCTCAAAGAAGCAATCCATGCTTACACAGGTCTTTCTCCTG<br>TTGTCTTCTTCACTGCTCTAGCTCTCGCCTTCGCCATTTACCAAGTCATCTCAGGCTGGTTTGC<br>CTCGCCGTTCGATGATGTTAACCGACATCAGAGAGCTAGATCCTTGGCTCAAGAGGAGGAGCCA<br>CCGATTCCTCAGCCTGTTCAAGTCGGTGAGATCACGGAGGAGGAGCTTAAACAGTACGATGGCT<br>CTGATCCTCAAAAGCCCCTTCTTATGGCTATCAAACATCAGATCTATGATGTTACACAAAGCAG<br>GATGTTCTACGGACCAGGAGGACCATATGCTTTGTTTGCAGGGAAAGACGCTAGCCGAGCTCTT<br>GCAAAGATGTCATTTGAGGAGAAAGACTTGACTTGGGATGTCTCTGGTCTTGGTCCCTTTGAGC<br>TAGATGCTCTTCAAGATTGGGAGTACAAGTTCATGAGCAAGTATGCTAAGGTTGGTACTGTCAA<br>AGTGGCTGGTTCAGAACCTGAAACCGCATCTGTCTCTGAACCCACAGAGAATGTTGAGCAAGAT<br>GCTCATGTAACCACAACGCCTGGGAAGACCGTTGTTGATAAGAGTGATGATGCTCCTGCTGAGA<br>CTGTGTTGAAGAAGGAGGAGTAG</td></tr>
<tr><td></td><td>Arabidopsis thaliana</td><td>AtMSBP2</td><td>AF153283.1</td></tr>
<tr><td></td><td>Arabidopsis thaliana</td><td>AtMSBP3</td><td>NM_114748.3</td></tr>
<tr><td></td><td>Arabidopsis thaliana</td><td>AtMSBP4</td><td>NM_117583.8</td></tr>
<tr><td></td><td>Catharanthus roseus</td><td>CrMSBP1</td><td>This study</td></tr>
<tr><td>CrMSBP1 synthetic gene</td><td colspan="3">ATGGCGATTGCTCTGTGGACAACAATAACGGAGGCGATAGACCACTATACTGGACTTTCGCCGA<br>CGGCGTTTTTCACTATAATGGCTTTGATGGTGGTCACTTACAGAGTAGTGTGTGGAATGTTTGT<br>GGCAGCTGAGGATTATGTCGCGGTTAAGAAGGCGAACGAGTTTGTTGTTCGTGAACCGGTACAG<br>TTAGGTGAGGTGACGGAGGAAGAATTGAGGGAGTATGATGGCAAGGACCCGAATAAACCCTTGC<br>TTATGGCCATCAAAGGACAGATCTATGATGTTTCTCGTTCCAGGATGTTTTATGGTCCTGGTGG<br>ACCATACGCACTCTTTGCTGGTAGGGATGCAAGTCGTGCCTTAGCTCTCATGTCTTTCGACCCA<br>AAAGACCTCACCGGAAACATTGAAGGGCTCAGTGATTCAGAGCTGGAGGTTCTGCAAGACTGGG<br>AATACAAATTCATGGAGAAGTACGTTAAGGTGGGGCAGATTGTCTCGACGAAAACAAAAAATGA<br>AGGTGGAGAGAATGTACATCAAACTAACAATGGCGAAGCAAGTTCAGAACAATAA</td></tr>
<tr><td></td><td>Catharanthus roseus</td><td>CrMSBP2</td><td>This study</td></tr>
<tr><td>CrMSBP2 synthetic gene</td><td colspan="3">ATGGCCCTTCAATTATGGGAGACTTTCAAAGAATCAATCACGACGTATACAGGTCTATCTCCGG<br>CGACTTTCTTTACAGTTGTTGCTCTGGGCCTTGCCGTCTACTACTTGGTCTCAACCATGTTTGG<br>GTCCGATGAGCATGGTCATCACTCTAGGCCTAGAGAGTTTGAGGAACAGATGGAGCCTCTCCCA<br>CCTCCAGTTCAGCTTGGAGAGGTCACTGAGGAGGTCTTGAAGGAGTACGATGGTTCTGATCCCA<br>AGAAGCCTTTACTTATGGCGATTAAGGGGCAGATCTATGATGTTTCACAGAGCAGGATCTTTTA<br>TGGACCTGGTGGACCTTATCATCTCTTTGCAGGAAAGGATGCTAGCAGGGCCCTTGCAAAGATG<br>TCCTTTGAAGAAAAAGACCTTACGGGTGACATCTCTGGCCTTGGTGTCTTTGAGCTTGAGGCCT<br>TACAAGATTGGGAGTACAAGTTCATGAGCAAATATGTGAAGGTCGGAACTGTAAAGCAGTCAGT<br>GCCTGTGACTGAGGGAGAAGCTGCAGAAACTTCAGGTGAAGCTCACAGCGTGGACAAAAAACCT<br>GTTGAAGCTTCTGATCGCGATATTCCTGCTGCTCCTGAAAATGGTCCATTGGAAGGTGCCGCGG<br>AGAAGAAGATTGAAAGCAAAACCGTGGCCGAAGCTGACAAGAAAGAATAG</td></tr>
<tr><td></td><td>Camptotheca acuminata</td><td>CaMSBP1</td><td>This study</td></tr>
<tr><td>CaMSBP1 synthetic gene</td><td colspan="3">ATGGCCCTTCAACTATGGGAGACTCTCAAGGAATCGATCACAGTCTACACTGGACTGTCTCCGG<br>CGACCTTCTTTACAGTAGTTGCTTTGGGGCTTGCAATCTACTACGTCATATCAGGGCTGTTTGG<br>ATCGTCCGATCGTCAACAGAGGTCTAGGGATTTCGAAGAAAAGATGGAGCCTCTACCTCCTCCA<br>GTTCAGCTTGGGGAGATCAGTGAAGAGGAGTTGAAGGCTTATGATGGCAAAGATCCCAAAAAGC<br>CTCTGCTCATGGCGATCAAGGGTCAGATCTATGATGTGTCGCAGAGCAGGGTGTTTTATGGACC<br>TGGTGGGCCTTACGCTCTTTTTGCTGGGAAGGATGCTAGCAGAGCTCTTGCAAAGATGTCCTTC<br>GAAGAAAAAGATCTTACAGGGGATATATCCGGTCTTGGTCCATTTGAAATTGATGCCTTGCAAG<br>ACTGGGAATACAAGTTTATGAGCAAGTATGTTAAGGTTGGAACTGTTAAGAAGTCGGTAGCAGT<br>AACCGATGGGTCATCTGCTTCTGAACCTGCAGAAGCTACTGACTGTGATGTTGCTAAGCCTTCA<br>GAAGGTGAATCAAAGAGCCCTGCTGTTGAAACTGCAGAACCTGCATCTGTCGGTGATGCCAACA<br>AAGAGTAA</td></tr>
<tr><td></td><td>Camptotheca acuminata</td><td>CaMSBP2</td><td>This study</td></tr>
<tr><td>CaMSBP2 synthetic gene</td><td colspan="3">ATGGGTTTTTACACAACTTTGATGGAAGCGATTCCTGAATACACCGGCCTCTCACCAGCCGCAT<br>TCTTCACTATCGCCGCCTTCATGGTGGTGGTCTACAAGGTGGTGTGCGGAATGTTCGTGGCGGC<br>CGAAGATTACGTGGCTGTCAAAAATGCCAACCAGTACGCTCTACGCGAACCAGTCCAATTGGGA<br>GATGTTACCGAGGGCGAATTGAGGGCTTATGACGGTTCTGATCCAAGCAAGCCCTTGTTGATGG<br>CCATCAAAGGTCAGATCTACGATGTTTCTCGCTCCAGGATGTTTTATGGTCCTGGTGGGCCATA<br>TGCCCTGTTTGCTGGTAGGGATGCTAGCCGAGCCCTAGCTCTCATGTCGTTTGACCCTCAAGAC<br>CTTACTGGGAACATTGATGGTCTCAGTGCTTCTGAGCTTGAAGTTTTGCAAGACTGGGAATATA<br>AATTCATGGAGAAATATGTAAAGGTTGGGCAGATCGTTTCAGAAAGAATGAGTAATGAACAGAT<br>GGGGAATAAAGTGCAGGATACTCAGAATCTAGAAGGAAATGAATCCACCCCAAATTGA</td></tr>
<tr><td></td><td>Ailanthus altissima</td><td>AaMSBP1</td><td>This study</td></tr>
</table>

| | | | | |
|---|---|---|---|---|
| | *Aa*MSBP1 amplified from cDNA | ATGGCTCTGCAAGTATGGGAGACTCTTAAGGAGGCGATTACGGCGTACACGGGTCTGTCCCCGG<br>CGACCTTCTTTACGGTTCTTGCGTTACTGTGGGCCATATACTACGTGTTTTCCGGAATGTTTAG<br>GTCGTCCGATAATCACTATCAACAGAGATCCAGAGATTACGAGGAGCAGAGTGAGCCTCTGCCG<br>CCTCCGGTTCAGCTGGGGGAGATCGCCGAGGAGGAGCTGAAACAGTACGATGGCTCCGATCCCA<br>AAAAGCCCCTTCTCATGGCCATCAAGGGCCAGATCTACGATGTTTCTCAAAGCAGGATGTTTTA<br>TGGACCCGGTGGTCCATACGCATTGTTTGCCGGGAAGGATGCTAGCAGAGCACTTGCAAAGATG<br>TCTTTTGAGGAAAAAGATTTGACCGGTGATATATCTGGTCTTGGTCCATTCGAATTGGAGGCCT<br>TGCAGGACTGGGAGTACAAGTTTATGAGCAAGTACGTTAAGGTTGGAACTATCCAGAAAACAGT<br>TCTGGTAACAGAAGGAGCCTCCACTGCTGAATCTACAGAAACTAGAGAAGTAGATGTCGCCAAG<br>CCTGTGGAAACTAAAGAAGGAGATGACGCCAAACCTGTGGAAACTAAAGAAGGAGATGATGCAA<br>AACCTGTGGAAACTAAAGAAGGAGATGACAAGAAGCCTGCAGAAGATGGTCCATCAGAAACTGC<br>AGCTGTTGAAGCCGAGGAATCCAAATCTGGTAATGATGATAAGGAGTAA | | |
| | | *Citrus sinensis* | *Cs*MSBP1 | This study |
| | *Cs*MSBP1 amplified from cDNA | ATGGCTCTGCAACTATGGGAGACTCTAAAGGAGGCGATAACAGCGTACACGGGACTCTCCCCGG<br>CGGCGTTTTTTACAGTTCTTGCGTTGCTGTGGGCCATATACTATGTTCTGTCGGGAATGTTTGG<br>TTCGTCCGATAATCATCAGCAGCAGAGATCAAGGGAGTACGAGGAGCAGATGGAGCCTCTGCCA<br>CCGCCTGTTCAGCTCGGCGAGATTTCCGAGGAGGAGCTCAAACAGTACGATGGCTCCGATTCCA<br>ACAAGCCCCTTCTCATGGCCATCAAGAGCCAGATCTACGATGTCTCTCAGAGCAGGATGTTTTA<br>TGGACCTGGTGGGCCATATGCGTTGTTTGCTGGAAAGGAGGCTAGCAGGGCCCTTGCGAAGATG<br>TCTTTTGAGGAAAAAGATTTAACTGGTGATATCTCTGGTCTTGGTCCATTTGAGTTGGAGGCAT<br>TGCAGGACTGGGAGTATAAGTTCATGAGCAAGTATGTTAAGGTTGGATCTATCAAGTCGACAGT<br>TCCAGTAACAGATGGAGCCTCCTCTGGTGAATCCACTGAGCCTAAAGGAGTTGTTGATACACCT<br>GCAGAAACTAAAGAAGTAGATATCGCTAAGCCTGCTGAAGATGGTCCGTCAGAAACTGCAGCCG<br>CTGGACCTGTGGCAACCCCATCTTCTGATGATACTAAGAAGGAGTAA | | |
| | | *Citrus sinensis* | *Cs*MSBP2 | This study |
| | *Cs*MSBP2 amplified from cDNA | ATGGCTCTGCAACTATGGGAGACTCTAAAGGAGGCGATAACAGCGTACACGGGACTCTCCCCGG<br>CGGCGTTTTTTACAGTTGTTGCGTTGCTGTGGGCCATATACTATGTTCTGTCGGGAATGTTTGG<br>TTCGTCCGATAATCATCATCAGCAGAGATCGAGGGAGTACGAGGAACAGATGGAGCCTCTGCCA<br>CCGCCTGTTCAGCTAGGCGAGATTACCGAGGAGGAGCTCAAACAGTACGATGGCTCCGATTCCA<br>AAAAGCCCCTTCTCATGGCCATCAAGAGCCAGATCTACGATGTCTCTCAGAGCAGGATGTTTTA<br>TGGACCTGGTGGGCCATATGCGTTGTTTGCTGGAAAGGATGCTAGCAGGGCCCTTGCGAAGATG<br>TCTTTTGAGGAAAAAGATTTAACTGGTGATATCTCTGGTCTTGGTCCATTTGAGTTGGAGGCAT<br>TGCAGGACTGGGAGTATAAGTTCATGAGCAAGTATGTTAAGGTTGGATCTATCAAGTCGACAGT<br>TCCAGTAACAGATGGAGCCTCCTCTGGTGAATCCACTGAGCCTAAAGAAGGAGTTGTTGATACA<br>CCTGCAGAAAGTAAAGGAGTTGTTGATACACCTGCAGAAAGTAAAGGAGTTGTTGATACACCTG<br>CAGAAACTAAAGAAGTAGATATCGCTAAGCCTGCTGAATATGGTCCGTCAGAAACTGCAGCCGC<br>TGGACCTGAGGCAACCCCATCTTCTGATGATACTAAGAAGGAGTAA | | |
| CYP72A565 | | *Camptotheca acuminata* | *Ca*CYP72A565 | MH763569.1 |
| Codon optimised CYP72A565 | | ATGAAGATGGAAGTCATGCATATGTCTGTTGCCGCTTCGCTTGCTGTTGTATTCTTAGTCTGCA<br>TTTGGCGTGCTTTGAATTGGGCGTGGTTTATGCCGAAGAAAATAGAGAAGAGGTTGAGGCAACA<br>GGGATTTAACGGCAATCCATACAGACTACTTGTAGGTGATCTAAAGGAATCTTCTATGATGTTA<br>AAAGAGGCGATGTCGAAACCTATACCTGTATCACAGGATATAGTGCAAAGGTTAATGCCGCATG<br>TAGTAAAGACTATACAGACGTACGGTAAGAACTCTTTCACATGGATTGGGAGAATGCCTAGAGT<br>GCATATAATGGAGCCAGAGCTAATTAAAGACATTTTAGCTAATCACAATAATTTCCAGAAGAAC<br>CACCACGCTTATAACCCTCTGACTAAATTCTTGTTGACCGGTATAGGGTCTTTAGAAGGGGAAA<br>AGTGGGCTAAGCATAGGCGTATTATTTCGCCATCCTTTCACTTAGAAAAGCTTAAAACCATGTT<br>ACCAGCTTTCTACGTTTCATACGACGAACTACTTGGAAAATGGGAACGTGAATCATCAACCAAA<br>GGTAGTGTCGAAGTTGACTTGTTCCCAACGTTCGACACATTGACTTCGGACGTTATCAGTCGTG<br>TTGCATTCGGCTCATCGTACGGTGAGGGTGGCAGAATCTTTATCTTGTTAAAGGAGTTAATGGA<br>TCTTACAGTCGACGTAATGAGGAGCGTATACGTTCCGGGTTGGTCATTGCTACCTACGAAAAGA<br>AATCAACGTATGAGAGAGGTAGATAGGGAGATTAGGGAAAGACTTTCTGGAATAATAAATTCAA<br>GGGTCAAAGCTATGAAGGCCGGTGAACCATCCGGAGACGATCTTTTAGGGACTTTATTGGAGTC<br>TAACTTTAGGGAAATCGAGAGGTTAGGGAATAAGAAGAACGCGGGCATGTCAATCGAAGACGTT<br>ATATCCGAGTGTAAACTGTTTTATTTTGCAGGACAAGAAACTACTGGCATACTACTTACATGGA<br>CCTGCGTCATCTTATCGAGACATCCTGAGTGGCAGGAAAGAGCACGTGAGGAAATATTTCAGGT<br>TTTCGGTAACGGTAAGTTAGATTTCGACAGAGTTCAAGGATTAAAGATAGTACCCATGATATTA<br>TACGAAGTGCTGAGGCTATATCCCCCTGTAATCGAGTTAACCAAGGTAACATACGAGGAACAGA<br>AACTAGGTAACCTGACCATTCCAGCTGGTGTACAATTGATGATGCCGTCTATTCTTTTGCATAG<br>AGATAAAGAAATGTGGGGTGACGACGCTACCGAGTTTAACCCGGGCCGTTTCGCAGAGGGTGTC<br>GCAAAAGCGGTCAAATCACCTTTCTTCTATATACCATTTTCCTGGGGCCCAAGAATATGTGTCG<br>GCCAGAATTTTGCTTTATTGCAGGCAAAGATGGCGTTGGCAATGATATTGCAGAGATTTTCATT<br>CGACTTATCTCCTACATACGCTCACGCACCATTTACCGTTCTTACGTTGCAACCACAACACGGA<br>GCCCAGGTTATATTTAGAAGATTAAAGTGCTAG | | |
| CYP72A610 | | *Camptotheca acuminata* | *Ca*CYP72A610 | MH763570.1 |
| Codon optimised CYP72A610 | | ATGGAAATTCAGATGGATGTTCTATACAAGTCCATAGCAGCCAGTGTGGCCGTTGTATTTCTAG<br>TATATGCGTGGAAAATGCTAAACTGGGCATATTTAACACCAAAGAGGATTGAGAAATGCTTGAG<br>AAAGCAAGGTTTCAAAGGCAACAGCTACCGTTTGTTGGTGGGCGATTTAAAAGAATCTTCAATG<br>ATGCTGAAAGAAACCATGTCTAAGCCTATCAACGTTTCTGAAGACATTGTCCAAAGGGTCATGC<br>CACATGTCATTAAAACTATCGATACCTATGGTAAGAATTCCTTTACTTGGATAGGTAGAATGCC<br>TAGGGTGCACATCATGGAACCAGATCTAATTAAGGATATTCTAGCAAACCACAACGACTTCATG<br>AAGAATCACCATGCATACAACCCGCTTACTAAATTCCTTTTGACTGGAATTGGCTCCCTAGAAG | | |

| | | | |
|---|---|---|---|
| | GGGACAAGTGGGCCAAACACAGAAGAATTATTTCTCCTTCTTTCCATTTGGAGAAACTAAAGAC<br>TATGCTACCTGCCTTTTACGTCTCGTACGATGATTTATTGACAAAATGGGAGCAACAATGCTCT<br>TCGAAGGGCAGCGTTGAGATTGATTTATTTCCAACATTTGATACACTTACCTCTGATGTTATTT<br>CCAGAGTCGCTTTTGGCAGTTCCTATGGCGAGGGCGGAAGAATTTTCATTCTTTTAAAAGAACT<br>TATGGACTTAACAGTAGATGTAATGAGATCTGTCTACGTCCCGGGTTCTAGTTTTCTACCTACC<br>AAGAGGAACAACAGGATGAGAGAAGTTGATGGTGAGATCAAAGACAGGCTGTCCGGTATCATTA<br>ATTCAAGAGTTAAGGCGATGAAGGCCGGTGAGCCTAGTGGCGAAGACCTATTGGGTACCCTGTT<br>GGAATCTAATTTCAAAGAGATAGAAAGGTTAGGAAACAAGAAGAACGCCGGTATGAGTATAGAA<br>GATGTCATCTCTGAATGCAAGCTGTTCTACTTTGCGGGCCAGGAGACGACCGGAATTTTGCTAA<br>CCTGGACCTGCGTTCTTTTGTCGAGGCATCCCGAATGGCAAGAAAGAGCCAGAGAGGAAATTTT<br>CCAGGTTTTCGGAAATGGAAAAGTGGACTTTGATCGTGTCCAGAATCTGAAAATCGTACCAATG<br>ATATTGTATGAGGTGCTTAGGTTATATCCCCCAGTTATAGAGCTTACGAAGGTGACCTATGAAG<br>AGCAAAAGTTAGGTAACTTGACTATCCCTGCTGGGGTCCAGCTGATGATGCCTTCAATACTTCT<br>TCACAGGGACCAAGAGATGTGGGGCGCGGATTCAAAGGAATTCAATCCAGGTAGATTTGCAGAT<br>GGAATAAGCAAAGCAGTCAAGAGTCCGTTCTTCTACATTCCGTTCTCATGGGGCCCGAGAATTT<br>GCGTTGGCCAAAACTTTGCCCTATTGCAAGCTAAAATGGCGTTGACAATGATTCTACAAAGGTT<br>CACATTCGATTTAAGCCCAACATATGCACATGCCCCTTTCACAGTACTAACGCTACAGCCTCAA<br>CATGGCGCCCAAGTCGTGTTTAGAAAGATAAAGTGTTAG | | |
| Medium-chain dehydrogenase/reductase | *Camptotheca acuminata* | *Ca*MDR | This study |
| *Ca*MDR sequence | ATGGCGAAATCACCGGAACAGGAACACCCAGTAAAGGCTTTTGGATGGGCAGCCAGAGACACAT<br>CTGGTGTCCTTTCTCCATTCAATTTCTCACGCAGAGCAACCGGCGAGAAAGATGTGACGTTCAA<br>AGTGCTGTATTGTGGAATCTGTCACTCCGACCTTCATACAATCAAGAATGAATGGGGCGCAACT<br>AAGTACCCTATTATCCCCGGGCACGAGCTTGCTGGTGTCGTAACAGAGGTAGGCAGCAAGGTAC<br>AAAAATTTAAAGTTGGAGACAGAGTTGGGGTCGGCTGCCTAGTTGGAGCTTGTCATTCATGCGA<br>TAGCTGTGGCCATGATCTCGAAAATTACTGTCATAAAGCCATATTCACCTATGGTTCGACCTAC<br>TATGATGGAACCCCCACACATGGAGGTTACTCTAACATCATGGTAGCTAACGAGCACTTTGTGG<br>TTCGCATTCCTGACAATTTGCCTCTTGATGCTGCTGCTCCTCTTTTGTGTGCTGGGATCACAAC<br>TTACAGCCCCCTGAAATATTTTGGGCTTGACAAACCTGGTTTGCATGTTGGTGTGGTTGGTCTA<br>GGTGGGCTCGGCCATGTGGCTGTGAAGTTTGCAAAGGCTTTTGGTGCTAAGGTCACCGTGATCA<br>GTACATCCCCAAACAAAAAGGATGAGGCCCTTAAACATCTTGGTGCGGATTCATTTTTGGTCAG<br>CCGCGACCCTGATCAGATACAGGCTGCAGCGGGCACGATGGATGGTATCATTGATACCGTCTCT<br>GCAATTCACCCTCTCCTACCATTAATTGGCTTGTTGAAATCTCATGGAAAGCTCATTATGGTTG<br>GGGCACCAGAGAAGCCACTTGAGCTACCAGTCATGCCTTTGCTTATGGGGAGGAAGATTGTGGC<br>TGGGAGTTGCATTGGAGGCATGAAGGAGACACAAGAGATGATGGATTTTGCAGCAAAACATAAT<br>GTAACAGCAGACATTGAGGTTATTTCAGTGGACTACGTAAACACTGCAATGGAGCGGCTCGCAA<br>AAGCCGATGTTAGATACCGATTCGTCATTGATATTGGGAACACATTGAAAGCTGCCTAG | | |

**Supplementary Table 3. List of primers used for USER cloning of genes in this study.**

Overhangs for USER cloning are highlighted in bold.

Start and stop codons are underlined.

| Primers for USER cloning of genes | | Comment |
|---|---|---|
| p179_BB1_CrCPR_f | **AGTGCAGGU**ATGGATTCTAGCTCGGAGAAGTTG | fw/rv primer for amplification of CPR from *Catharanthus roseus* |
| p180_BB1_CrCPR_r | **CGTGCGAU**TCACCAGACATCTCGGAGATACCTT | |
| p181_BB4_CrCYPb5_f | **ATCTGTCAU**ATGGCGTCGGATCAGAAATTGC | fw/rv primer for amplification of Cyb5 from *Catharanthus roseus* |
| p182_BB4_CrCYPb5_r | **CACGCGAU**TTACTTCTCCTTAGTATAGTGTCGGAC | |
| p278_BB1_CrCYPADH_f | **AGTGCAGGU**ATGCAGATCATAACTTGCAAGGCTGTG | fw/rv primer for amplification of CYPADH from *Catharanthus roseus* |
| p279_BB1_CrCYPADH_r | **CGTGCGAU**TCACAATGTGATGAGAACCTTCACGC | |
| p288_BB1_CrIO_f | **AGTGCAGGU**ATGGCGACCATCACTTTCGATT | fw/rv primer for amplification of IO from *Catharanthus roseus* |
| p289_BB1_CrIO_r | **CGTGCGAU**TTAGATATGAACTCTCTTCTTAGGGAT | |
| p290_BB4_Cr7DLGT_f | **ATCTGTCAU**ATGGGTTCTCAAGAAACAAATTT | fw/rv primer for amplification of 7DLGT from *Catharanthus roseus* |
| p291_BB4_Cr7DLGT_r | **CACGCGAU**TCAAATAATCAGTGATTTTATGTAATC | |
| p302_BB4_Cr7DLH_f | **ATCTGTCAU**ATGGAATTGAACTTCAAATCAATTATTTTC | fw/rv primer for amplification of 7DLH from *Catharanthus roseus* |
| p303_BB4_Cr7DLH_r | **CACGCGAU**TTAGAGTTTGTGCAGAATCAAATGAG | |
| p346_BB1_CrSTR_f | **AGTGCAGGU**ATGGCAAACTTTTCTGAATCTAAATCC | fw/rv primer for amplification of STR from *Catharanthus roseus* |
| p347_BB1_CrSTR_r | **CGTGCGAU**CTAGCTAGAAACATAAGAATTTCCCTTG | |
| P404_BB1_CrGOR_f | **AGTGCAGGU**ATGACCAAGACCAATTCCCCTGC | fw/rv primer for amplification of 8HGO from *Catharanthus roseus* |
| P405_BB1_CrGOR_r | **CGTGCGAU**TTAGAACTTGATAACAACTTTGACACAATCA | |
| P406_BB4_CrG8H_f | **ATCTGTCAU**ATGGATTACCTTACCATAATATTAACTTTAC | fw/rv primer for amplification of G8H from *Catharanthus roseus* |
| P407_BB4_CrG8H_r | **CACGCGAU**TTAAAGGGTGCTTGGTACAGC | |
| P412_BB1_CoCa565_f | **AGTGCAGGU**ATGGAAATTCAGATGGATGTTCTATACAAGT | fw/rv primer for amplification of CaCYP72A565$^{opt}$ from *Camptotheca acuminata* |
| P413_BB1_CoCa565_r | **CGTGCGAU**CTAACACTTTATCTTTCTAAACACGACTTGG | |
| P414_BB4_CoCa610_f | **ATCTGTCAU**ATGAAGATGGAAGTCATGCATATGTCTG | fw/rv primer for amplification of CaCYP72A610$^{opt}$ from *Camptotheca acuminata* |
| P415_BB4_CoCa610_r | **CACGCGAU**CTAGCACTTTAATCTTCTAAATATAACCTGGGC | |
| P468_BB1_CrLAMT_f | **AGTGCAGGU**ATGGTTGCCACAATTGATTCCATT | fw/rv primer for amplification of LAMT from *Catharanthus roseus* |
| P469_BB1_CrLAMT_r | **CGTGCGAU**TTAATTTCCCTTGCGTTTCAAGACAA | |
| P470_BB4_CrSLS_f | **ATCTGTCAU**ATGGAGATGGATATGTATACCATTAG | fw/rv primer for amplification of SLS from *Catharanthus roseus* |
| P471_BB4_CrSLS_r | **CACGCGAU**CTAACTCTCAAGCTTCTTGTAGATG | |
| P559_BB1_NcISY2_f | **AGTGCAGGU**ATGAGCATGAACTGGTGGAG | fw/rv primer for amplification of ISY2 from *Nepeta cataria* |
| P560_BB1_NcISY2_r | **CGTGCGAU**TTAAGAAATAGTAGAGGAAGGAAC | |
| P561_BB4_NcMLPLa_f | **ATCTGTCAU**ATGGCTTCCAAGCTTGAAATAG | fw/rv primer for amplification of MLPLa from *Nepeta cataria* |
| P562_BB4_NcMLPLa_r | **CACGCGAU**TTAATTTTGACATGTGTGGTTCATG | |
| p627_BB1_tCrGES_f | **AGTGCAGGU**ATGTCATCCATGTCTCTGCC | fw/rv primer for amplification of truncated GES from *Catharanthus roseus* |
| p628_BB1_tCrGES_r | **CGTGCGAU**TTAAAAACAAGGTGTAAAAAACAAAGC | |
| p629_BB4_Erg20ww_f | **ATCTGTCAU**ATGGCTTCAGAAAAAGAAATTAGG | fw/rv primer for amplification of Erg20 from *Saccharomyces cerevisiae* |
| p630_BB4_Erg20ww_r | **CACGCGAU**CTATTTGCTTCTCTTGTAAACTTTG | |
| p640_BB1_tHMGR_f | **AGTGCAGGU**ATGCAATTGGTGAAAACTGAAG | fw/rv primer for amplification of truncated HMGR from *Saccharomyces cerevisiae* |
| p641_BB1_tHMGR_r | **CGTGCGAU**TTAGGATTTAATGCAGGTGACG | |
| p642_BB4_IDI1_f | **ATCTGTCAU**ATGACTGCCGACAACAATAGTATG | fw/rv primer for amplification of IDI1 from *Saccharomyces cerevisiae* |
| p643_BB4_IDI1_r | **CACGCGAU**TTATAGCATTCTATGAATTTGCCTGTC | |
| p745_BB1_CrTDC_f | **AGTGCAGGU**ATGGGCAGCATTGATTCAACAAATG | fw/rv primer for amplification of TDC from *Catharanthus roseus* |
| p746_BB1_CrTDC_r | **CGTGCGAU**TCAAGCTTCTTTGAGCAAATCATCG | |
| p747_BB4_ScZWF1_f | **ATCTGTCAU**ATGAGTGAAGGCCCCGTCAAATTC | fw/rv primer for amplification of ZWF1 from *Saccharomyces cerevisiae* |
| p748_BB4_ScZWF1_r | **CACGCGAU**CTAATTATCCTTCGTATCTTCTGGCTTAGTC | |
| P812_BB1_CoCaCYP72A610_fw | **AGTGCAGGU**ATGAAGATGGAAGTCATGCATATGTCTG | fw/rv primer for amplification of codon optimised CYP72A610$^{opt}$ from *Camptotheca acuminata* |
| P813_BB1_CoCaCYP72A610_rv | **CGTGCGAU**CTAGCACTTTAATCTTCTAAATATAACCTGGGC | |
| p886_BB1_At_Msbp1_fw | **AGTGCAGGU**ATGGCGTTAGAACTATGGC | fw/rv primer for amplification of MSBP1 from *Arabidopsis thaliana* |
| p887_BB1_At_Msbp1_rv | **CGTGCGAU**CTACTCCTCCTTCTTCAAC | |
| P1284_BB1_AtMSBP2_fw | **AGTGCAGGU**ATGGTTCAGCAAATATGGGAGACG | fw/rv primer for amplification of MSBP2 from *Arabidopsis thaliana* |
| P1285_BB1_AtMSBP2_rv | **CGTGCGAU**TTAAACCCTGAAAAATCTCACTAATTTCTC | |
| P1286_BB1_AtMSBP3_fw | **AGTGCAGGU**ATGGTTCAGCAAATATGGGAGACG | fw/rv primer for amplification of MSBP3 from *Arabidopsis thaliana* |
| P1287_BB1_AtMSBP3_rv | **CGTGCGAU**TTACTCCTTTGCAGCATCATCATCATC | |
| P1288_BB1_AtMSBP4_fw | **AGTGCAGGU**ATGATTCCGGCGAGGAGGTTTC | fw/rv primer for amplification of MSBP4 from *Arabidopsis thaliana* |
| P1289_BB1_AtMSBP4_rv | **CGTGCGAU**TCAAACCCTGCAAGTCTTGGCAAG | |
| P1290_BB1_CrMSBP1_fw | **AGTGCAGGU**ATGGCGATTGCTCTGTGGACAAC | fw/rv primer for amplification of MSBP1 from *Catharanthus roseus* |
| P1291_BB1_CrMSBP1_rv | **CGTGCGAU**TTATTGTTCTGAACTTGCTTCGCCATTG | |
| P1292_BB1_CrMSBP2_fw | **AGTGCAGGU**ATGGCCCTTCAATTATGGGAGAC | fw/rv primer for amplification of MSBP2 from *Catharanthus roseus* |
| P1293_BB1_CrMSBP2_rv | **CGTGCGAU**CTATTCTTTCTTGTCAGCTTCGGCC | |
| P1294_BB1_CaMSBP1_fw | **AGTGCAGGU**ATGGCCCTTCAACTATGGGAGAC | fw/rv primer for amplification of MSBP1 from *Camptotheca acuminata* |
| P1295_BB1_CaMSBP1_rv | **CGTGCGAU**TTACTCTTTGTTGGCATCACCGACAG | |
| P1296_BB1_CaMSBP1_fw | **AGTGCAGGU**ATGGGTTTTTACACAACTTTGATGGAAGC | |

| | | |
|---|---|---|
| P1297_BB1_CaMSBP1_rv | **CGTGCGAU**TCAATTTGGGGTGGATTCATTTCCTTC | fw/rv primer for amplification of MSBP2 from *Camptotheca acuminata* |
| P1298_BB1_AaMSBP1_fw | **AGTGCAGGU**ATGGCTCTGCAAGTATGGGAGAC | fw/rv primer for amplification of MSBP1 from *Ailanthus altissima* |
| P1299_BB1_AaMSBP1_rv | **CGTGCGAU**TTACTCCTTATCATCATTACCAGATTTGGA | |
| P1300_BB1_CsMSBP1_fw | **AGTGCAGGU**ATGGCTCTGCAACTATGGGAGAC | fw/rv primer for amplification of MSBP1 from *Citrus sinensis* |
| P1301_BB1_CsMSBP1_rv | **CGTGCGAU**TTACTCCTTCTTAGTATCATCAGAAGATGG | |
| P1302_BB1_CsMSBP2_fw | **AGTGCAGGU**ATGGCTCTGCAACTATGGGAGAC | fw/rv primer for amplification of MSBP2 from *Citrus sinensis* |
| P1303_BB1_CsMSBP2_rv | **CGTGCGAU**TTACTCCTTCTTAGTATCATCAGAAGATGG | |
| P1304_BB1_ScDap1_fw | **AGTGCAGGU**ATGTCCTTCATTAAAAACTTGTTATTTGG | fw/rv primer for amplification of Dap1 from *Saccharomyces cerevisiae* |
| P1305_BB1_ScDap1_rv | **CGTGCGAU**TCATACGTTCACGCCAGGCTC | |

### Supplementary Table 4. List of primers used for USER cloning of promoters in this study.

Overhangs for USER cloning are highlighted in bold.

| Primers for USER cloning of promoters | | Comment |
|---|---|---|
| p183_BB2_pADH2_f | **ACCTGCACU**TGTGTATTACGATATAGTTAATAGTTG | fw/rv primer for amplification of ADH2 promoter from *Saccharomyces cerevisiae* |
| p184_BB2_pADH2_r | **AGTAGCTAU**TATCTAAAAATTGCCTTATGATCCG | |
| p185_BB3_pPCK1_f | **ATGACAGAU**GTTGTTATTTTATTATGGAATAATTA | fw/rv primer for amplification of PCK1 promoter from *Saccharomyces cerevisiae* |
| p186_BB3_pPCK1_r | **ATAGCTACU**ATAGGAAAAAACCGAGCTTC | |
| p282_BB3_pICL1_f | **ATGACAGAU**TTTTCGTTGACTTTTTGTTATGTT | fw/rv primer for amplification of ICL1 promoter from *Saccharomyces cerevisiae* |
| p283_BB3_pICL1_r | **ATAGCTACU**ATTTATTGAAAAGTAAATATCTCG | |
| p284_BB2_pPCK1_f | **ACCTGCACU**GTTGTTATTTTATTATGGAATAATTAG | fw/rv primer for amplification of PCK1 promoter from *Saccharomyces cerevisiae* |
| p285_BB2_pPCK1_r | **AGTAGCTAU**ATAGGAAAAAACCGAGCTTC | |
| p286_BB3_pADH2_f | **ATGACAGAU**TGTGTATTACGATATAGTTAATAGTTG | fw/rv primer for amplification of ADH2 promoter from *Saccharomyces cerevisiae* |
| p287_BB3_pADH2_r | **ATAGCTACU**TATCTAAAAATTGCCTTATGATCCG | |
| p336_BB2_pICL1_f | **ACCTGCACU**TTTTCGTTGACTTTTTGTTATG | fw/rv primer for amplification of ICL1 promoter from *Saccharomyces cerevisiae* |
| p337_BB2_pICL1_f | **AGTAGCTAU**ATTTATTGAAAAGTAAATATCTCG | |
| p348_BB2/2_pADH2_r | **CACGCGAU**TATCTAAAAATTGCCTTATGATCCGTCTCTCCGGTT | rv primer for amplification of ADH2 promoter from *Saccharomyces cerevisiae* |
| p644_BB2_pMLS1_f | **ACCTGCACU**TTTCTTAATTCTTTTATGTGCTTTTACTAC | fw/rv primer for amplification of MLS1 promoter from *Saccharomyces cerevisiae* |
| p645_BB2_pMLS1_r | **AGTAGCTAU**CCATTGGGCCGATGAAGTTAG | |
| p657_BB3_pMLS1_f | **ATGACAGAU**TTTCTTAATTCTTTTATGTGCTTTTACTAC | fw/rv primer for amplification of MLS1 promoter from *Saccharomyces cerevisiae* |
| p658_BB3_pMLS1_r | **ATAGCTACU**CCATTGGGCCGATGAAGTTAG | |

**Supplementary Table 5. List of primers used for generating gRNA plasmids in this study.**

| Primers for gRNA plasmid generation | | Comment |
|---|---|---|
| p483_gRNA_OYE2_U | TGGGGGTTACGATCATTTATCTTTCACTGC | fw/rv primer for gRNA plasmid to delete OYE2 from *Saccharomyces cerevisiae* |
| p484_gRNA_OYE2_D | GATTGTGGAGGTTTTAGAGCTAGAAATAGC | |
| p485_gRNA_OYE3_U | GTCTGATGAGGATCATTTATCTTTCACTGC | fw/rv primer for gRNA plasmid to delete OYE3 from *Saccharomyces cerevisiae* |
| p486_gRNA_OYE3_D | CAAATCCCAGGTTTTAGAGCTAGAAATAGC | |
| p487_gRNA_ATF1_U | AGAATGCCTGGATCATTTATCTTTCACTGC | fw/rv primer for gRNA plasmid to delete ATF1 from *Saccharomyces cerevisiae* |
| p488_gRNA_ATF1_D | TGTTGCACGGGTTTTAGAGCTAGAAATAGC | |
| p489_gRNA_ARI1_U | TTCTGGCGCAGATCATTTATCTTTCACTGC | fw/rv primer for gRNA plasmid to delete ARI1 from *Saccharomyces cerevisiae* |
| p490_gRNA_ARI1_D | ACGAAAACAGGTTTTAGAGCTAGAAATAGC | |
| p503_gRNA_ADH6_U | TCTTAATGTCGATCATTTATCTTTCACTGC | fw/rv primer for gRNA plasmid to delete ADH6 from *Saccharomyces cerevisiae* |
| p504_gRNA_ADH6_D | TCGAAGCATGGTTTTAGAGCTAGAAATAGC | |
| p651_Erg20p_gRNA_f | TATTTATCGGGATCATTTATCTTTCACTGC | fw/rv primer for gRNA plasmid to delete Erg20 promoter from *Saccharomyces cerevisiae* |
| p652_Erg20p_gRNA_r | GAGGAAGCAAGTTTTAGAGCTAGAAATAGC | |
| P1167_gRNA_CrGES_fw | CCTTGTTATCTCAGACAATTGATCATTTATCTTTCACTGC | fw/rv primer for gRNA plasmid to delete GES from *Catharanthus roseus* in *Saccharomyces cerevisiae* |
| P1168_gRNA_CrGES_rv | AATTGTCTGAGATAACAAGGGTTTTAGAGCTAGAAATAGC | |
| p1372_gRNA_CrLAMT_fw | GAAGCCCACCCAATGAAAGGGTTTTAGAGCTAGAAATAGC | fw/rv primer for gRNA plasmid to delete LAMT from *Catharanthus roseus* in *Saccharomyces cerevisiae* |
| p1373_gRNA_CrLAMT_rv | CCTTTCATTGGGTGGGCTTCGATCATTTATCTTTCACTGC | |
| p1427_gRNA_ADH3_fw | CAAGCCGCCAAAATTCAACAGTTTTAGAGCTAGAAATAGC | fw/rv primer for gRNA plasmid to delete ADH3 from *Saccharomyces cerevisiae* |
| p1428_gRNA_ADH3_rv | TGTTGAATTTTGGCGGCTTGGATCATTTATCTTTCACTGC | |
| p1429_gRNA_ADL6_fw | AACGGTAGAACAATCAACACGTTTTAGAGCTAGAAATAGC | fw/rv primer for gRNA plasmid to delete ALD6 from *Saccharomyces cerevisiae* |
| p1430_gRNA_ADL6_rv | GTGTTGATTGTTCTACCGTTGATCATTTATCTTTCACTGC | |
| p1431_gRNA_YPL062W_fw | CATATGCAACAATGACGTCTGTTTTAGAGCTAGAAATAGC | fw/rv primer for gRNA plasmid to delete YPL062W from *Saccharomyces cerevisiae* |
| p1432_gRNA_YPL062W_rv | AGACGTCATTGTTGCATATGGATCATTTATCTTTCACTGC | |
| p1471_gRNA_7DLH2_fw | TGTCCACACGAGTAAAGTCGGTTTTAGAGCTAGAAATAGC | fw/rv primer for gRNA plasmid to delete 7DLH from *Catharanthus roseus* in *Saccharomyces cerevisiae* |
| p1472_gRNA_7DLH2_rv | CGACTTTACTCGTGTGGACAGATCATTTATCTTTCACTGC | |

### Supplementary Table 6. List of primers used for generating integration fragments in this study.

| IF-fragments for repairing double strand breaks | | |
|---|---|---|
| p475_IF_OYE2_U | GAATAAATCATCATATTAAGCTAAATATAGACGATAATATAGTATCGATATAGTGTTAAC | primer to generate integration fragment to repair DSB for OYE2 deletion |
| p476_IF_OYE2_D | ATATAAATTAGAAGAAAAAGAAATGGTGCTACAAAGTACGGTTAACACTATATCGATACT | |
| p477_IF_OYE3_U | TGATATATACAACAACTGTAGTTCAGTATAGCGAAGTTTAAATTTAGAAGAATCATGAAT | primer to generate integration fragment to repair DSB for OYE3 deletion |
| p478_IF_OYE3_D | TATGGCAGGAATATGAAAAATACATAACATCAATGTCTTTATTCATGATTCTTCTAAATT | |
| p479_IF_ATF1_U | AAAAACGGCACTTCATCAGTATCACAAATACCATCAATTTATCAGCTCTCATCTCACATG | primer to generate integration fragment to repair DSB for ATF1 deletion |
| p480_IF_ATF1_D | CACGACATAATCATATTGTCGAATAATATCAGTCAAGCATCATGTGAGATGAGAGCTGAT | |
| p481_IF_ARI1_U | TAATTGTGCATTGTACAACTGTGCTAAACAGACTTAAAAAAGTAATAATTGTATCACGCT | primer to generate integration fragment to repair DSB for ARI1 deletion |
| p482_IF_ARI1_D | ATAGATTTGCCTATTGGAGTGATCAAAAAAAACTTCAATTAGCGTGATACAATTATTACT | |
| p499_IF_ADH6_U | CATGGAGCAGTTAAAAAGAAAGGAGCTACATTTATCAAGAGCTTGACAACGATTTTGGCT | primer to generate integration fragment to repair DSB for ADH6 deletion |
| p500_IF_ADH6_D | ATCCACATTCGAGGAAGAAATTCAACACAACAACAAGAAAAGCCAAAATCGTTGTCAAGC | |
| p653_Erg20p_IF_f | GGATACTGTCCTTATTACTGCGATATACAGTGTGAGGTATTCTAAGCGGTTGCAGGTCTCATCTGGAATATAATTC | primer to generate integration fragment to repair DSB for Erg20 promoter switch |
| p654_Erg20p_IF_r | GGGAAAACGTTCAAGAATCTCTCTCTCCTAATTTCTTTTTCTGAAGCCATGATTTTACGTATATCAACTAGTTGACGATT | |
| P810_IF_XI3_repair_fw | ACTGATTAGTTTTCCGTTTTAGGATATTGACGCCAAGCGTGCGTCTGATTTCTACGATATGTCTCTAATTTTGGAAGGGCCATTTTATTTTTGAG | primer to generate integration fragment to repair DSB for locus XI3 |
| P811_IF_XI3_repair_rv | ATGTACAAAACAGTTTAATAATGATCTGTATTGCTGGCTCAATCCACGTAAGGAAAATGATCAGCCCAATCCTCAAAAATAAAATGGCCCTTCCA | |
| P1279_IF_repair_XII5_new_fw | ACCGGTACCGGAGGAGACCGCTATAACCGGTTTGAATTTATTGTCACAGTGTCACATCAGCGGCAACTCAG | primer to generate integration fragment to repair DSB for locus XII5 |
| P1280_IF_repair_XII5_new_rv | AACTAAAACAATAAGGCTAGTTCGAATGATGAACTTGCTTGCTGTCAAACTTCTGAGTTGCCGCTGATGTGA | |
| p1374_LAMT_del_repair_fw | ATAAGAAATTCGCTTATTTAGAAGTGTCAACAACGTATCTACCAACGGAATGCGTGCGATTATCTAAAAATTG | primer to generate integration fragment to repair DSB for LAMT deletion |
| p1375_LAMT_del_repair_rv | GGATTAATCAGTTACACAGGCTGTAACCGGAGAGACGGATCATAAGGCAATTTTTAGATAATCGCACGCATTC | |
| p1433_IF_ADH3_repair_fw | TATATTATCTTCTGTTCACAGTTAAAACTAGGAATAGTATAGTCATAAGTTAACACCATCTAGCGTGTTAC | primer to generate integration fragment to repair DSB for ADH3 deletion |
| p1434_IF_ADH3_repair_rv | TATAAACCTCATCATTATAAACAAAGACTTTCATAAAAAGTTTGGGTGCGTAACACGCTAGATGGTGTTAA | |
| p1435_IF_ALD6_repair_fw | TAGAAGAAAAAACATCAAGAAACATCTTTAACATACACAAACACATACTATCAGAATACATGTACCAACCTG | primer to generate integration fragment to repair DSB for ALD6 deletion |
| p1436_IF_ALD6_repair_rv | CAAGTAAGTTTATATGAAAGTATTTTGTGTATATGACGGAAAGAAATGCAGGTTGGTACATGTATTCTGATAG | |
| p1437_IF_YPL062W_repair_fw | TAGACACACTATCAGGTCAGGAACTGCCGTCACATACGACACTGCCCCTCACGTAAGGGCCACCGACCAT | primer to generate integration fragment to repair DSB for YPL062W deletion |
| p1438_IF_YPL062W_repair_rv | TTGCTTGTCTTGAATCCCCCTCACCCCGAATTTATTACGAATTTGCCCACATGGTCGGTGGCCCTTAC | |

### Supplementary Table 7. List of primers used for sequencing and colony PCR

| Sequencing and colony PCR | |
|---|---|
| p187_C_cPCR_pADH2 | GTTTTTATCACTTCTTGTTTCTTC |
| p188_C_cPCR_pPCK1 | CAGCTTAAACAATAATTATATTTGTT |
| p199_seq_ScADH2 | ACATTAGAATGGTGATTAGAAAGG |
| p200_seq_ScPCK1 | TCTTTCCCTTGTATAACTTAAAAT |
| p189_seq_CrCPR_1f | ATGATTATGCGGCTGATGATG |
| p190_seq_CrCPR_1r | CGCCAGAATCTGAAGAGTTAG |
| p191_seq_CrCPR_2f | GAAGCAAATGGCCATGCC |
| p192_seq_CrCPR_2r | AACCTTGCAGCATTGTCG |
| p193_seq_CrCPR_3f | ACTAGTTGCAAATCAGAGAAGC |
| p194_seq_CrCPR_3r | ACGATCAGATGCTGGAGTAT |
| p195_seq_CrCPR_4f | CTGCAGTTTTCTTTTTTGGATGC |
| p196_seq_CrCPR_4r | CTGGGTTGGAGGCGTGGA |
| p197_seq_CrCPR_5r | CACGTGAGAAAGCAACAAG |
| p198_seq_CrCYPb5_1r | GCCAGCAATCCTTTGTTTT |
| p294_seq_ScICL1 | AATGGAAACCTGGGGCAAAAG |
| p292_C_cPCR_pICL1 | TATTTGTCTTGGCTTGCTAATTTC |
| p297_seq_CrCYPADH_r1 | TAGCAAACAAAATCTTAATCCTG |
| p298_seq_CrCYPADH_r2 | TCGAGCTCCCTCCACAGC |
| p299_seq_CrCYPADH_f1 | GAAATGTTGTACCAATTCCTTAGC |
| p300_seq_CrCYPADH_f2 | GAACCCAGAAAAGCATGACC |
| P14_A-tADH1-p8379-Rv | GAAATTCGCTTATTTAGAAGTGTC |
| P16_A-tCYC1-p8379-Rv | CTCCTTCCTTTTCGGTTAGAG |
| p306_seq_CrIO1 | TGAACGAAACTACAGATCTAAGA |
| p307_seq_CrIO2 | TACTCTGTCGATCTCTTCCCTTACT |
| p308_seq_CrIO3 | CGTGCGATTTAGATATGAACTCTCT |
| p309_seq_Cr7DLGT1 | GTTTTTGTATGCCTGAACTAATTGA |
| p310_seq_Cr7DLGT2 | AAGAGGGCACAAAAGCCAGA |
| p311_seq_Cr7DLGT3 | ATTCAAATAATCAGTGATTTTATG |
| p312_seq_Cr7DLGT4 | GTCATCATGGAAATGTTTTTTGCAGG |
| p313_seq_Cr7DLH1 | TTTTGACTGGAATTGGAAGCTTAG |
| p314_seq_Cr7DLH2 | TGAAATCAGGAGAAGCAGCAAG |
| p315_seq_Cr7DLH3 | ATGGCGATGTCTCTAATCTTG |
| p349_seq_CrSTR_1 | GAACCCTTTATCAGTTGAATCG |
| p360_seq_CrSTR_2 | CCATCTTTGTGTGGTTGG |
| p361_seq_CrSTR_3 | GTTTGATGGATTTGGGAATATTC |
| P408_seq_CrG8H_1 | TGGTTTGGAATATAATGGTTGAGGC |
| P409_seq_CrG8H_2 | ACTGTTTGGGATGATGCTTTGGC |
| P410_seq_CrGOR_1 | CTGTTCGACATGGTCAGAGTATATAGTG |
| P411_seq_CrGOR_2 | GGAACTGCAGTATTGATTGGGGC |
| P416_seq_CoCa610_1 | ACGTGAATCATCAACCAAAGGTAGTGTCG |
| P417_seq_CoCa610_2 | CTTTTGCATAGAGATAAAGAAATGTGGGGT |
| P418_seq_CoCa565_1 | GATTGATTTATTTCCAACATTTGATACACT |
| P419_seq_CoCa565_2 | AGAATCTGAAAATCGTACCAATGATATTG |
| P472_seq_CrLAMT_1 | TGGTTATCTAAAGTGCCCAAGG |
| P473_seq_CrSLS_1 | AAACAAGGATCCCATGAAATTG |
| P474_seq_CrSLS_2 | GGACACAAAGTTAGGTCCGTACACA |
| p491_cPCR_del_OYE2_U | AAATATCTTACGTAATGAACTTCCG |
| p492_cPCR_del_OYE2_D | ACTCTCAGATTCATATCAACTTCCT |
| p493_cPCR_del_OYE3_U | AAATAAAACAAGGAAGGTAGGGTAA |
| p494_cPCR_del_OYE3_D | AAAAAGAGATACTATAGGCCTAAAA |
| p495_cPCR_del_ATF1_U | TATGTTAGAGTACTTGTACTTGACA |
| p496_cPCR_del_ATF1_D | AACGGTTTTTTTTCAGGGACAAT |
| p497_cPCR_del_ARI1_U | CAGCATATTCGAAAATATATCAACT |
| p498_cPCR_del_ARI1_D | TTGAAACTCTTTTGCAGCCTTAAT |
| p507_cPCR_del_ADH6_U | TTATCTTTTTACAGTAAATGGGTG |
| p508_cPCR_del_ADH6_D | TGCACCTTTTTTGTTAGTGTATTG |
| P565_NcISY2_seq_1 | TACCTCGACTGAACACCAACA |
| p637_seq_Erg20.1 | CACTTCAGAAACGAAAAATA |
| p638_seq_Erg20.2 | TCTAAGTAGTCATCTTGAAT |
| p639_seq_tCrGES.1 | CATATTTGGGAGCCAATGGA |
| p646_seq_tCrGES.2 | CCTAAAGGCAACGTGGATTG |
| p647_seq_tHMGR.1 | TCTTATCATATACCAATGGC |
| p648_seq_tHMGR.2 | ACATGCAGCTAATTTAGTGA |
| p649_seq_MLS1.1 | TTATGCTATAGTACCTAAGA |
| p650_seq_MLS1.2 | CAATACAAAATTTATCCGAA |
| p655_Erg20p_ver_1 | GACAATCATTACCACAAGATGAACA |
| p659_seq_IDI1_1 | GAATTAGGTTTGAAGGGTAA |
| p660_seq_Erg20_switch1 | ACGTTCTGATAAATAACCAC |

| | |
|---|---|
| p661_seq_Erg20_switch2 | CCAAGGAATTAATCGATATC |
| p662_seq_Erg20_switch3 | TCTGGGGTCTCATGTTCTTT |
| P744_Erg20promoter_seq | CTTCACACTTTTAACTGGACGTT |
| p749_seq_CrTDC_1 | GTATTGGAAAACTTGTCTGTTACG |
| p750_seq_CrTDC_2 | TTACTAAGGGCACTCACTACGA |
| p751_seq_ScZWF1_1 | GTAATCGTAGAGAAACCTTTCG |
| p752_seq_ScZWF1_2 | ATGACTTTCAACATCGAAAACGAG |
| P753_seq_Erg20_switch4 | GAGAATTCTTCATGTAATTTACCA |
| P754_seq_Erg20_switch5 | TCTGGGGTCTCATGTTCTTT |
| p979_seqBB4 | CAAGCAAGGTTTTCAGTATAATG |
| p1439_cPCR_ADH3_fw | CTTTTCGCCAGCTCCTAAAC |
| p1440_cPCR_ADH3_rv | CTCGATGCTTGATGGTGATAATG |
| p1441_cPCR_ALD6_fw | CCTGGCGTGTTTAACAAGTTC |
| p1442_cPCR_ALD6_rv | GCAGTTGTTGTACACTAGCTTAAG |
| p1443_cPCR_YPL062W_fw | TCAATTCGTACAATGCCTGGCAT |
| p1444_cPCR_YPL062W_rv | GACTGAACACTTCGAATTGAATATTACC |

### Supplementary Table 8. List of primers used for mutagenesis in this study.

| Primers for mutagenesis | | Comment |
|---|---|---|
| p631_Infu_ERG20.A_f | CACTATAGGGAATATTAAGCTATGGCTTCAGAAAAAGAAATTAGGAGAG | fw/rv primer generating fragment for Infusion Cloning of Erg20$^{WW}$ |
| p632_Infu_ERG20.A_r | GTAAGCCTGCAACAACTCAATGCACCAACCTAGAATGGCAACC | |
| p633_Infu_ERG20.B_f | TTGAGTTGTTGCAGGCTTACTGGTTGGTCGCCGATGATATGATGGACAAGTCCATTACCAGAAGAGGCCAACCATG | fw/rv primer generating fragment for Infusion Cloning of Erg20$^{WW}$ |
| p634_Infu_ERG20.B_r | GCCTCTAACATGAATGCGTCCCAGATGGCAATTTCCCCAACTTCAGGAACCTTGTACCAACATGGTTGGCCTCTTC | |
| p635_Infu_ERG20.C_f | GACGCATTCATGTTAGAGGCTGCTATCTACAAGCTTTTGAAATCTC | fw/rv primer generating fragment for Infusion Cloning of Erg20$^{WW}$ |
| p636_Infu_ERG20.C_r | CATGATGCGGCCCTCTAGCTATTTGCTTCTCTTGTAAACTTTGTTC | |
| p1085_Erg20WWG_INFU_fw | TTACTTTCGGTACTGCTTACTATTCTTTCTACTTG | fw/rv primer generating fragment for Infusion Cloning of Erg20$^{WWG}$ |
| p1086_Erg20WWG_INFU_rv | GTAAGCAGTACCGAAAGTAACTATGAAGGAGTG | |

**Supplementary Table 9. List of custom gRNA sequences for gene deletions used in this study.**

| Target gene to be deleted | gRNA sequence |
|---|---|
| ScATF1 | CAGGCATTCTTGTTGCACGGNGG |
| ScARI1 | TGCGCCAGAAACGAAAACAGNGG |
| ScOYE2 | GTAACCCCCAGATTGTGGAGNGG |
| ScOYE3 | CTCATCAGACCAAATCCCAGNGG |
| ScADH6 | GACATTAAGATCGAAGCATGNGG |
| CrGES | AATTGTCTGAGATAACAAGG |
| ScADH3 | CAAGCCGCCAAAATTCAACA |
| ScALD6 | AACGGTAGAACAATCAACAC |
| ScYPL062W | CATATGCAACAATGACGTCT |
| CrLAMT | GAAGCCCACCCAATGAAAGG |
| ScPHXT1 | CCGATAAATAGAGGAAGCAA |
| Cr7DLH | TGTCCACACGAGTAAAGTCG |

**Supplementary Table 10. List of metabolites used in this study.**

| Metabolite | CAS # | Source | Reference / Catalogue # |
|---|---|---|---|
| Geraniol (**11**) | 106-24-1 | Purchased from Sigma-Aldrich | 163333-25G |
| 8-Hydroxygeraniol (**12**) | 26488-97-1 | Synthesised | See Supplementary methods |
| 8-Oxogeraniol (**21**) | 38290-51-6 | Synthesised | See Supplementary methods |
| 8-Oxogeranial (**13**) | 80054-40-6 | Synthesised | See Supplementary methods |
| 7-Deoxyloganic acid (**16**) | 22487-36-1 | Prepared by yeast biotransformation | See Supplementary methods |
| Loganic acid (**17**) | 22255-40-9 | Purchased from Biomol | Cay28402-5 |
| Loganin (**18**) | 18524-94-2 | Purchased from Biomol | Cay19997-5 |
| Secologanin (**19**) (authentic reference) | 19351-63-4 | Purchased from Biomol | Cay20647-5 |
| Secologanin (**19**) | 19351-63-4 | Isolated from *Lonicera tatarica* | As reported by (Zhu et al., 2021) |
| Secologanic acid (**22**) (authentic reference) | 60077-46-5 | Purchased from Biozol | CAS [60077-46-5] |
| Secologanic acid (**22**) | 60077-46-5 | Isolated from *Lonicera tatarica* | As reported by (Zhu et al., 2021) |
| Secoxyloganic acid (**23**) | 59472-23-0 | Purchased from Biopurify | BP2107 |
| Strictosidine (**3**) | 20824-29-7 | Prepared by yeast biotransformation | See Supplementary methods |
| Strictosidinic acid (**4**) | 150148-81-5 | Prepared by yeast biotransformation | See Supplementary methods |
| (2E,6E)-2,6-dimethylocta-2,6-dienedioic acid (DOA) (**6**) | | Isolated from yeast culture | See Methods section 2.6 |

**Supplementary Table 11. NMR assignments of (2*E*,6*E*)-2,6-dimethylocta-2,6-dienedioic acid (DOA) (6).**

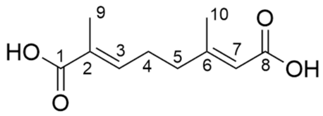


| Atom | $^{13}$C ppm | $^{1}$H ppm (m, Hz) |
|---|---|---|
| 1 | 171.76 | - |
| 2 | 130.06 | - |
| 3 | 141.81 | 6.77 – 6.69 (m, 1H) |
| 4 | 27.63 | 2.41 (q, $J$ = 7.3 Hz, 2H) |
| 5 | 40.35 | 2.31 (t, $J$ = 7.5 Hz, 2H) |
| 6 | 159.44 | - |
| 7 | 117.77 | 5.69 (s, 1H) |
| 8 | 170.42 | - |
| 9 | 12.54 | 1.83 (s, 3H) |
| 10 | 18.75 | 2.15 (d, $J$ = 1.2 Hz, 3H) |
